# Identification of RNA virus-derived RdRp sequences in publicly available transcriptomic datasets

**DOI:** 10.1101/2022.10.18.512700

**Authors:** Ingrida Olendraite, Katherine Brown, Andrew E. Firth

## Abstract

RNA viruses are abundant, highly diverse, and infect all or most eukaryotic organisms. However, only a tiny fraction of the number and diversity of RNA virus species have been catalogued. To cost effectively expand the diversity of known RNA virus sequences we mined publicly available transcriptomic datasets. We developed 77 family-level Hidden Markov Model profiles for the viral RNA dependent RNA polymerase – the only universal “hall-mark” gene of RNA viruses. By using these to search the NCBI Transcriptome Shotgun Assembly database, we identified 5,867 contigs encoding RNA virus RdRps or fragments thereof and analysed their diversity, taxonomic classification, phylogeny and host associations. Our study expands the known diversity of RNA viruses, and the 77 curated RdRp pHMMs provide a useful resource for the virus discovery community.

## Introduction

RNA viruses evolve rapidly and are extremely diverse, with an ancient evolutionary ancestry (Koonin et al., 2015). They are highly abundant in eukaryotic hosts, whereas the viromes of prokaryotes tend to be dominated by DNA viruses (Koonin et al., 2015, though cf. Callanan et al., 2020 and Neri et al., 2022). Genomes range in size from ∼2 kb to ∼41 kb (Nibert, 2017; Saberi et al., 2018). Over long evolutionary timescales, RNA viruses evolve via extensive recombination of protein-coding genes between different lineages, between virus and host, and via *de novo* gene formation (Keese & Gibbs, 1992; Shi et al., 2016; Dolja & Koonin, 2018; Wolf et al., 2018). The unique hallmark gene of RNA viruses – and indeed the only protein that is ubiquitously conserved throughout all RNA viruses – is the RNA dependent RNA polymerase (RdRp). Despite immense divergence and, in some cases, almost imperceptible similarity at the primary sequence level, structural studies have confirmed homology – and therefore shared evolutionary ancestry – between the RdRps of all three Baltimore RNA virus groups (Mönttinen et al., 2021; Jácome et al., 2022). Besides being the only hallmark protein of RNA viruses, the RdRp is also relatively highly conserved compared to other RNA virus proteins, and is therefore the most appropriate protein by which to identify new RNA viruses with homology search algorithms such as BLASTP (Altschul et al., 1990) or HMMER (Eddy, 2011).

In the traditional Baltimore Classification system, RNA viruses are classified into three groups based on the nature of their genomic nucleic acid: single-stranded positive-sense RNA (+ssRNA) viruses, double-stranded RNA (dsRNA) viruses, and single-stranded negative-sense RNA (−ssRNA) viruses (Baltimore, 1971). “Positive-sense” refers to the coding sense, although a small number of RNA viruses have also evolved additional genes on the opposite strand (Nguyen & Haenni, 2003; Dinan et al., 2020). During replication in a host cell, all RNA viruses must obviously produce the complementary RNA. Thus, the virus genome is defined to be the nucleic acid that is packaged into virus particles for extracellular transmission. A small proportion of viruses lack capsids, and here the appropriate Baltimore group is determined by phylogenetic relatedness to viruses with capsids. Depending on species, RNA viruses can have segmented or non-segmented genomes that are almost always linear, but in a few cases (notably chuviruses and the recently discovered ambiviruses) are or appear to be circular (Li et al., 2015; Forgia et al., 2022b; Lee et al., 2022).

Deep phylogenetic analysis of RNA virus RdRps has outlined five putative major branches of RNA viruses (Wolf et al., 2018; Koonin et al., 2020). The basal Branch 1 comprises +ssRNA narna-, mito- and ourmiaviruses and their bacterial levivirus relatives. Branch 2 comprises the vast picornavirus-like supergroup including +ssRNA picorna-, calici-, polero-, nido-, astro- and potyviruses, and the dsRNA partiti-, picobirna-, amalga- and hypoviruses. Branch 3 includes the +ssRNA alpha-, flavi-, tombus-, noda-, yan-, zhao- and weiviruses and their relatives. Branch 4 comprises the dsRNA reo-, toti-, chryso-, megabirna-, quadri- and giardiaviruses, besides their bacterial cystovirus relatives. Finally, Branch 5 – predicted to have evolved out of Branch 4 – comprises all known −ssRNA viruses, including bunya-, orthomyxo-, mononega-, chu-, qin- and yueviruses. These branches were subsequently designated as phyla, with Branches 1 to 5 corresponding to phyla *Lenarviricota*, *Pisuviricota*, *Kitrinoviricota*, *Duplornaviricota* and *Negarnaviricota*, respectively. Some recent studies have indicated the existance of additional phylum-level groups (Neri et al., 2022; Zayed et al., 2022), though, beyond sequence data, very little is known about the members of these new clades. Virus taxonomy is supervised by the International Committee on Taxonomy of Viruses (ICTV) and, over the past several years, has been in a state of flux as the system adapts to the “metagenomic era”, adopts additional levels of classification, and moves to binomial genus/species names to increase consistency with the Latin binomials used for cellular organisms. In this study we used the ICTV 2017 taxonomy (except when referring to more recent literature) (King et al., 2018).

The ∼5,500 RNA virus (kingdom *Orthornavirae*) species currently represented in GenBank constitute just a tiny fraction of the estimated millions of RNA virus species on Earth (Geoghegan & Holmes, 2017; Kuhn et al., 2019; Dance, 2021; Harvey & Holmes, 2022). Recent high-throughput RNA sequencing (RNA-Seq) studies – performed with the express purpose of identifying RNA viruses – have revealed vast numbers of novel RNA viruses, and many new family-level virus clades in diverse eukaryotic host organisms (Cook et al., 2013; Li et al., 2015; Shi et al., 2016, 2018; Olendraite et al., 2017; Charon et al., 2020; Chiapello et al., 2020; Sutela et al., 2020; Wolf et al., 2020; Batson et al., 2021; Chen et al., 2022; Forgia et al., 2022a; Kinsella et al., 2022; Rosario et al., 2022; reviewed in Dolja & Koonin, 2018; Greninger, 2018; Obbard, 2018; Zhang et al., 2019; Cobbin et al., 2021; Harvey & Holmes, 2022). However, a much larger number of RNA-Seq studies are performed for projects that are unrelated to virus discovery, but instead aim to study the transcriptomes of the targeted cellular organisms. The results of these studies are often deposited in the National Center for Biotechnology Information (NCBI) Short Read Archive (SRA; raw sequencing reads) and Transcriptome Shotgun Assembly (TSA; assembled contigs) databases (Sayers et al., 2022). It should be noted that only a small fraction of SRA datasets have currently been assembled into TSA datasets. Nonetheless the TSA database still holds assemblies for thousands of RNA-Seq datasets from diverse host organisms. By analysing public RNA-Seq data in the TSA database, we aimed to expand the known diversity of RNA viruses and their host species.

Profile Hidden Markov Models (pHMMs) provide a more sensitive method than BLASTP for finding distant homologues of known protein sequences while (in contrast to structure-based approaches) maintaining computational speed (Eddy, 2011). In this analysis, we used known RNA virus sequences to develop 77 family-level RdRp pHMMs. We used these to search with HMMER (Eddy, 2011) for sequences encoding putative RNA virus RdRps in the TSA database, supplemented with virus sequences from the NCBI RefSeq (ref) and non-redundant nucleotide (nr/nt) databases. We identified 12,109 RdRp-encoding sequences and analysed their diversity, taxonomic classification, phylogeny, and host associations. We provide a listing of all sequences found and our curated family level pHMMs, both of which will be useful resources for evolutionary, taxonomic, host association, ecological and comparative genomic studies.

## Results

### Enriching viral RdRp diversity

Profile Hidden Markov Models (pHMMs) provide a fast and sensitive method for identifying distant homologues of protein sequences (Eddy, 2011). We began by constructing pHMMs for the RNA dependent RNA polymerase (RdRp) proteins of 77 RNA virus family-level clades (see Methods). We used these pHMMs together with HMMER (Eddy, 2011) to search for candidate RdRp-encoding sequences in the TSA database, supplemented with virus sequences from the NCBI RefSeq (ref) and non-redundant nucleotide (nr/nt) databases. Since some RNA viruses are segmented, a caveat with this strategy is that it is not always possible to retrieve entire virus genomes.

We first discarded nr/nt and ref sequences that had ≥80% nucleotide identity to longer sequences. Next, for the TSA sequences and remaining nr/nt and ref sequences, we extracted all ORFs of length ≥60 nucleotides, and applied HMMER to each ORF (see Methods). 15,044 putative virus RdRp sequence fragments with a significant match (p-value ≤10^−6^) against at least one pHMM were identified, corresponding to 12,136 translated ORFs in 12,109 unique accessions (Supplementary Dataset 1). 5,867 accessions were from the TSA database. These sequences were sorted into “classified”, “ambiguously classified” or “unclassified” groups, based on the HMMER bit score divided by the length in amino acids of the alignment between an ORF and a matched pHMM (referred to herein as “IDscore”). In brief, if the highest IDscore for a sequence:profile match was lower than 0.25, a sequence was sorted into the unclassified group. Otherwise, if a sequence had statistically significant hits to more than one of the pHMMs and the top two IDscores were within 20% of each other, the sequence was sorted into the ambiguously classified group. Finally, sequences with an IDscore of 0.25 or higher, and at least 20% difference in IDscore between the first and second best hits, were sorted into the classified group and classified according to the pHMM of the top IDscore. 55% of sequences were sorted into the classified group, 43% into the ambiguously classified group and 2% into the unclassified group (Figure 1).

**Figure 1.**
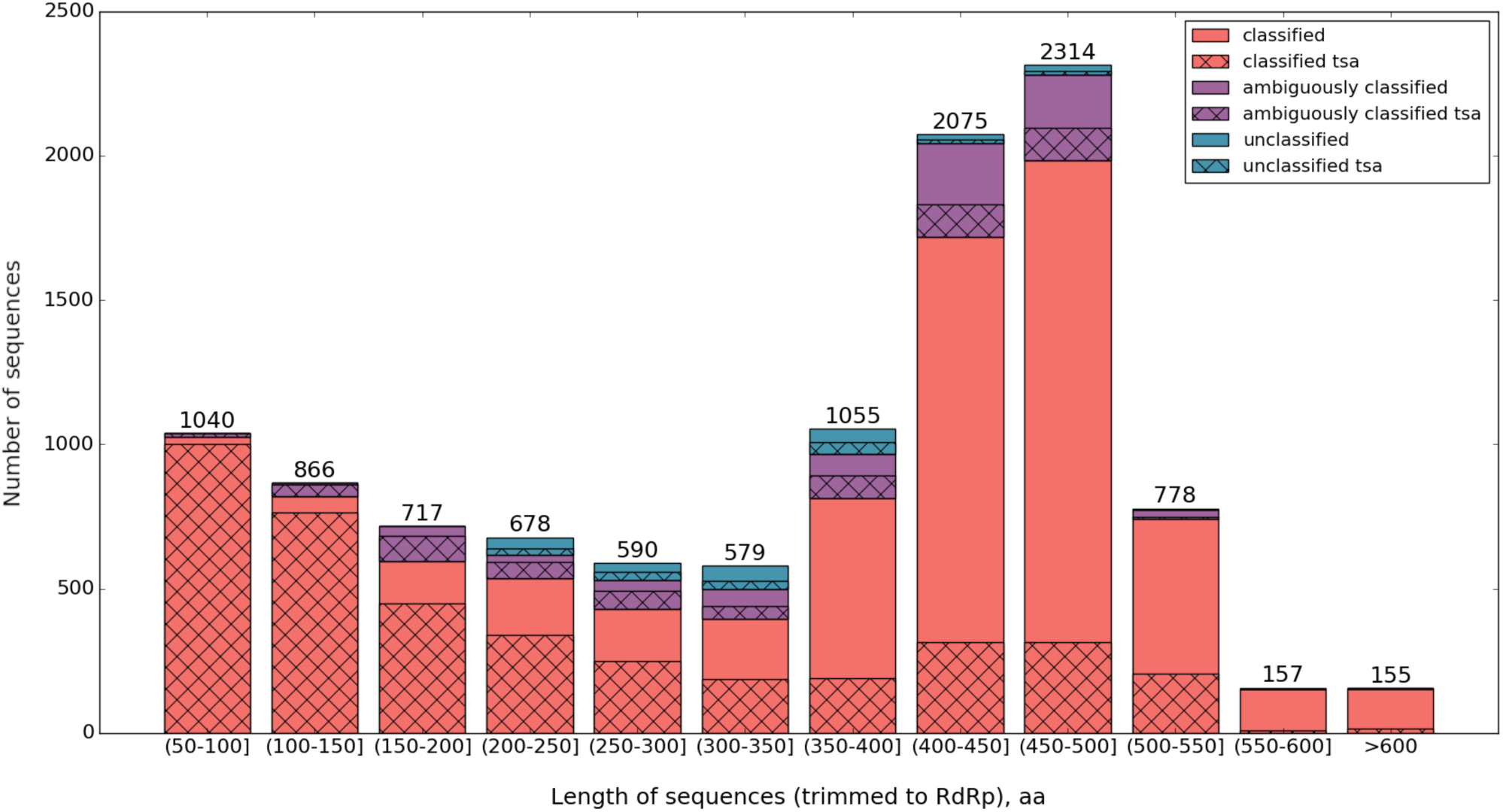
Numbers of identified RdRp-encoding ORFs (ref, nr/nt and TSA) and their lengths after trimming to the RdRp core (see main text) and removing duplicate identical sequences.

In most cases, after trimming translated ORF sequences to the RdRp core (i.e. the best pHMM match positions), the identified RdRp sequences were between 400 and 500 amino acids (a.a.) in length (Figure 1). Typically, actual full-length viral RdRp core sequences range approximately from 350 to 600 a.a. in length (depending on virus clade). Most of the shorter sequences found derived from the TSA database. This was expected, as transcriptome shotgun assembly often results in fragmented rather than full-length sequences (Bushmanova et al., 2019). Thus many of the RdRp sequences found – but especially the shorter ones – are expected to be incomplete.

We verified that putative RNA viral ORFs were more similar to a known RNA virus than to any non-RNA-viral sequence by using a low stringency BLASTp search (Altschul et al., 1990) against the non-redundant protein (nr) database (Supplementary Dataset 2). All sequences showed significant similarity (E-value ≤ 0.05) to a known RNA virus sequence. 25 had better scores (E-value ≤ 0.05) against another sequence not derived from an RNA virus. *In silico* assembly can sometimes lead to artefactual virus-virus or virus-host chimeric sequences (reanalyzing the TSA source data is possible, but was not attempted in this study). It is also possible that some TSA sequences derive from transcribed genome-integrated virus fragments (known as endogenized virus elements or EVEs;Katzourakis & Gifford, 2010; Aiewsakun & Katzourakis, 2015; Gilbert & Belliardo, 2022). Either of these scenarios could result in sequences with both RNA viral and non-RNA-viral regions, in this work we do not distinguish viruses from transcribed EVEs.

We also used HHSearch (Steinegger et al., 2019) against the Pfam database (Finn et al., 2014) to confirm the presence of RNA viral domains in our putative viral ORFs (Supplementary Dataset 2). 98.9% of sequences contained recognisable viral domains (including RdRp in 98.4% of cases). The sequences not identified using this method were primarily (250 of 281 cases) members of the orders *Tymovirales*, *Durnavirales* and *Patatavirales*, all of which form parts of phylum-level Pfam pHMM models, which may be less sensitive than other order- or class-level models.

Whereas the ambiguously classified and unclassified groups contain many novel and divergent viral sequences (discussed further below), sequences in the classified group share substantial similarity with previously known sequences from currently existing virus families. Nonetheless, we wanted to investigate the extent to which the addition of the TSA-derived classified RdRp sequences increases the diversity of RdRps within currently defined RNA virus groups. For each of the 77 (mostly family level) pHMMs, we took all the TSA, ref and nr/nt RdRp sequences that were classified to that group and, using BLASTP, compared all versus all pairwise identities within each group to find the greatest pairwise divergences by group (Supplementary Figure 1). For several of the pHMM groups (e.g. roni-, fimo-, yue-, xinmo-, arto-, deltaflexi-, nyami- and chu-like viruses), the diversity was substantially increased with the addition of the TSA-derived sequences. In some such cases, this may indicate the presence of novel sister-clades (or sister families) within the group. Inspired by Wolf et al. (2020) we also plotted the percentage increase in number of RdRp clusters, as a function of trimmed RdRp core fragment length and clustering identity threshold, upon adding the TSA-derived sequences to the nr/nt sequences. For a length cut-off threshold of 400 a.a., we saw about a 30% increase in the number of RdRp clusters at CDHIT identity thresholds of 70% or 50% (Figure 2).

**Figure 2.**
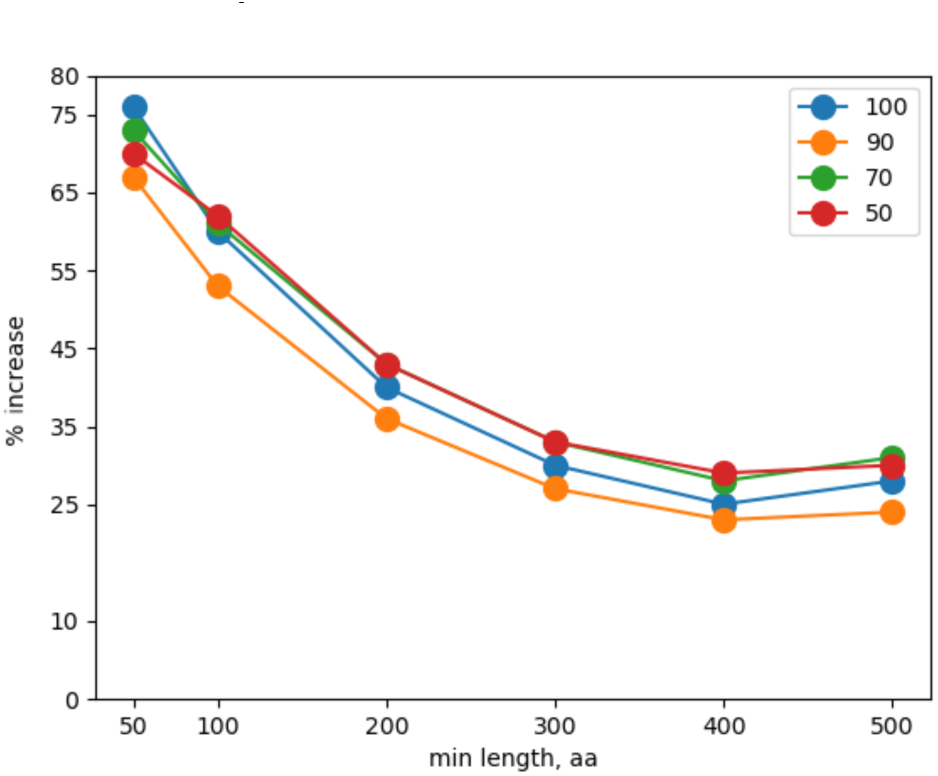
Percentage increase in the number of RdRp clusters as a function of trimmed RdRp core fragment length (x-axis) and clustering identity threshold, upon adding the TSA-derived sequences to the nr/nt and ref sequences. The y-axis shows the percentage increase in clusters after using different CD-HIT (Li & Godzik, 2006; Fu et al., 2012) identity thresholds (50%, 70%, 90% and 100%, as indicated) for nr/nt + TSA sequences compared to nr/nt sequences alone.

After the sorting step, some RdRp pHMM groups had hundreds of sequences whereas others had very few. For simplicity of presentation, we appended the groups containing very few sequences to larger RdRp groups, based (to a certain extent) on their RdRp similarities and the size of the larger groups. This resulted in 60 clusters, with sizes and amino acid sequence diversity as illustrated in Figure 3. PhyML trees of the different virus groups are available in Supplementary Dataset 3.

**Figure 3.**
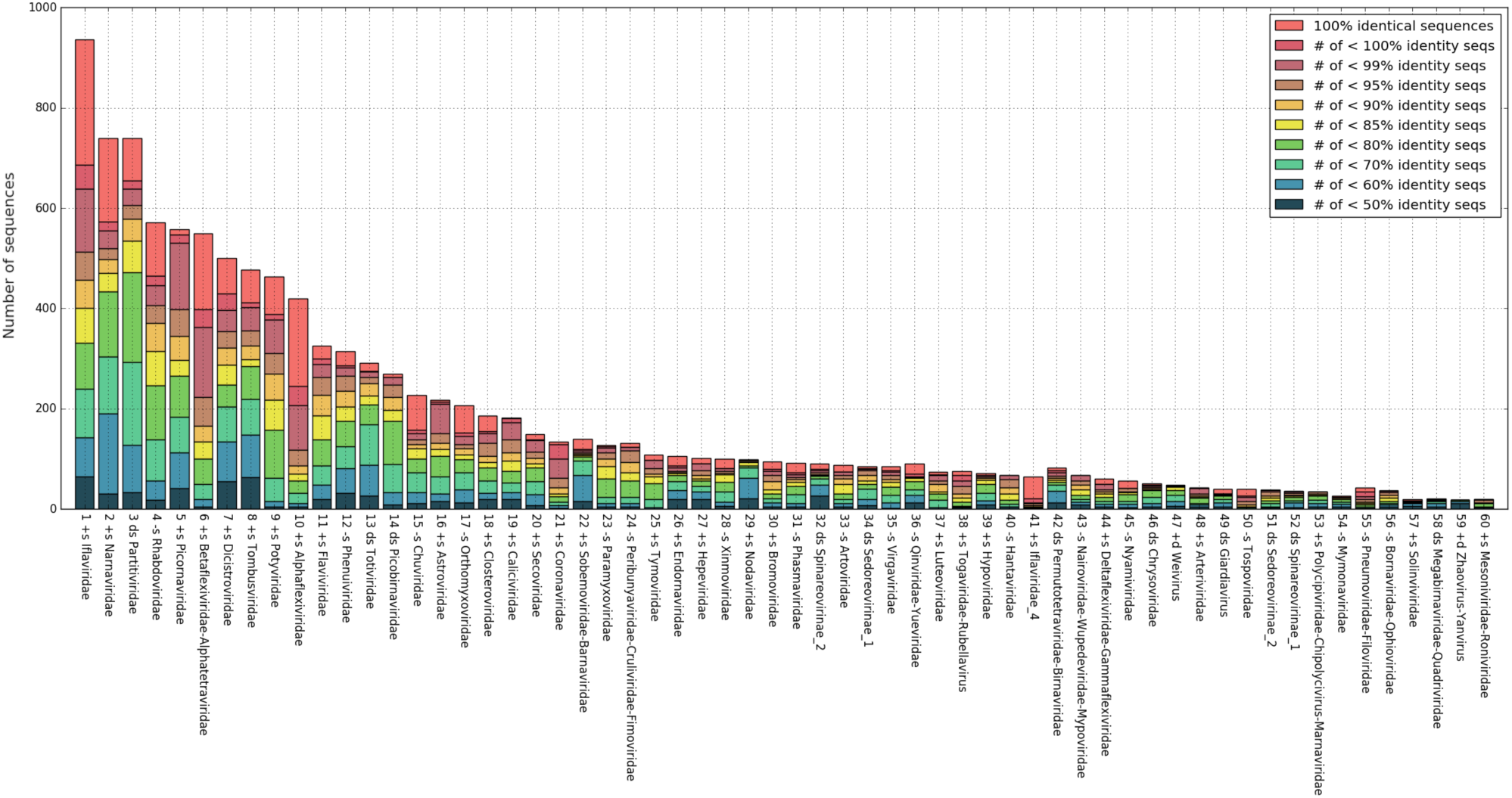
Numbers of sequences identified in each cluster at different pairwise amino acid identity thresholds. Duplicate identical sequences were removed. Identities were calculated via pairwise alignment in Biopython (Cock et al., 2009, see Methods) and dividing the number of identical aligned residues by the shorter sequence length.

We also compared the number of RdRp sequences found per group in the TSA and in the nr/nt databases (Figure 4). Our TSA search revealed large numbers (>100 per group) of new ifla-, narna-, partiti-, betaflexi-, dicistro-, alphaflexi-, rhabdo-, tombus-, chu-, poty-, phenui-, orthomyxo-, toti- and picobirna-like RdRps. New TSA sequences also represented >75% of some groups such as the ifla_4-, arto-, xinmo-, deltaflexi-, ifla-, qin-, giardia-, nyami-, yue-, chu-, alphaflexi-, phasma- and narna-like viruses (note, ifla-4 denotes a partition of the iflaviruses in the Aiewsakun and Simmonds (2018) groupings that we used for HMM production). Many of these virus groups have previously been found to be associated with arthropods, fungi, plants or protists. On the other hand, for virus groups that previously have been found to be mainly or strictly vertebrate-associated, such as the picorna-, astro-, calici-, corona-, paramyxo-, hanta- and arteri-like viruses, we found relatively few new TSA sequences, and the TSA sequences we found comprised <7% (average 2.4%) of the total number of sequences found for each group.

**Figure 4.**
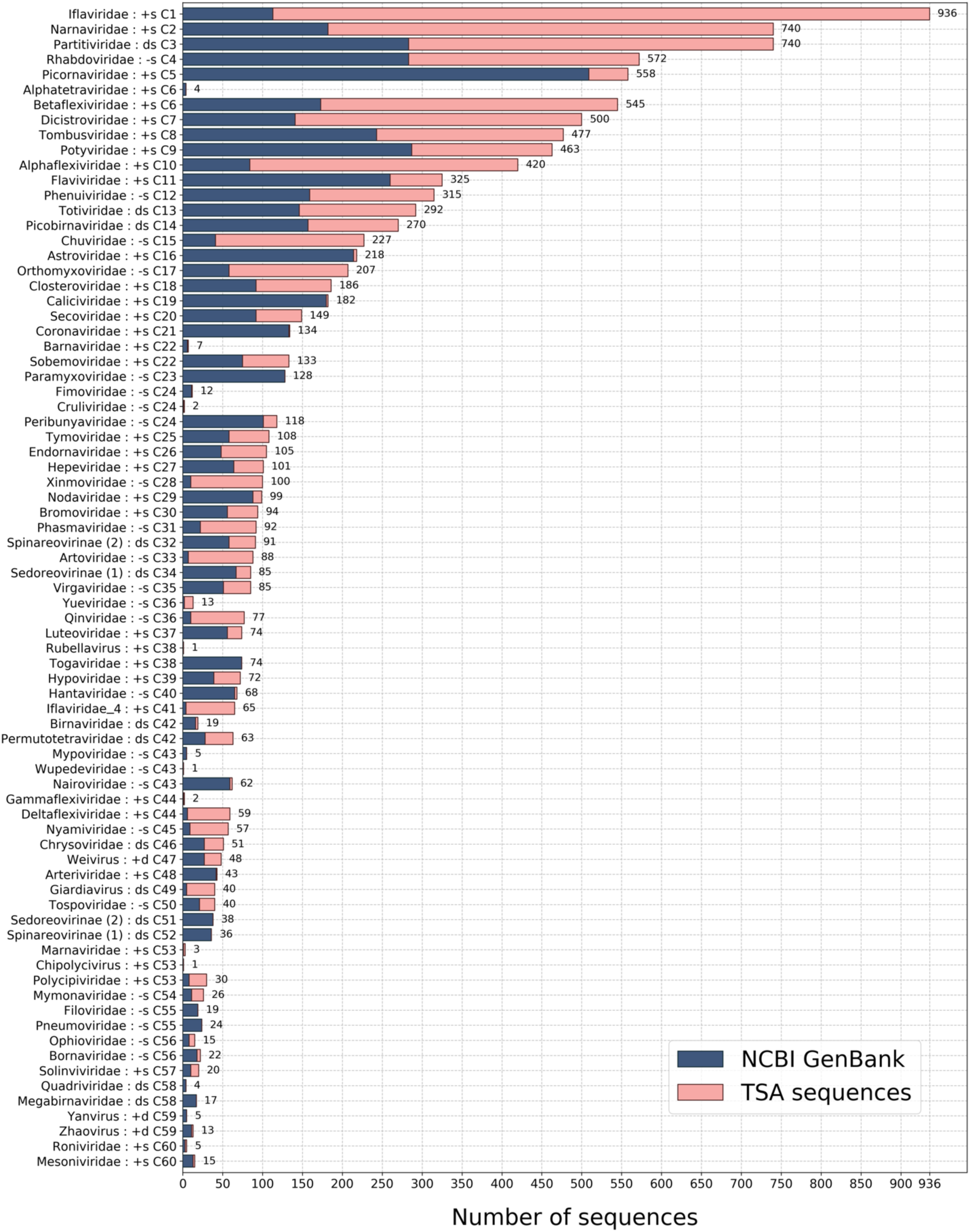
Total number of classified sequences in each group of classified sequences (cluster numbers C1 to C60). Blue – nr/nt and ref sequences; pink – TSA sequences; +s –+ssRNA; −s – −ssRNA; ds – dsRNA.

For several virus groups, our analysis revealed large numbers of TSA-derived sequences. For example, 227 (186 TSA and 41 nr/nt) RdRp sequences were classified as best matching the chu-like virus pHMM, from which we obtained 158 non-identical trimmed sequences for phylogenetic analysis (Supplementary Figure 2). Some of the sequences comprise fragments that, in some cases, might derive from the same genome and appear in slightly different places on the tree depending on which region of the RdRp core each fragment covers. A likely full-length RdRp core was present in 85 (45 TSA and 40 nr/nt or ref) of the 158 sequences. There were 14 TSA-derived sequences (from 14 different invertebrate NCBI BioProjects) with contig length >10,000 nt that likely correspond to substantially complete representatives of the non-segmented form of the chu-like virus genome.

### Host associations of TSA-derived RdRp sequences

Next we looked at which types of cellular life forms the classified TSA-derived sequences putatively infect. For this, we extracted the target organism species name from the metadata accompanying each TSA dataset and downloaded the corresponding taxonomic information (family, order, class, phylum, etc) from NCBI. We then grouped TSA datasets into the following categories: vertebrates (subphylum Vertebrata), arthropods (phylum Arthropoda), invertebrates (kingdom Metazoa/Animalia excluding Arthropoda and Vertebrata), plants (unranked clade Viridiplantae), fungi (kingdom Fungi), protists (domain Eukaryota excluding Metazoa, Fungi and Viridiplantae) and metagenomic samples (datasets annotated as metagenomic, environmental or bacterial; though note that the eight bacterial datasets did not produce any matches to our RdRp pHMMs). Next, we calculated the number of different RdRps, number of TSA datasets, and number of distinct host species represented within each host category (Figure 5; Supplementary Table 1). We identified relatively few RdRps in vertebrates (1.7% of total number), despite vertebrates representing almost 20% of the TSA datasets. Meanwhile, 48.6% of all identified RdRps were found in arthropods which represent 34.7% of the TSA datasets, equating to ∼2.2 RdRps per dataset on average (1,976 RdRps / 918 datasets). A similar ratio was observed for plants (1,440 RdRps / 650 datasets). Not surprisingly, the highest ratio, ∼6.6 RdRps per dataset, was observed for metagenomic samples. This could be due to more fragmented sequences in metagenomic samples and/or the multi-species nature of such samples.

**Figure 5.**
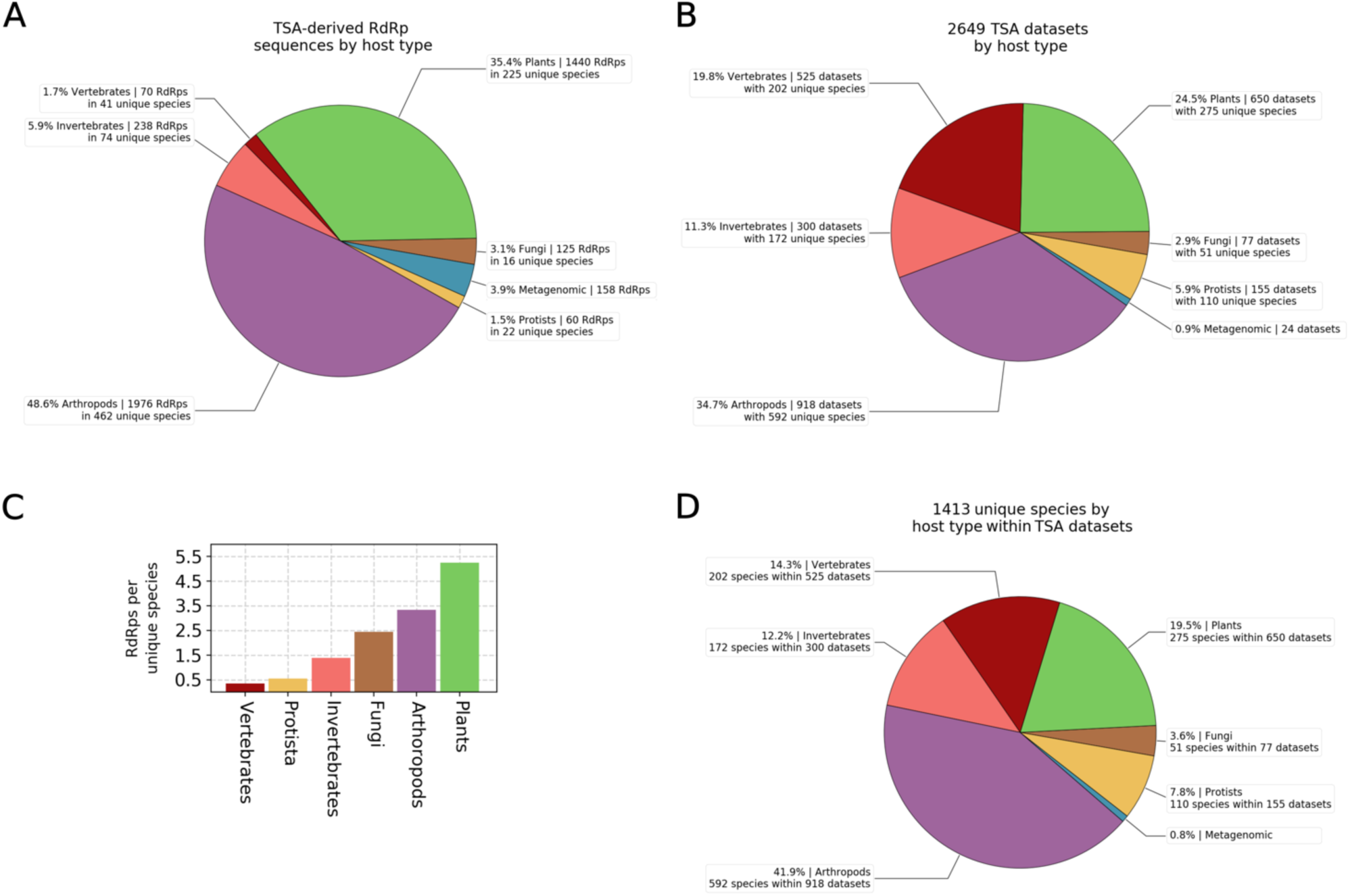
Distributions of RdRps and host species across TSA datasets. Only non-identical RdRp core sequences were used (i.e. discarding duplicate 100%-identical RdRp sequences within each classified-pHMM group, including any identical to nr/nt or ref sequences, leaving the longest representative). No RdRps were detected in the 8 bacteria TSA datasets with our pHMMs. **(A)** RdRp counts per host type. **(B)** TSA dataset counts per host type. **(C)** Mean number of RdRps per host species. Note that within metagenomics samples, the majority of ‘species’ were named “gut metagenome”. **(D)** Numbers of unique TSA dataset host species, grouped by host type.

We also checked which individual host species provided the most non-identical RdRp core sequences (combined over multiple TSA datasets and excluding metagenome datasets). The plants *Saccharum* hybrid (with 78 RdRp sequences), *Tinospora cordifolia* (74) and *Agave tequilana* (53), followed by the insect *Bemisia tabaci* (38), provided the most RdRp sequences. Many other plant and invertebrate species also provided >10 RdRp sequences per species (Supplementary Figure 3). Factors contributing to RdRp richness may include number of independent samples for a host species, sequencing depth, pooling strategy (e.g. pooling RNA from multiple individual organisms into one sample), and likelihood of contamination (e.g. fungi on plant leaves, gut contents in whole-insect samples, etc). In any case, from these numbers it is clear that there remains an enormous amount of unsampled RNA virus diversity in plants and arthropods.

We also calculated the number of unique host species for each classified pHMM cluster (Figure 6). In five cases – ifla-, partiti-, narna-, dicistro- and rhabdo-like viruses – the identified RdRps derived from >100 putative host species. Meanwhile, for picornaviridae-like viruses, despite this family having numerous NCBI sequences, TSA sequences were found only in a few datasets with the majority being from vertebrate samples. As expected, groups such as ifla-, chu-, xinmo-, phasma-, arto- and solinvi-like RdRps were largely arthropod-associated. This is consistent with the diversity of arthropod viruses observed in previous studies, such as Shi et al. (2016), Batson et al. (2021) and Chang et al. (2021). Groups such as alphaflexi-, poty-, betaflexi- and tospo-like RdRps were largely plant-associated, again as observed previously, for example by Mifsud et al. (2022).

**Figure 6.**
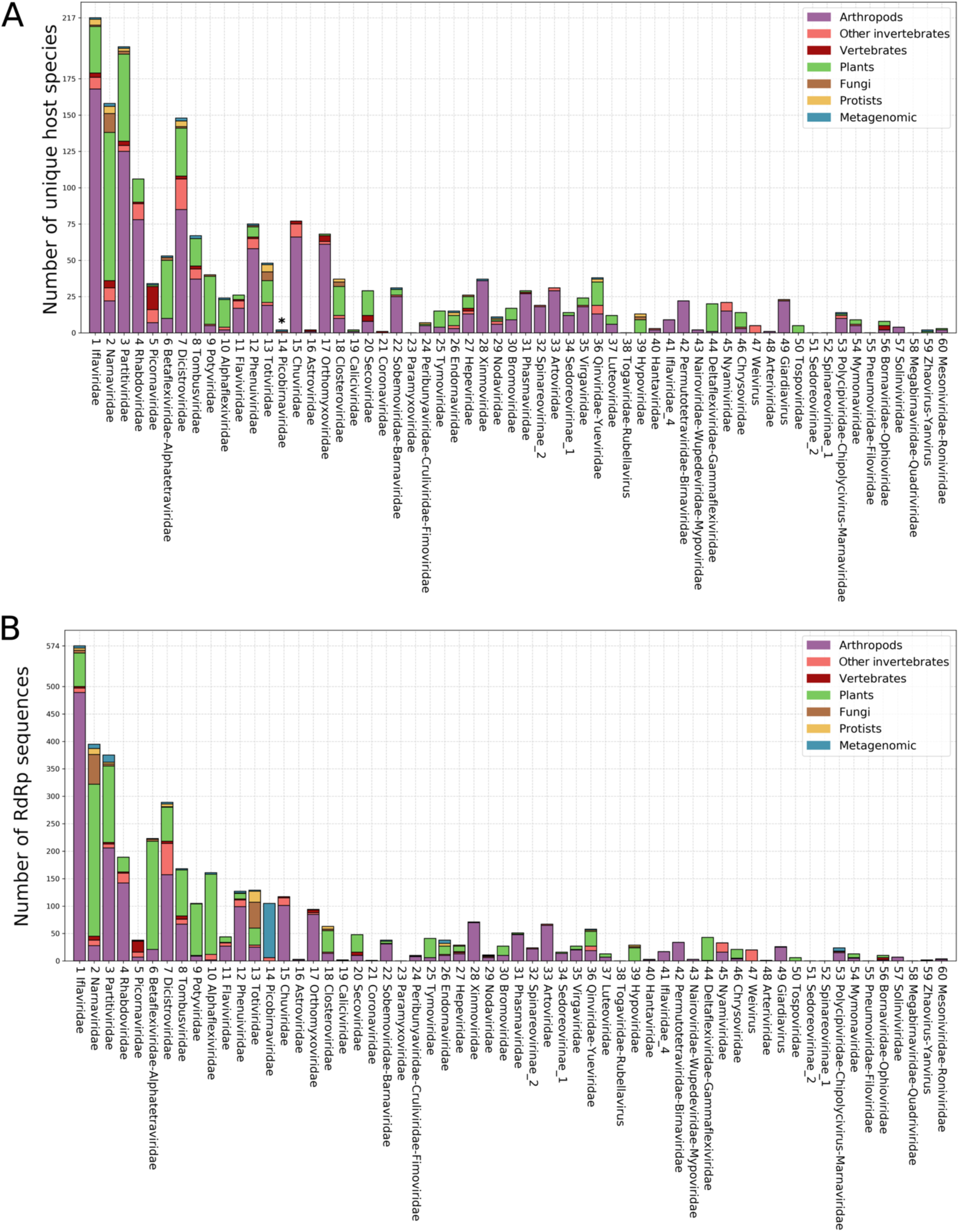
Numbers of unique putative host species **(A)** and numbers of TSA-derived RdRps **(B)** for different classified-pHMM clusters, separated by host species category as indicated in the key. Duplicate 100%-identical RdRp core sequences were removed (as in Figure 5). Asterisk - note that the majority of the metagenomic datasets are labelled as ‘gut metagenome’ which is here counted as a single “species” name.

However, there were also unexpected associations. For example, classical dicistroviruses are an arthropod-associated group, yet around 45% of the dicistro-like RdRps derived from non-arthropod TSA datasets. Many of these dicistro-like RdRps originated from non-arthropod invertebrate (∼20%) and plant (∼22%) datasets (Figure 6A). In some cases, this might result from contamination (e.g. arthropods accidentally sequenced along with plant leaves). Mifsud et al. (2022) quantified contamination in plant TSA datasets and found arthropod contamination in many libraries. Plant-grazing arthropods can also transiently introduce their viruses into plants (Gildow & D’Arcy, 1988; Wamonje et al., 2017). In addition, some dicistro-like sequences found in datasets labelled as vertebrate likely derive from arthropod viruses present in vertebrate faecal samples.

Interestingly, the majority (>60%) of classified metagenomic-derived RdRps were placed within the picobirna-like group. It has been suggested that picobirnaviruses may in fact be RNA bacteriophages rather than viruses of eukaryotes (Krishnamurthy & Wang, 2018; Wang, 2022), which may explain why they are largely absent from our eukaryote-derived TSA datasets but abundant in the metagenomic datasets which typically comprise gut metagenomes (Figure 6B). This is also consistent with Chen et al. (2022) where the virome of metagenomic faecal samples was found to contain many picobirna-like viruses.

### Evolution of motif C of the RdRp

RNA virus RdRps contain a number of highly conserved motifs, labelled A to G (or I to VIII) (Koonin, 1991; Koonin & Dolja, 1993; Bruenn, 2003; te Velthuis, 2014). Of these, motif C (or motif VI) is the most distinctive and most highly conserved. We decided to leverage our collection of diverse RdRps to more fully understand the extent of variation within the core triplet (typically GDD) of motif C among classified viruses. We took all classified sequences within each pHMM group, aligned them, and manually located the conserved motif C. Consistent with previous work, we saw six possible variations of the core amino acid triplet: GDD, SDD, GDN, IDD, ADN and ADD (in order of frequency) (Figure 7). Interestingly, Forgia et al. (2022a) identified additional variations (NDD, GDQ and HDD; besides SDD and ADD) in the ormycoviruses – a recently discovered group of fungi-infecting viruses that are highly divergent from other known RNA viruses.

**Figure 7.**
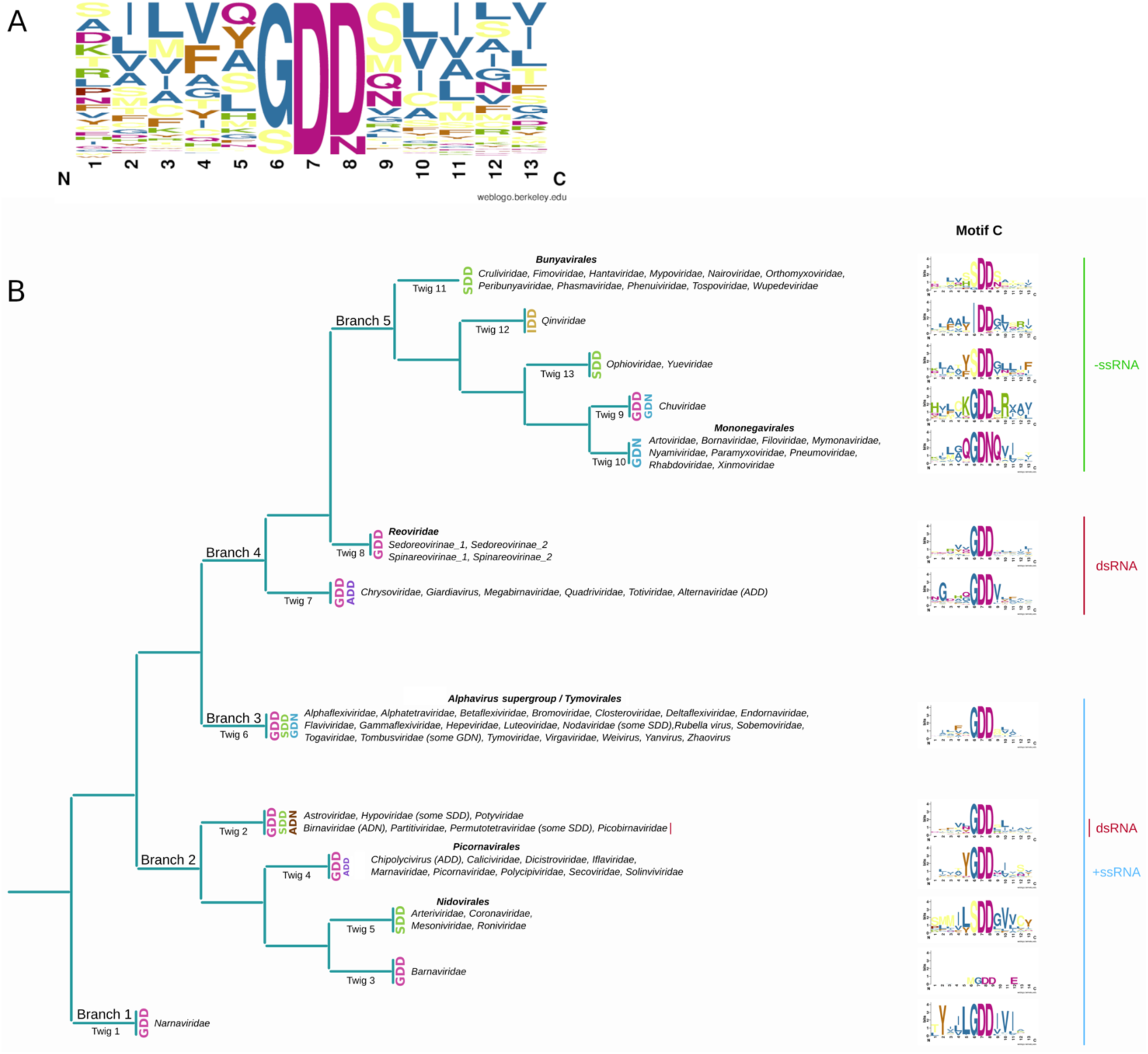
Conservation and diversity in RdRp motif C. **(A)** Sequence logo, produced with WebLogo (Crooks et al., 2004) showing overall amino acid frequencies in the core amino acid triplet and the five flanking amino acids on either side. All non-identical trimmed RdRp sequences in our study were used (cdhit -c 1.0). **(B)** Schematic representation of motif C central triplet variability overlaid on the inferred evolutionary relationships of RNA viruses from Wolf et al. (2018, 2020).

In most +ssRNA virus families, GDD was found to be predominant. As expected, ADD was present in Chipolycipivirus-like sequences (Olendraite et al., 2017) and SDD in members of the order *Nidovirales* (Koonin & Dolja, 1993). Within the group classified to the noda-like pHMM, we noticed four sequences (NC_033077.1, KX883125.1, KX883170.1, GU976287.1) with SDD instead of GDD. Although highly divergent from each other (46– 61% a.a. pairwise identities) these sequences form a distinct monophyletic clade with bootstrap support of 1.0 within the noda-like phylogenetic tree (Supplementary Figures 4A and 5A). Thus the SDD is likely to be a true variation and not the result of sequencing errors or sequencing of defective sequences. This is further supported by the fact that changing a glycine codon (GGN) to one of the serine codons (UCC, UCU, UCA) utilised in these four sequences would require changes in both the first and second position of the codon. We saw a similar phenomenon among permutotetra-like sequences, where a small distinct clade (bootstrap support 1.0) have SDD instead of GDD (Supplementary Figures 4B and 5B). Here, the sequences comprise NC_028381.1, GBSU01004473.1, GBSU01004474.1 and NC_033140.1, where the former three are similar sequences (>90% nt identity) from *Aphis glycines* (soybean aphid), but the latter is more divergent (99% coverage and 83% a.a. identity to NC_028381.1 in the RdRp ORF). In this case the utilised serine codons are AGC and AGU. In the tree of hypo-like sequences, GDD and SDD are both well represented, and in this case there are multiple different clades of SDD-containing or GDD-containing sequences (Supplementary Figures 4C and 5C) indicating multiple switches from GDD to SDD or *vice versa* during the evolution of hypo-like viruses (although it is possible that this could partly be an artefact of poor sequence alignment and non-robust phylogenetic inference). These various cases indicate that an ancestral GDD has mutated to SDD on several different occasions in different groups of +ssRNA viruses. Interestingly, we noticed some cases of tombus-like sequences with GDN, which is normally associated with members of the −ssRNA order *Mononegavirales*. Additional tombus-like sequences with GDN have been noted by Gilbert et al. (2019) who proposed a new family, Ambiguiviridae, to contain these viruses. Again these GDN-containing sequences form a distinct monophyletic clade (bootstrap support 0.97; Supplementary Figures 4D and 5D).

The dsRNA viruses are the least diverse in motif C and mostly have GDD. However, members of the proposed family Alternaviridae (Gilbert et al., 2019) were found to have ADD. As already established for family *Birnaviridae* (Gorbalenya et al., 2002), a deviation from GDD to ADN was observed within birna-like sequences. We noticed a few partiti-like sequences with GDE, namely GFDF01011954.1, GBMJ01010875.1, GBBP01108788.1, GBBP01108783.1, GEFG01022027.1, GEFD01014070.1 and GEEY01016471.1. However, they do not exclusively cluster together. Moreover, they appeared to have fragmented RdRp ORFs and therefore are likely defective sequences, perhaps corresponding to transcribed endogenized viral elements (EVEs) that might have mutated since the original integration into the host genome, leading to a broken ORF and mutation of the aspartic acid codon to a glutamic acid codon which only requires a single nucleotide change. Notably, partitivirus sequences have been found to be particularly frequently integrated into their host genomes (Chiba et al., 2011). Similarly, a single betaflexi-like sequence, GAGH01076983, with GDE clearly contains a fragmented RdRp ORF and therefore is also presumably defective.

The −ssRNA viruses employ many different motif C variations and GDD is not the most frequently used central triplet. In line with previous knowledge (te Velthuis, 2014), members of the order *Bunyavirales* and the families *Orthomyxoviridae*, *Ophioviridae* and *Yueviridae* as well as the TSA sequences which best match the pHMMs of these families, have SDD, whereas all members of the order *Mononegavirales* have GDN. Uniquely, IDD is represented within the sequences that matched the *Qinviridae* pHMM, as observed previously (for example by Charon et al., 2022). Members of the family *Chuviridae* predominantly have GDD though we also noticed three distinct sequences with GDN. Since these sequences (GDRW01001314.1, GDRW0122005.1 and NC_033704.1) are distinct and come from two unrelated projects, but form a monophyletic clade within the chu-like virus phylogeny (Supplementary Figures 2, 4E and 5E), the GDN is likely to be a true variation and not the result of sequencing errors or sequencing of defective sequences.

### Evolutionary relationships of the RdRp above the family level

RNA viruses are extremely divergent at the level of primary sequence and relationships above the level of family are often unclear due to difficulties in making robust sequence alignments for phylogenetic analysis. We sought to utilise our large number of RdRp sequences to further investigate higher level evolutionary relationships. We noticed that each RdRp ORF usually had statistically significant matches to multiple pHMMs (*p* < 10^−6^). Indeed some ORFs matched as many as 11 different RdRp profiles. Therefore we investigated which matched pHMMs tended to occur together, as this might uncover high-level evolutionary relationships. For this analysis, we used the classified group of RdRp sequences from nr/nt, ref and TSA combined. All pHMM match scores were sorted, and the best and the second best matched pHMM for each ORF were taken to the next step. A heatmap and a network diagram were used to visualise the co-occurrence of different pHMMs (Figure 8, Supplementary Figure 6).

**Figure 8.**
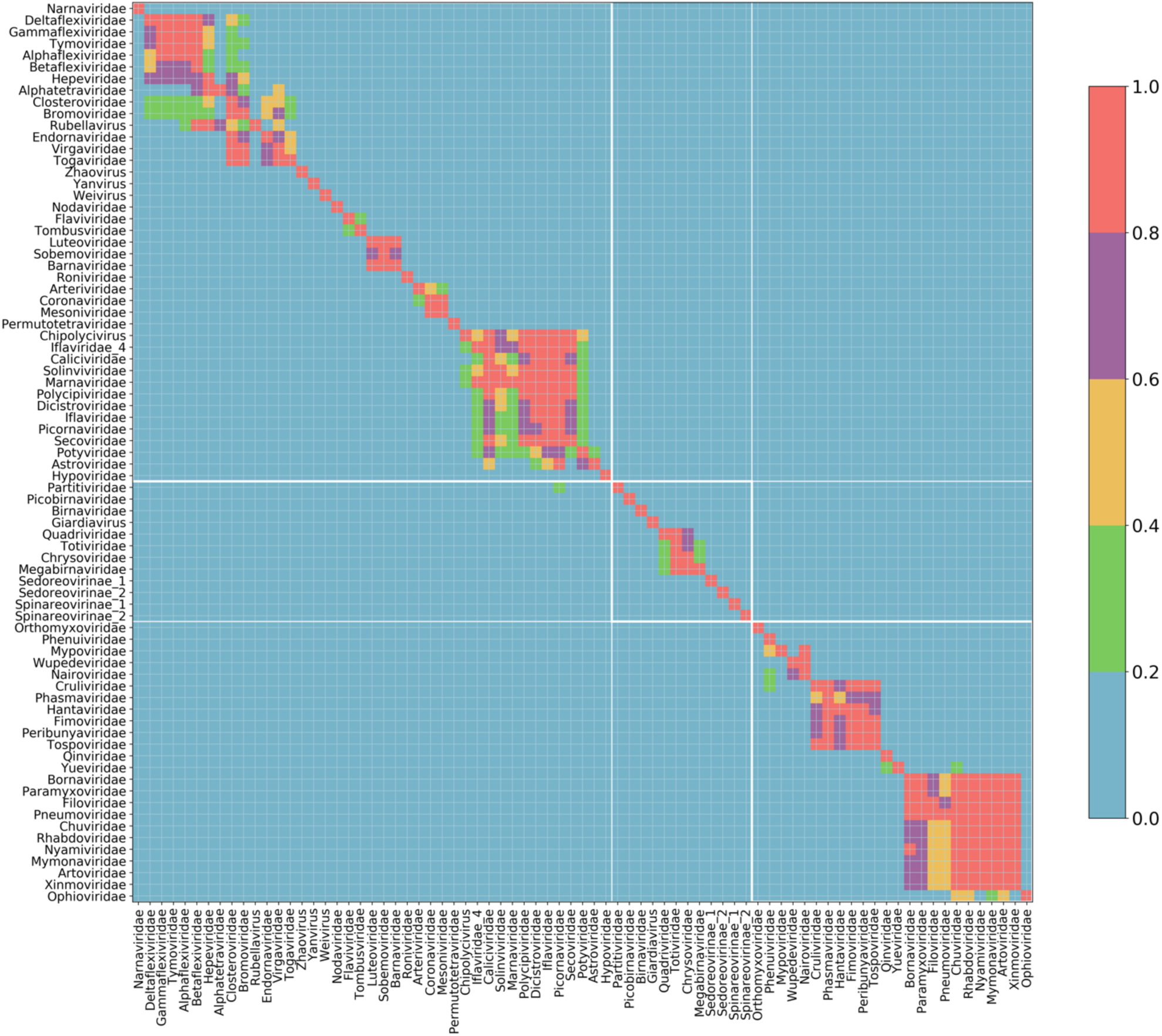
Heatmap of pHMM match co-occurrences for each RdRp sequence. All classified-group ref, nr/nt and TSA RdRp ORFs were used (Supplementary Dataset 1). For each group on the y-axis (best match pHMM), the number of co-occurrences with each group on the x-axis (second best match pHMM) was determined and the count was normalised by the maximum count for the group given on the y-axis. Thus 1.0 is the highest co-occurrence score whereas 0.0 corresponds to pairs of pHMMs that were never matched by the same sequence.

Firstly, it was clear that endorna-like RdRp sequences match the RdRp profiles of +ssRNA rather than dsRNA viruses as their second best match. When we performed this analysis, *Endornaviridae* was considered to be a dsRNA virus family. However, a phylogenetic association with +ssRNA viruses was observed by Roossinck et al. (2011), and Wolf et al. (2018) inferred that endornaviruses, which in fact are capsidless, had been erroneously labelled as dsRNA viruses. In our analysis, the most closely related families to *Endornaviridae* were *Virgaviridae*, *Togaviridae*, *Closteroviridae* and *Bromoviridae* which were recently grouped into the newly established order *Martellivirales*. These co-occurrences also connect families within other orders such as the *Tymovirales* (*Alphaflexiviridae*, *Betaflexiviridae*, *Deltaflexiviridae*, *Gammaflexiviridae*) and the *Hepelivirales* (*Hepeviridae*, *Alphatetraviridae*) (Koonin & Dolja, 1993; Koonin et al., 2015; Wolf et al., 2018), and more loosely link these three orders together, consistent with the 2021 ICTV taxonomy which groups *Martellivirales*, *Tymovirales* and *Hepelivirales* into the class *Alsuviricetes* (phylum *Kitrinoviricota*).

Co-occurrence outside of the same Baltimore group was also apparent for the dsRNA *Partitiviridae* and the +ssRNA *Picornaviridae*. For 20–40% of the RdRp ORFs that best-matched the *Partitiviridae* pHMM, the only other match was the *Picornaviridae* pHMM. This result is consistent with the placement by Wolf et al. (2018) of *Partitiviridae* together with multiple groups of +ssRNA viruses in Branch 2, making the recently established phylum *Pisuviricota*. The *Picornaviridae* pHMM contained many divergent sequences, enabling it to accommodate and tolerate very high diversity. This may explain why partiti-like sequences more easily had second-best matches to the *Picornaviridae* pHMM as opposed to the pHMMs of the other *Picornavirales* families. The members of the order *Picornavirales* formed a robustly connected group, with strong links also to the *Potyviridae* (order *Patatavirales*) and *Astroviridae* (order *Stellavirales*), while other members of class *Pisoniviricetes* clustered elsewhere.

For −ssRNA viruses, families within the class *Monjiviricetes* formed a very distinctive group, incorporating the *Mononegavirales* and the *Chuviridae* (the only member of the order *Jingchuvirales*). The *Chuviridae* provide a tentative link between this class and two other classes of −ssRNA virus – the *Yunchangviricetes* (*Yueviridae*) and *Chunqiuviricetes* (*Qinviridae*). In contrast, the *Bunyavirales* group was split into two clusters. The first cluster comprised *Phenuiviridae*, *Mypoviridae*, *Wupedeviridae* and *Nairoviridae* whereas the second comprised *Cruliviridae*, *Phasmaviridae*, *Hantaviridae*, *Fimoviridae*, *Peribunyaviridae* and *Tospoviridae*. Interestingly, sequences classified to the *Ophioviridae* pHMM sometimes had secondary matches to pHMMs of families within the *Mononegavirales* order, but not *vice versa*. For dsRNA viruses there was a clear clustering of *Quadriviridae*, *Totiviridae*, *Chrysoviridae* and *Megabirnaviridae* that form the recently established order *Ghabrivirales*. Interestingly, and again consistent with the phylogeny of Wolf et al. (2018) and the classification of Aiewsakun and Simmonds (2018), the *Giardiavirus* pHMM did not form part of this cluster even though the genus *Giardiavirus* is currently classified in the *Totiviridae* family. Another order-like clustering was observed for the *Luteoviridae*, *Sobemoviridae* and *Barnaviridae* pHMMs (note that our *Sobemoviridae* pHMM corresponds to what is now designated genus *Sobemovirus* in family *Solemoviridae* along with *Polemovirus*, *Polerovirus* and *Enamovirus*; and our *Luteoviridae* pHMM – following the Aiewsakun and Simmonds groupings – contains only *Enamovirus* sequences).

### Sensitive detection of divergent RdRps using pHMMs

To search for novel and unusual sequences, the most interesting group of putative RdRps is the unclassified group. We placed 361 sequences within the unclassified group, of which 142 are TSA-derived (Supplementary Figure 7). First, we investigated the lengths of these contigs, their full RdRp-encoding ORFs, and the RdRp core (full ORF trimmed to the best pHMM match positions) (Supplementary Figure 8). The RdRp-encoding contigs varied from 700 nt to 35,913 nt, with 51 sequences longer than 10,000 nt. The full RdRp-encoding ORFs varied from 240 to 8,398 codons, with 185 out of 361 longer than 1,000 codons. After trimming these ORFs to the pHMM match positions, the sequences were 300–400 a.a. in length, as expected for the core of an RdRp. The unclassified group contains particularly divergent viruses, with a mean of 26.1% identity to the most similar reference sequence, compared to 64.6% for the classified sequences (measured with BLASTP against sequences included in the input pHMMs, excluding self matches for the GenBank sequences) (Supplementary Figure 9).

We also checked which classified family pHMM each of the 361 unclassified sequences best matched. By our definition of unclassified sequences, in these cases the match score is very low though still statistically significant. We found that unclassified sequences best matched only 43 of the 77 pHMMs (Supplementary Figure 7). It is important to note however that some pHMMs were created using only very similar input sequences and therefore are less able than other pHMMs to match related but divergent sequences. The highest proportion of unclassified sequences matched the *Hepeviridae* pHMM. There were 32 such sequences and 18 of them were TSA-derived. For the *Picornaviridae* pHMM, there were 18 sequences and 11 of them were TSA-derived. In the case of the *Qinviridae* pHMM, all 11 matched unclassified sequences were from the TSA database.

Within the unclassified sequences, we found some clades – such as family *Arenaviridae* and genus *Sinaivirus* – that correspond to known virus taxa for which for various reasons we had not included pHMMs in our analysis. These provided a useful control, demonstrating the ability of our pipeline to find new family-level groups or divergent singletons which sometimes showed very remote similarity to existing taxonomic groups. There was also a group of five very long *Nidovirales* sequences – corresponding to a group of viruses with the largest known RNA genomes to date – which were published (Debat, 2018; Saberi et al., 2018) but not yet classified when we performed our analysis. Sometimes newly identified unclassified-group sequences formed new sister clades to the clade of their best-match pHMM. Although there were multiple such instances, below we describe examples among toti-like and giardia-like viruses. We also highlight a divergent mononegavirus with apparently splicing-dependent RdRp expression, besides new clades of orthomyxo-like viruses.

### An expansion of the toti-like viruses

There were 22 unclassified sequences which had the best match to the *Totiviridae* pHMM. Among these sequences, six were TSA-derived and the shortest one was 2,697 nt in length. Based on genome organisation, they appeared to resemble typical *Totiviridae* viruses with a capsid-encoding ORF followed by an RdRp-encoding ORF. To see if these unclassified sequences form a separate clade, and how phylogenetically different they are from sequences with a strong match to the *Totiviridae* pHMM, we compared the 22 sequences with classified sequences. There were 292 sequences classified to the *Totiviridae* pHMM; therefore we discarded sequences which were shorter than 400 a.a. in length for the RdRp core region and that shared more than 70% similarity (CDHIT -c 0.7) to a longer sequence, leaving 80 sequences. The 22 unclassified and remaining 80 classified sequences were used to produce a PhyML phylogenetic tree (Supplementary Figure 10). Four main clades were observed, two of which contained only classified sequences and covered all ref and nr/nt sequences of the respective genera: *Totivirus* (clade 1) and *Victorivirus*, *Leishmaniavirus* and *Trichomonavirus* (clade 2).

A third clade contained 16 ref or nr/nt and 5 TSA sequences and, of these, four were classified sequences and the remainder were unclassified sequences. A strongly supported sub-clade within clade 3 contained 10 sequences of 8.1–9.4 kb in length, substantially longer than most toti-like sequences. All 10 sequences were found in fungal hosts – nine in Ascomycota and one in Basidiomycota. These sequences correspond to a previously proposed new family, Fusagraviridae (Wang et al., 2016; Lee et al., 2017; Arjona-Lopez et al., 2018). This family is not yet recognised by the ICTV but is supported by our analysis as a monophyletic group distinct from the *Totiviridae*. Another sub-clade within clade 3 contains Ustilago maydis virus H1, currently classified in ICTV as part of the *Totivirus* genus. All the other members of genus *Totivirus* fall firmly within clade 1, which suggests this species should be reclassified. In our phylogeny, Ustilago maydis virus H1 clusters with a member of the *Botybirnavirus* genus (which is currently not classified above genus level), besides diatom colony associated virus 17 types A and B (both of which have not been taxonomically classified beyond a provisional link to the *Totiviridae*). Our phylogeny would support the addition of the *Botybirnavirus* genus to the *Ghabrivirales*, the order which contains family *Totiviridae*, but not to the *Totiviridae* family itself. By incorporating the results of other recent studies, it is apparent that the sub-clade of clade 3 containing our three TSA sequences (from the stone-fly *Perla marginata*, the red alga *Kappaphycus alvarezii*, and the orchid *Chiloglottis trapeziformis*) corresponds to the totivirus-like clade identified by Kartali et al. (2019) as containing their Umbelopsis ramanniana virus 3. This clade also incorporates the clade shown by Charon et al. (2021) as containing their Chrysaor toti-like virus and Laestrygon toti-like virus, and the clade shown in Shi et al. (2016) to contain diatom colony associated dsRNA virus 17, Beihai barnacle virus 15, Hubei toti-like virus 5, Beihai sesarmid crab virus 7 and Beihai razor shell virus 4. Mifsud et al. (2022) identified Elkhorn sea moss toti-like virus – likely to be the same virus as our red alga TSA sequence – and a number of other plant-associated viruses which also fall into this clade. Thus, this grouping brings together our three novel sequences with a number of previously scattered known sequences.

The fourth clade contains five sequences from arthropods, all with an unclassified status in our analysis – two TSA sequences (GFQL01013152.1 from the moth *Carposina sasakii*, GBNZ01013113.1 from a *Heterodontonyx* sp. wasp) and three ref or nt/nt sequences (NC_007915.3 – penaeid shrimp infectious myonecrosis virus, NC_033467.1 – Wuhan insect virus 31, and NC_032948.1 – Hubei toti-like virus 18).

### An expansion of the giardia-like viruses

In addition to the *Totiviridae* pHMM, we also had a separate profile for the Giardia lamblia virus RdRp. In total, there were 65 sequences (nr/nt, ref and TSA) which had the best match to this pHMM, with lengths ranging from 308 to 12,427 nt (the longest being LC333746.2, Rosellinia necatrix megatotivirus 1; Arjona-Lopez et al., 2018). Among our identified sequences there were 24 with lengths >6,000 nt (cf. the Giardia lamblia virus reference sequence NC_003555.1 has 6,277 nt). As the *Giardiavirus* RdRp is translated via ribosomal frameshifting, for all these 65 sequences, we defined ORFs between stop codons to ensure coverage of the entire RdRp. For each contig, the ORF with longest BLASTP match to the Giardia lamblia virus RdRp a.a. sequence was selected. The translated ORFs were aligned and a phylogenetic tree was generated with PhyML (Supplementary Figure 11).

The original Giardia lamblia virus sequence formed a clade together with seven unclassified nr/nt and ref sequences and one TSA sequence (GBYF01047348.1), which was identical to a known nr/nt sequence (MG256177, Gigaspora margarita giardia-like virus 1). A neighbouring clade to this one encompassed five nr/nt and ref sequences together with 13 TSA sequences – one divergent sequence from a microfungus TSA sample (*Rhizopus oryzae*, GDUK01008098.1) and 12 sequences from five crustacean TSA samples (swimming crab, great spider crab, signal crayfish and two species of *Proassellus* isopods). These 12 TSA sequences together with Wenzhou crab virus 5 form a strong clade (bootstrap support 1.0) of crustacean-associated giardia-like viruses. The remaining sequences comprised 30 TSA sequences from many different hosts which loosely clustered together with 7 GenBank unclassified viruses.

### Amphibian-associated orthomyxo-like viruses

Within the *Orthomyxoviridae* family, the four influenzavirus genera form a highly supported monophyletic clade of vertebrate-infecting viruses (Figure 9). In our study, we identified additional influenza-like sequences in three different amphibian species. In the TSA dataset GFMT01 (cane toad, *Rhinella marina*) we found a full PB1-encoding sequence (GFMT01051794.1, 2,350 nt), and in the TSA dataset GECV01 (ornate chorus frog, *Microhyla fissipes*) we found three PB1-matching contigs (GECV01084760.1, 713 nt; GECV01039268.1, 338 nt; and GECV01050644.1, 473 nt). Using amino acid sequences derived from these PB1 contigs, we searched online amphibian TSA datasets (TBLASTN) and identified another PB1-encoding contig, namely JV207023.1 (1,419 nt) in an *Ambystoma mexicanum* (axolotl) TSA dataset (NCBI BioProject PRJNA157225). Comparison of the 7,332 contigs in BioProject PRJNA157225 with *Orthomyxoviridae* NCBI reference proteins using BLASTX revealed additional influenzavirus-like contigs, including JV207023.1, JV205532.1 and JV206720.1 that matched PB1 and could be merged into a 2018 nt sequence. These amphibian-associated sequences – which were also found concurrently by Parry et al. (2020) – all cluster within the influenzavirus clade (Figure 9).

**Figure 9.**
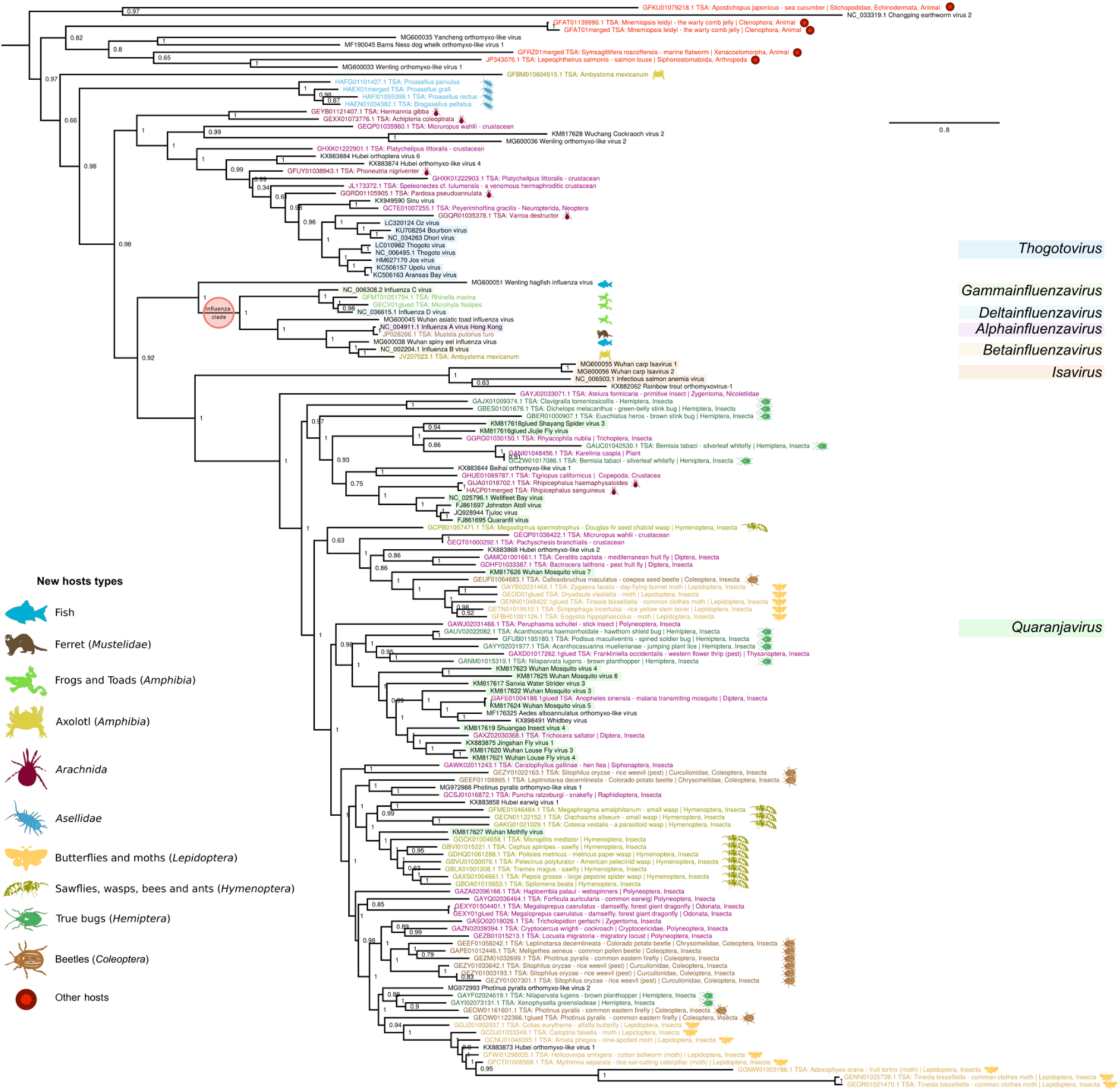
Phylogenetic tree of sequences (classified or unclassified) with best match to the orthomyxovirus-like pHMM. Sequences shorter than 100 a.a. were removed and then sequences with >95% identity were clustered and only the longest sequence in each cluster was retained as a representative. NCBI accession numbers, virus names, TSA target organism names, and group-representative icons/colours are shown (key at left). Currently defined genera are identified with coloured highlighting (key at right). Sequences labelled as “merged” derive from merged overlapping contigs. Sequences labelled as “glued” comprise multiple concatenated ORFs from a contig that was inferred to likely have sequence quality issues which introduce stop codons (e.g. via frameshift errors – common with 454 sequencing) or potentially derive from mutated endogenised viral elements, EVEs).

Surprisingly, we also identified influenza-like PB1-encoding sequences in two fish TSA datasets (*Salaria pavo* – NCBI BioProject PRJNA329073 and *Nibea albiflora* – NCBI BioProject PRJNA359138). We identified sequences for the other segments by searching the respective TSA datasets using TBLASTN with *Orthomyxoviridae* NCBI reference proteins as queries. This revealed 9 and 15 influenza A virus-like contigs for *Salaria pavo* and *Nibea albiflora*, respectively. Using BLASTX, we identified all eight virus segments and found the encoded proteins to have 93–100% a.a. identity to influenza A virus proteins (Supplementary Tables 2 and 3). These identity levels are typical for different strains of influenza A virus (Supplementary Table 4) and much higher than identity levels between, for example, the homologous proteins of influenza A and B viruses (Supplementary Table 5). Thus, the fish TSA dataset sequences may be considered to represent the influenza A virus species and, since fish are not known hosts of influenza A virus, are likely contaminants (a conclusion also reached by Parry et al., 2020) – perhaps from bird faecal material or lab contamination. Similarly to Mifsud et al. (2022), who found influenza A virus in 16 plant transcriptomic datasets, we also found influenza A virus sequences in a plant dataset (*Thlaspi arvense* – NCBI BioProject PRJNA183631), with the PB1 fragment GAKE01008984.1 having 99% nt identity to the avian influenza A virus sequence CY149610.

Among the four unclassified sequences which had the best match to the *Orthomyxoviridae* PB1 pHMM (Supplementary Figure 7), there was a 2,551 nt sequence, GFBM010604515.1, from another *Ambystoma mexicanum* dataset. In contrast to the influenzavirus-like amphibian-associated sequences mentioned above and in Shi et al. (2018) and Parry et al. (2020), this sequence was far more divergent (e.g. 22.7% a.a. identity, 60% coverage in a TBLASTN comparison with influenza B virus PB1) and did not cluster within established genera (Figure 9). Note that the tree of orthomyxo-like viruses (Figure 9) does not extend to other *Articulavirales* clades such as the *Amnoonviridae* (Turnbull et al., 2020) and the unclassified gecko-derived Lauta virus (Ortiz-Baez et al., 2020), and the PB1 encoded by GFBM010604515.1 is more closely related to *Orthomyxoviridae* PB1 proteins than the PB1 proteins of these other virus groups.

Orthyomyxoviruses have segmented genomes, with typically 6–8 segments. In an attempt to find other segments of this novel virus, we used TBLASTN to query 164 NCBI *Orthomyxoviridae* protein reference sequences (covering all segments) against the *Ambystoma mexicanum* TSA dataset (NCBI BioProject PRJNA300706). However, the only match was to the original contig, GFBM010604515.1. To increase sensitivity, we downloaded all TSA contigs (∼1.5 × 10^6^) from BioProject PRJNA300706 and, using HMMsearch, compared them with new pHMMs generated for *Orthomyxoviridae* reference proteins. In this manner, we identified contigs encoding the three replicase components: GFBM010604515.1 (2,551 nt, PB1), GFBM010554880.1 (2,714 nt, PB2) and GFBM010538345.1 (2,195 nt, PA). All three contigs contain the ORF stop codon. Furthermore (after reverse complementing where appropriate) all three contigs have an identical 5′-end AAAAAGCAGU sequence (plus 0–2 extra 5′-terminal nucleotides) consistent with the conserved segment ends expected for orthomyxovirus sequences. Thus, the encoded proteins PB1, PB2 and PA appear to be full-length.

We applied BLASTP to the three retrieved protein sequences, querying against the NCBI nr protein database. The best match for PB1 was Sinu virus (bitscore 72.4, 91% cover, 20% identity, evalue 6 × 10^−9^), followed by Neke Harbour virus (bitscore 71.2, 33% cover, 25% identity, E-value 10^−8^), Wuhan mosquito virus 4 (bitscore 69.3, 41% cover, 24% identity, E-value 5 × 10^−8^), and many influenza B virus sequences. The best matches for PB2 were all influenza A virus sequences (top match bitscore 60.8, 43% cover, 21% identity, E-value 2 × 10^−5^). Finally, the PA protein matched only Barns Ness dog whelk orthomyxo-like virus 1 (bitscore 58.2, 42% cover, 24% identity, E-value 10^−4^). Thus, this novel amphibian-associated virus appears more closely related to orthomyxoviruses than to other virus families, though it falls outside of currently defined *Orthomyxoviridae* genera. It is possible that the host species of this virus is not in fact the axolotl but might instead be a contaminant (e.g. an invertebrate).

### A new clade of orthomyxo-like viruses associated with isopods

In addition to the amphibian-associated sequences noted above, within the tree of all sequences with the best match to the *Orthomyxoviridae* pHMM, we also saw some new clades that fall outside of currently recognised genera (e.g. the top 14 sequences in Figure 9). One of these clades was specific to TSA datasets sampling the family Asellidae – a group of isopod crustaceans. A second clade contained a mixture of previously published virus sequences together with sequences from marine flatworm (*Symsagittifera roscoffensis*), salmon louse (*Lepeophtheirus salmonis*) and warty comb jelly (*Mnemiopsis leidyi*) TSA datasets. The *L. salmonis* sequence was previously identified by Waldron et al. (2018) and clusters with their Barns Ness dog whelk orthomyxo-like virus, whereas the other two sequences appear to be novel. A third clade contained the previously published Changping earthworm virus 2 and a sequence from a sea cucumber (*Apostichopus japonicus*) TSA dataset.

For the Asellidae-associated clade, we initially identified nine contigs but, after merging two overlapping contigs, there were eight PB1-encoding sequences from six different TSA datasets representing six different Asellidae species within NCBI BioProject PRJEB14193 (Supplementary Table 6). The eight contigs have 65–98% nucleotide identity to each other (Supplementary Figure 12A). Using BLASTX, we compared these sequences with *Orthomyxoviridae* NCBI reference PB1 proteins and found them to be highly divergent, with all a.a. identity levels <32% (Supplementary Table 7; Supplementary Figure 12B). When we compared the Asellidae-associated PB1 sequences with the entire NCBI non-redundant (nr) protein database using BLASTX, aside from orthomyxovirus-like sequences, there were no other significant matches, thus confirming a closer relationship to the *Orthomyxoviridae* family than to any other viruses (including other viruses in the *Articulavirales* order). For the phylogenetic tree, we discarded shorter sequences that had >95% amino acid identity to longer sequences, leaving the four PB1 Asellidae-associated sequences shown in Figure 9. The four sequences are all >1,000 nt and two are >2,000 nt, making them possibly full-length coding sequences (Supplementary Table 6).

Using TBLASTN, we queried all *Orthomyxoviridae* NCBI reference proteins against the Asellidae BioProject PRJEB14193 to search for contigs matching to orthomyxovirus proteins other than PB1. Four contigs showed significant matches to the PA protein of thogotoviruses. When these contigs were queried against the Asellidae BioProject using TBLASTX, a total of 11 contigs from 5 host species were found that separated into 3 groups (97–100% a.a. identity within a group) plus a fourth small fragment. The longest segments by group were HAEN01028927.1 (2,008 nt), HAFG01097557.1 (1,986 nt), and HAEX01036584.1 (1,799 nt). In comparisons with thogotovirus PA (YP_145795), these sequences had coverage values of 32–51% and amino acid identities of 21.7–24.0%. Compared to each other, coverage and amino acid identities were in the range of 86–97% and 51.2–59.6% respectively.

The other divergent orthomyxovirus-like clades and sequences mentioned above comprise the *Ambystoma mexicanum* TSA sequence GFBM010604515.1 discussed above, five other TSA sequences from a variety of organisms, and four previously identified but not currently classified NCBI nr/nt viruses (Shi et al., 2018; Waldron et al., 2018) (upper part of tree in Figure 9). When these sequences were compared with the entire nr protein database using BLASTX, most of the best hits were encoded by the other nr/nt sequences identified in this group, with identities (variable coverage) mostly in the range 21–34% (Supplementary Table 8). There were no hits to viruses other than orthomyxo-like viruses. Among the *Orthomyxoviridae*, there were unclassified *Orthomyxoviridae*, *Quaranjavirus* and *Isavirus* PB1 matches (identities 20–30%, coverage 15–50%, bit scores 41.2–70.9, E-values 0.038 to 3 × 10^−11^). Since these sequences do not cluster according to host taxa, and pairwise identity scores are relatively low, it is perhaps premature to propose new taxa at this stage. Nonetheless, these sequences enrich the apparent host range of orthomyxo-like viruses to include (with the caveat of potential contamination) such hosts as, the marine flatworm (phylum Xenacoelomorpha), warty comb jelly (phylum Ctenophora) and sea cucumber (phylum Echinodermata), in addition to the published orthomyxo-like viruses from whelk (phylum Mollusca), earthworm (phylum Annelida), and the well-established *Orthomyxoviridae* host taxa Arthropoda and Vertebrata.

### A divergent mononegavirus with splicing-dependent RdRp expression

While looking at genome graphs of unclassified sequences, we noticed an unusual and divergent TSA sequence, GEZL01043288.1 (from a common ragweed dataset, *Ambrosia artemisiifolia*). By analysing the original TSA dataset, GEZL01, we were able to extend GEZL01043288.1 at the 3′ end with contigs GEZL01043287.1, GEZL01043289.1 and GEZL01043290.1. The resulting 12,569 nt sequence had best TBLASTX hit to Wuchan romanomeris nematode virus 2 (NCBI nr/nt; KX884441.1; *Nematovirus*, *Lispiviridae*, *Mononegavirales*). This match mapped around RdRp motif C (QGDNQ) where there was 38% identity over 187 a.a. of the RdRp. The original TSA sequence best matched the *Artoviridae* pHMM and, when extended, showed features typical of viruses in the order *Mononegavirales* (Figure 10A). Phylogenetically, the sequence falls in a sister clade to the *Rhabdoviridae*-like family *Lispiviridae* (Figure 10B, Supplementary Figure 13). ORF1 apparently encodes the nucleoprotein (HHpred E-value 3.6 × 10^−9^, PDB_mmCIF70_14Oct:1N93_X), whereas we were not able to identify the putative ORF2 and ORF3 products by homology search with HHpred. Downstream of ORF3, the 3′ region contains several disjoint ORFs, three of which have highly significant a.a. matches to the *Mononegavirales* L protein (TBLASTN against KX884441.1, E-value 5 × 10^−13^ for the shortest fragment), and together cover its RdRp, capping, connector and methyltransferase domains (HHpred of concatenated a.a. sequences has an E-value 5.1 × 10^−189^ hit to PDB_mmCIF70_14Oct:6V85_A) (Figure 10A). Further inspection revealed the presence of three introns (see below; Figure 10A, Supplementary Figure 14). When these introns are not spliced the RdRp core is split between disjoint ORFs, whereas removal of all three introns fuses the RdRp/L protein-coding region into a single long ORF.

**Figure 10.**
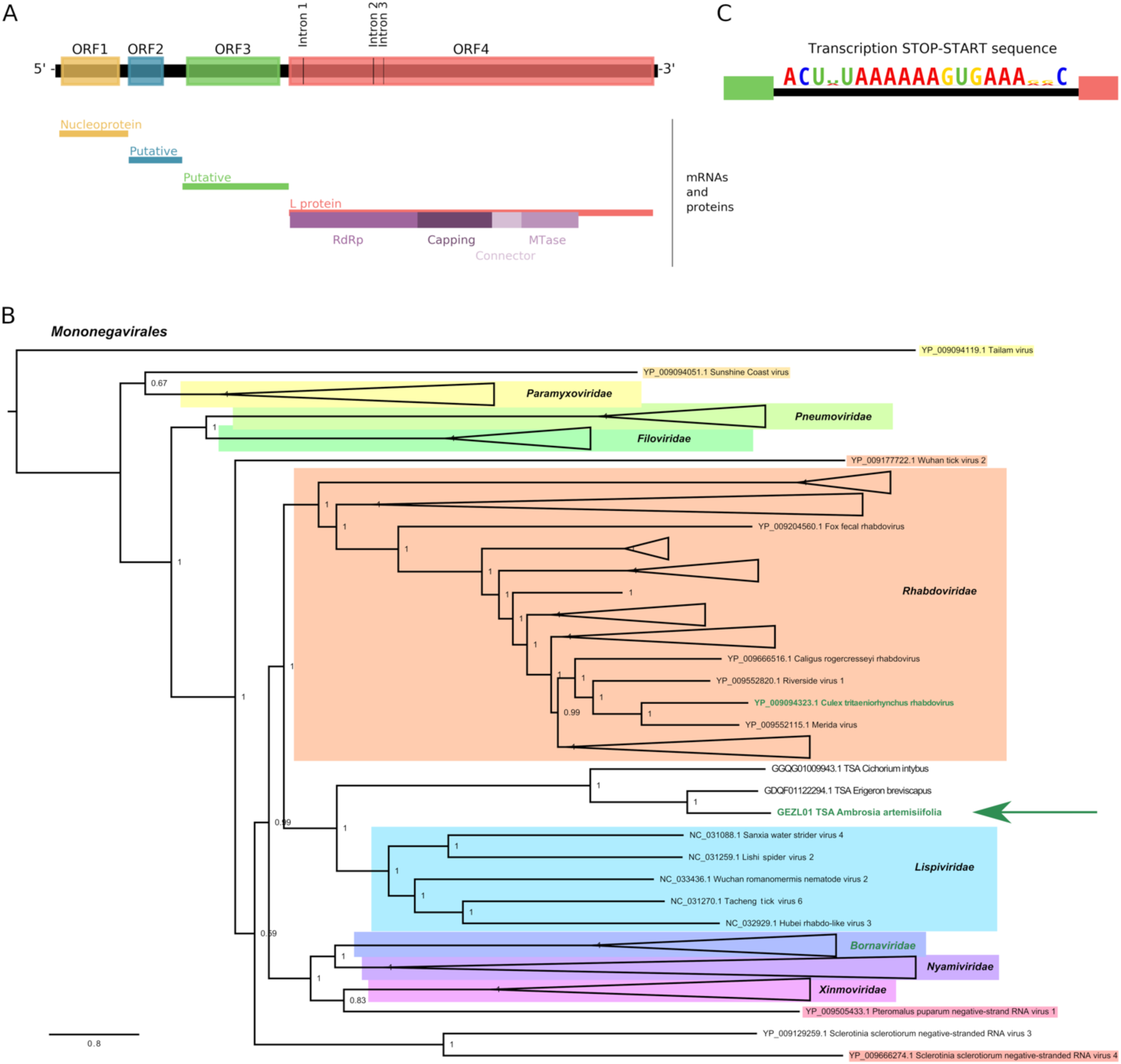
Splicing in a new rhabdo-like virus sequence. **(A)** Genome map of the rhabdo-like virus derived from the GEZL01 TSA dataset. The diagram illustrates ORFs in the antigenome after removal of the identified introns. The positions of the removed introns are indicated. Putative transcription stop-start (TSS) sequences were identified between the ORFs and the corresponding inferred mRNAs and their products (where identified) are indicated below as well as domains of the L protein. **(B)** Phylogenetic tree of *Mononegavirales* L protein sequences showing the placement of the GEZL01-derived rhabdo-like virus. For visual convenience, some clades are collapsed into isoceles triangles. Names of sequences/clades with known splicing are written in green. See Supplementary Figure 13 for the complete tree. **(C)** Sequence logo generated from the three identified copies of the putative TSS sequence (shown in the antigenome sense), using CIAlign v 1.1.0 (Tumescheit et al., 2022).

Evidence for the introns initially came from a comparison of the TSA contigs GEZL01043287.1 and GEZL01043290.1 which are identical except that GEZL01043290.1 has a single 98 nt deletion (intron 2) with canonical GU/AG exon/intron boundaries and flanking sequences that closely match the splice site consensus sequence of *Arabidopsis thaliana* (Supplementary Figure 14; Brown et al., 1996); the human splice site consensus sequence is very similar). Comparison of the TSA contigs GEZL01043288.1 and GEZL01043289.1 revealed a 132 nt intron (intron 1) that is deleted in GEZL01043289.1, whereas we found the 131 nt intron 3 manually. Like intron 2, introns 1 and 3 have canonical GU/AG boundaries and favourable (though less extensive) flanking intron-exon junction sequences (Supplementary Figure 14). All three introns are AU-rich (68.2%, 71.4% and 68.7%, respectively) compared with a mean AU fraction for the entire 12,569 nt sequence of 56.5%.

The antigenomes of viruses in the −ssRNA virus order *Mononegavirales* typically contain a number of consecutive coding ORFs separated by intergenic regions. In the negative-sense template, these intergenic regions contain transcription stop-start signals (reviewed in Ogino & Green, 2019) which direct the production of a series of positive-sense polyadenylated mRNAs, one mRNA species for each main ORF. To better define the RdRp ORF initiation site, we next identified the transcription stop-start motif. For this, we aligned intragenic regions and used GLAM2 (Frith et al., 2008) to identify enriched motifs. The motif comprising the stop-start signal ACU(U/A)UAAAAAAGUGAAA(G/A)(G/A)C (represented in the positive-sense; Figure 10C) was found ending at nucleotide positions 1,500, 2,595 and 4,758. The first two signals could produce transcripts for ORF2 (1,503–2,177) and ORF3 (2,684–4,540), respectively. The third signal could produce a transcript for the L protein ORF if all three introns are removed (AUG initiation codon at nt 4,771–4,773; see Supplementary Figure 15 for protein sequence). In the absence of other transcription stop-start signals, initiation at this AUG codon and utilisation of all three splice sites appears to be the only way for the virus to express a functional RdRp/L protein. We used TBLASTN to compare the entire 2,338 a.a. predicted L protein sequence against the related TSA sequences (identified by TBLASTN) – GDQF01122294.1 (*Erigeron breviscapus* TSA) and GGQG01009943.1 (*Cichorium intybus* TSA) (Figure 10B). Both sequences display 99% coverage of the predicted L protein and, importantly, a lack of alignment gaps at the identified exon-exon junctions in the GEZL01-derived sequence supports introns 1–3 being functionally utilised (Supplementary Figure 16).

Next we wanted to check the abundance of viral reads in the datasets and also identify reads which would cover or span spliced regions of the sequence. For this purpose, we used HISAT2 (Kim et al., 2019) to map the raw reads from the corresponding BioProject (NCBI accession PRJNA335689, three different datasets) to spliced and unspliced versions of the viral sequence. The three datasets represent three different *Ambrosia artemisiifolia* (common ragweed) plant tissues (female flowers, male flowers, leaves) (Virág et al., 2016) and there was a clear difference in coverage between the three datasets (Supplementary Figure 17A). Interestingly, much higher levels of virus were found in the male flower sample, whereas the lowest levels were present in the leaf sample. As expected, we saw dips in mapped read counts corresponding to the transcription stop-start sites (Supplementary Figure 17A). We also saw dips corresponding to the exon-exon junctions, indicating that a substantial proportion of the RNA density in the L ORF region comes from RNA containing the intron sequences. This could be from pre-spliced L mRNAs, genomic vRNA, antigenomic cRNA, or transcripts from which the introns are never spliced. The latter could provide a mechanism to reduce or even temporally regulate L protein expression. Next, we plotted only spliced reads as determined by HISAT2 over the L-protein-encoding region (Supplementary Figure 17B). All three introns mentioned above were supported by multiple spliced reads. In addition, HISAT2 uncovered an additional intron (intron 4) besides alternative 5′ donor sites for introns 1 and 4 (Supplementary Figure 14). However utilisation of alternative introns 1 or 4 would disrupt the L ORF leading to a greatly truncated L protein, whereas excision of intron 4 would lead to a 70 a.a. deletion compared to the *Erigeron breviscapus* and *Cichorium intybus* sequences (see above; Supplementary Figure 16).

The presence of these three related sequences in *Ambrosia artemisiifolia*, *Erigeron breviscapus* and *Cichorium intybus* (three members of the Asteraceae family of flowering plants) besides related virus-derived sequences in more recent plant TSA datasets such as *Cenchrus americanus* (GEUY01006481.1), *Gymnadenia rhellicani* (GHXH01342866.1, GHXH01342865.1, GHXH01230927.1), *Ophrys sphegodes* (GHXJ01055800.1) and *Ophrys fusca* (GHXI01221739.1) suggest plants are the *bona fide* hosts of these viruses.

Splicing is a very rare phenomenon in RNA viruses (excluding retroviruses) and is known only in a few cases – notably bornaviruses (Cubitt et al., 1994; Schneider et al., 1994; Tomonaga et al., 2000; Pfaff & Rubbenstroth, 2021) and various orthomyxoviruses (Inglis & Brown, 1981; Shih et al., 1998; Kochs et al., 2000; Wise et al., 2012), and Culex tritaeniorhynchus rhabdovirus (family *Rhabdoviridae*) where a single 76 nt intron was identified in the RdRp ORF (Kuwata et al., 2011).

### Comparison with BLASTP

As discussed above, all our results were verified with BLASTP against the NCBI nr protein database. In all but 25 cases, the most significant BLAST match was to a protein likely to be derived from an RNA virus.

Using BLAST at this low level of stringency also results in many false positives, reducing its usefulness for large-scale screening. To provide a direct comparison, we performed a BLASTP search using the input RdRp sequences from our pHMMs as a database against two query datasets, our identified viral ORFs and a curated set of known human proteins, the 20,398 reviewed SwissProt proteins in UniprotKB (UniProt Consortium, 2021) which should not be RNA viral in origin. Using the default BLASTP settings plus a relaxed E-value cutoff of 0.05, all of our viral ORFs were detected. However, using this cutoff, significant hits were also detected for 824 of the human proteins. Using HMMSearch, with cutoffs as discussed in the Materials and Methods, with the Uniprot reviewed proteins, there were no false positives.

## Discussion

We generated pHMMs for 77 RNA virus groups and used them to sensitively search the NCBI Transcriptome Shotgun Assembly database. We identified 5,867 RNA virus-derived RdRp-encoding TSA sequences. We supplemented these via a similar search of the NCBI nr/nt virus database. Through this work we have expanded known virus clades and identified new virus clades. We have also illustrated how we can assess virus gene expression strategies by referring back to raw read data to analyse splicing in a new mononegavirus. Our pHMM search was fast enough to make searching >10 billion ORFs feasible, and post-analysis of the identified sequences confirmed high specificity (zero false positives among over 12,000 hits). We were able to detect many viral sequences in the “twilight zone” of sequence similarity (<35% similarity) (Rost, 1999; Cobbin et al., 2021). A list of all the sequences found (as well as representative sequences after clustering by high similarity) and PhyML trees of the different virus groups are available as Supplementary Datasets 1 and 3. Although a small proportion of these sequences may represent transcribed EVEs, or incorporate *in silico* misassemblies, we expect that most are likely to represent *bona fide* RNA viruses. The pHMM models and associated sequence alignments generated are available as Supplementary Dataset 4. We hope that these will be a useful resource for other virus discovery projects. Profile HMMs are not only a more sensitive and specific method than BLAST for finding distant homologues, but also faster (as their use can reduce the number of query-to-subject comparisons).

Although we have shown that our pHMM approach can identify virus groups not included in the original set of pHMMs, they may not be able to identify even more divergent RdRps. Employing an iterative pHMM search method such as JackHMMER (Johnson et al., 2010), whereby newly identified divergent sequences are used to update pHMMs for subsequent searches, might enable identification of even more divergent RdRps (cf. Callanan et al., 2020). Approaches based on predicted secondary or tertiary protein structure such as HHpred (Zimmermann et al., 2018), Phyre2 (Kelley et al., 2015) or AlphaFold (Jumper et al., 2021) could also be useful to find more divergent RdRp sequences (Wolf et al., 2020; Charon et al., 2022; Forgia et al., 2022a; Lee et al., 2022). For example, homology of the quenyavirus RdRp to previously known RNA virus RdRps was detectable with HHpred but not with BLASTP (Obbard et al., 2020) or our approach. On the other hand, pHMM searches are much faster than structural approaches, and this can be a critical issue for high throughput searches.

HMM profiles are sensitive to the input sequences used. In our study, we found that in some cases a family-level pHMM was able to identify many more sequences from one genus than another genus within the same family. Often this could be traced to a bias in sequence representation during pHMM construction. HMMbuild does not phylogenetically weight input sequences. Therefore if one genus is highly “overrepresented” in the profile, the profile will be better at finding similar such sequences. On the other hand, one family-level profile may accommodate the possibility of very high divergence, while another may be very specific, depending on the diversity provided during pHMM construction. When identifying the best match profile for a given sequence, a sequence from the latter family might have a higher score with the former profile as it tolerates more variation. Thus, when building pHMMs, one should focus on representation of diversity rather than just number of sequences.

Re-use of public transcriptomic data is a cost-effective means of exploring the diversity of the RNA virosphere. In most cases, the datasets were obtained for purposes completely different from virus identification – e.g. the divergent axolotl-associated orthomyxo-like sequence came from a transcriptomic dataset for a study on limb regeneration (Bryant et al., 2017). A variety of other studies have also searched for RNA viruses in the NCBI TSA database, or other transcriptomic studies generated without the express purpose of virus discovery, including Cook et al., 2013; Longdon et al., 2015; Mushegian et al., 2016; Olendraite et al., 2017; Gilbert et al., 2019; Käfer et al., 2019; Lauber et al., 2019, 2021; Rosani et al., 2019; Starr et al., 2019; Callanan et al., 2020; Obbard et al., 2020; Ott Rutar & Kordis, 2020; Parry et al., 2020; Wu et al., 2020; Chang et al., 2021; Charon et al., 2021; Paraskevopoulou et al., 2021; Bejerman & Debat, 2022; Dheilly et al., 2022; Lee et al., 2022; Mifsud et al., 2022; Neri et al., 2022; Sidharthan et al., 2022; Zayed et al., 2022. Most of these studies have been limited to certain virus groups and/or certain host groups. Furthermore, although HMM-based search strategies are being used more frequently (e.g. Gilbert et al., 2019; Käfer et al., 2019; Callanan et al., 2020; Charon et al., 2021, 2022; Lauber et al., 2021; Paraskevopoulou et al., 2021; Zayed et al., 2022), most studies to date have relied on BLAST-type search tools.

A recent study by Edgar et al. (2022) queried the entire SRA database of over three million RNA-seq datasets using a novel approach, combining read mapping to RdRp sequences and a novel tool, PalmScan (Babaian & Edgar, 2021). PalmScan identifies RdRp-like sequences using the order and composition of the RdRp A, B and C motifs. This methodology, which allows screening of unassembled sequence read datasets, alongside development of Serratus, a highly optimised computational architecture, allowed screening on an unprecedented scale and identification of over 10^5^ putative RdRp sequences. This work contributes a monumental step forward in the field of virus discovery and provides an extremely valuable resource, while also demonstrating the enormous potential of exploring the diversity of the RNA virosphere using public transcriptomic data. However, the approach used is likely to have reduced sensitivity for more divergent RdRp sequences.

When analysing publicly available data, it is difficult to assess the likelihood of contamination and one must therefore be particularly cautious with host species assignment. Potential sources of contamination include gut contents and microbiota, mould and insects on plant leaves, other internal and external parasites and commensal organisms, contamination during sampling and sample preparation, and contaminated reagents (Cobbin et al., 2021; Harvey & Holmes, 2022; Mifsud et al., 2022). One sign of potential contamination is when an identified virus has a very different TSA target host compared to the host species of similar previously known viruses. For example, in our analysis, we found sequences with high similarity to known influenza A virus sequences in two fish datasets. Influenza A virus is known to infect birds and mammals, whereas known fish orthomyxoviruses are much more divergent. The sequences are therefore very likely to derive from contamination, for example bird faecal material or lab contamination. Although beyond the scope of our study, identification of RNA virus fragments occasionally integrated into host genomes (i.e. EVEs) has been used by others as a means to support linkage of uncharacterized virus taxa to broad host taxonomic groups (Shi et al., 2016).

Despite the aforementioned caveats, the identification of multiple related virus sequences from multiple related host species in different studies lends credence to the assignment of virus-host associations – for example the Asellidae associated clade of orthomyxo-like viruses and the crustacean-associated clade of giardia-like viruses. Thus our study allowed us to assess large scale taxonomic associations and trends in sampled virus diversity. For vertebrate-specific groups such as paramyxoviruses and picornaviruses, we found relatively few new sequences in TSA datasets compared to viruses from non-vertebrate hosts. We also found a substantially larger number of RdRp sequences per dataset in non-vertebrate hosts (∼16 fold higher in plants and arthropods than in vertebrates). One possible explanation may be that vertebrate samples tend not to comprise whole organism samples, with exclusion of contaminating non-target organisms from gut contents and surface material, besides sampling a reduced number of tissues and cell types compared to non-vertebrate studies that often comprise the whole organism or even multiple pooled whole organisms. Another possible explanation may lie in the different purposes for which TSA datasets are generated (e.g. many samples from the same species under different experimental lab conditions versus samples from many different species obtained from the wild). Alternatively, it may be linked to differences in the immune systems of vertebrates (e.g. adaptive immunity) and non-vertebrates (e.g. RNA interference) or stem from other major events in the evolution of eukaryotes (cf. Harvey & Holmes, 2022). In any case, it is clear that there is an enormous amount of unsampled RNA virus diversity – especially in non-vertebrates – and repurposing of existing datasets provides a valuable route to increasing our understanding of virus diversity, taxonomy, evolution and ecology.

## Methods

### Construction of RNA dependent RNA polymerase pHMMs

Initially we used RNA virus groups and sequences from the “GRAViTy” analysis of Aiewsakun and Simmonds (2018). In cases where there was only a small number of sequences in a family-like group, or where some more recently published groups of viruses were not mentioned in Aiewsakun and Simmonds, we searched the NCBI taxonomy and nucleotide databases (April 2018) for additional reference sequences in order to make more representative pHMMs. In total, we used 1,793 RdRp protein and RdRp-containing polyprotein sequences to create 77 pHMMs.

Because many RdRps are contained within longer polyproteins, we wished to trim sequences to a core RdRp region. Therefore, we aligned the sequences within each group with MUSCLE v3.8.31 (Edgar, 2004) and compared the alignments using HHpred (Zimmermann et al., 2018) with the Pfam and PDB databases (Berman et al., 2000; Finn et al., 2014). Based on the second best matching RdRp (to avoid overfitting if the best match Pfam pHMM contained sequences from the same group), we cropped each alignment from both ends. Where appropriate, the alignments and coordinates for trimming were manually curated based on current knowledge of the families. The cropped alignments were formatted to Stockholm format using AlignIO (Biopython; Cock et al., 2009) and then pHMM profiles were created using HMMbuild (HMMER 3.1b2; Eddy, 2011) with the option --singlemx, to enable profile building if only one sequence was given, and default parameters, except MAP (yes) and STATS (LOCAL: MSV / VITERBI / FORWARD).

The pHMMs were further curated by running HMMsearch (HMMER 3.1b2; Eddy, 2011) on all the proteins which had been used to create the pHMMs. The results of this search were used to guide the selection of the threshold values (Supplementary Figure 18) for grouping sequences into the “classified”, “ambiguously classified” and “unclassified” categories. A list of the pHMMs, number of input sequences, cropping coordinates, and the HMMbuild output information is provided in Supplementary Table 9, and the pHMMs and alignments with accession numbers are available in Supplementary Dataset 4 and at github.com/ingridole/ViralRdRp_pHMMs.

### Sequence databases for the RdRp search

To obtain viral reference sequences, we used the assembly_summary.txt file at ftp.ncbi.nih.gov/genomes/refseq/viral/ (10 May 2018). Viral sequences from this file with a complete genome and the latest version number were downloaded with wget from the NCBI path above. In total, there were 9,566 viral nucleotide reference sequences.

Non-redundant nucleotide (nr/nt) sequences were downloaded (14–15 May 2018) from the NCBI nucleotide database in 12 sets covering all groups of RNA viruses as well as unclassified and unassigned viruses. The taxonomy and a minimum sequence length of 1,000 nt were specified and two over-represented species (hepatitis C virus and influenza A virus) were excluded. The exact search queries are provided in Supplementary Table 10. In total, there were 274,579 nr/nt sequences. All sequences which were not defined as dsRNA, −ssRNA or +ssRNA viral sequences were combined into one file. Within that file, we removed sequences named as phages or where the molecule type was specified as genomic DNA in the GenBank format files, and sequences that were longer than 30,360 nucleotides (so as to remove many phage and DNA virus sequences). Following this, 7,059 sequences remained in the “others” file. Next, clustering within each of the four groups (dsRNA, −ssRNA, +ssRNA, others) was performed using CDHIT (versions 4.6 and 4.7; Li & Godzik, 2006; Fu et al., 2012) to remove similar sequences (>80% pairwise nucleotide identity), in each case retaining the longest sequence from a group of similar sequences. After this step, 14,832 sequences were left for further processing (Supplementary Table 11).

Next, we used BLASTN (v2.2.31+, built 7 Jan 2016; Altschul et al., 1990; Camacho et al., 2009) to compare the remaining 14,832 nr/nt sequences with the 9,566 reference sequences, and we removed nr/nt sequences that had ≥80% nucleotide identity to, and ≥80% coverage by at least one reference sequence. After this step, 9,855 nr/nt sequences were left and used in the further analysis.

Sequences from the NCBI TSA database (National Center for Biotechnology Information (NCBI), 2017) were downloaded (29 Oct 2017) based on the wgs_selector file from the TSA browser at https://www.ncbi.nlm.nih.gov/Traces/wgs/?view=TSA. We downloaded 2,648 different TSA datasets covering ∼1,800 unique taxonomic groups.

The exact commands for downloading and clustering of sequences are provided in Supplementary Methods.

### Analysis of protein sequences

For all TSA, reference and nr/nt nucleotide sequences, open reading frames (ORFs) were retrieved using GETORF (EMBOSS v5.5 and v6.6; Rice et al., 2000) using three different genetic code tables (the standard genetic code, table = 1; stop codon UGA redefined as Trp, table = 4; and stop codons UAA and UAG redefined as Gln, table = 6), and identifying regions between consecutive stop codons (parameter -find 0) with a minimum length of 60 nucleotides (parameter -minsize 60) to allow for detection even of RdRp fragments. In total, this resulted in >13 × 10^9^ ORFs (Supplementary Table 12).

### HMMsearch

To search for viral RdRps, we searched the retrieved ORFs using HMMsearch (HMMER 3.1b2; Eddy, 2011) with the 77 family-level pHMM profiles. HMMsearch was performed for each genetic code table dataset separately, adjusting the E-value based on database size to maintain a constant *p*-value threshold of 1 × 10^−6^. The next step was to find the best hit for each ORF among the different matched pHMMs and sort the match to one of three groups: classified, ambiguously classified or unclassified based on our IDscore metric, which is the bit score divided by the length in amino acids of the alignment between an ORF and a matched pHMM (Supplementary Figure 18). If the highest IDscore was lower than 0.25, a sequence was sorted into the unclassified group. Otherwise, if a sequence had statistically significant hits to more than one of the pHMMs and the second best IDscore was less than 20% lower than the best IDscore, the sequence was sorted into the ambiguously classified group. Sequences with an IDscore of 0.25 or higher, and at least 20% difference in IDscore between the first and second best hits, were sorted into the classified group and classified according to the best match pHMM.

### Processing of RdRp-encoding ORFs

Since there could be multiple partially identical ORFs due to the use of three genetic code tables, we next used pairwise global alignments (Biopython pairwise2.align.globalxx; Cock et al., 2009) to compare all ORF sequences with the same original nucleotide accession. If, in any pairwise alignment, the number of identities divided by the shorter ORF length was 1, then the shorter ORF was removed from further analysis.

The remaining ORFs were trimmed according to the start and end positions of the amino acid region which mapped to the best hit pHMM to approximate the core part of the RdRp. Then alignments were generated for each pHMM group, combining reference, nr/nt and TSA-derived trimmed ORF sequences, using MUSCLE v3.8.31 (Edgar, 2004).

### Grouping into 60 clusters

After applying the grouping scheme and discarding similar sequences using CD-HIT-EST, for some of the 77 viral pHMM-based groups there were fewer than 10 classified sequences remaining. For convenience, these sequences were joined with other groups which were somewhat taxonomically similar, with preference given to those groups which themselves had fewer sequences, aiming for each cluster to contain close to or more than 20 sequences. Thus, 60 clusters of classified sequences were created. Clustering was not performed for the ambiguously classified or unclassified groups of sequences.

### Verification

In order to verify that sequences were true viral RdRps rather than false positives, and to identify chimeric sequences, all ORFs identified as encoding (part of) an RdRp were checked using a BLAST-based approach. For verification with a third approach, HHSearch (part of HHSuite v3.3.0; Steinegger et al., 2019) was used to compare our set of putative viral ORFs to the Pfam database (Finn et al., 2014) We compared the false positive rate for HMMER and BLASTP using a control set of Uniprot proteins. All verification steps are described in full in the Supplementary Materials and Methods. Full results are provided in Supplementary Dataset 2.

### Heatmap for relationships between viral groups

For all ORFs in the classified group, we identified the first and second best hit pHMMs based on IDscore. Then, for each represented pHMM group, we counted the number of co-occurrences with each of the pHMMs. This resulted in a large matrix, where rows represent the first best hit pHMM and columns the second best hit pHMM. Then, we normalised the values by row. Results were visualised in Python (Matplotlib, pyplot and colors, and Numpy packages). The virus taxonomic groups of the pHMMs were manually sorted in an order consistent with ICTV taxonomy and/or the phylogeny of Wolf et al. (2018).

### Phylogenetic trees

Phylogenetic trees were constructed using amino acid alignments and PhyML v3.1 (amino acid model LG, tree topology search operation SPR, and gamma distribution shape parameter 20; Guindon & Gascuel, 2003; Guindon et al., 2010). In some cases, similar sequences were discarded using pairwise comparisons in Python or using CDHIT. Alignments were prepared with MUSCLE v3.8.31 (Edgar, 2004) and their format changed to Phylip using SEQRET (EMBOSS v5.5 and v6.6; Rice et al., 2000). FigTree v1.4.3 (Rambaut, 2006) was used for visualisation of trees using the midpoint rooting option and showing the bootstrap support values as node labels.

### Host identification for TSA sequences

To identify the putative host species and taxonomic group for RdRp sequences found in the TSA database, we used EFETCH (Entrez Programming Utilities; Sayers et al., 2022) and each TSA contig accession number to retreive a corresponding taxonomy line. Next, we assigned the type of putative host as follows. If “Eukaryota” was not in the taxonomy line the type was set to metagenomics (or environmental). In other cases, we searched for keywords in the taxonomy line: fungi (keyword “Fungi”), plants (keyword “Viridiplantae”), vertebrates (keyword “Vertebrata”), arthropods (keyword “Arthropoda”). For invertebrates, we required the keyword “Metazoa” but not “Vertebrata” or “Arthropoda”, while for Protista the keywords “Metazoa”, “Fungi” and “Viridiplantae” had to be absent.

To generate the updated taxonomy column in Supplementary Dataset 1, corresponding to the most recent (28 June 2022) NCBI taxonomy, derived from the ICTV 2021 taxonomy (Schoch et al., 2020; Walker et al., 2021), the taxonomic classification assigned to the GenBank record for each accession number was used, again retrieved using EFETCH. Where this ID corresponded to the host rather than the virus, it was not included. To assign a taxonomic lineage to the pHMM family/genus level classifications, the name of the pHMM was used except in the cases of the *Sobemoviridae* (which is now the genus *Sobemovirus*), *Ophioviridae* (which is now renamed as *Aspaviridae*) and *Rubellavirus* (which is now *Rubivirus*). *Zhaovirus*, *Yanvirus* and *Weivirus* are not classified in the current ICTV taxonomy.

### Data Availability

The table of identified virus RdRp derived sequences (Supplementary Dataset 1), results of our BLAST and HHSearch verification (Supplementary Dataset 2) and PhyML trees of the different groups of sequences (Supplementary Dataset 3) are available at github.com/ingridole/ViralRdRp_pHMMs_2, release v1.0.1 is associated with this manuscript. The pHMMs and the alignments used to create them (Supplementary Dataset 4) are available at github.com/ingridole/ViralRdRp_pHMMs, release v1.0.1 is associated with this manuscript.

## Acknowledgements

This work was funded by Wellcome Trust Senior Research Fellowships [106207/Z/14/Z, 220814/Z/20/Z] and a European Research Council grant [646891]. For the purpose of open access, the author has applied a CC BY public copyright licence to any Author Accepted Manuscript version arising from this submission.

---

**Supplementary Figure 1.**
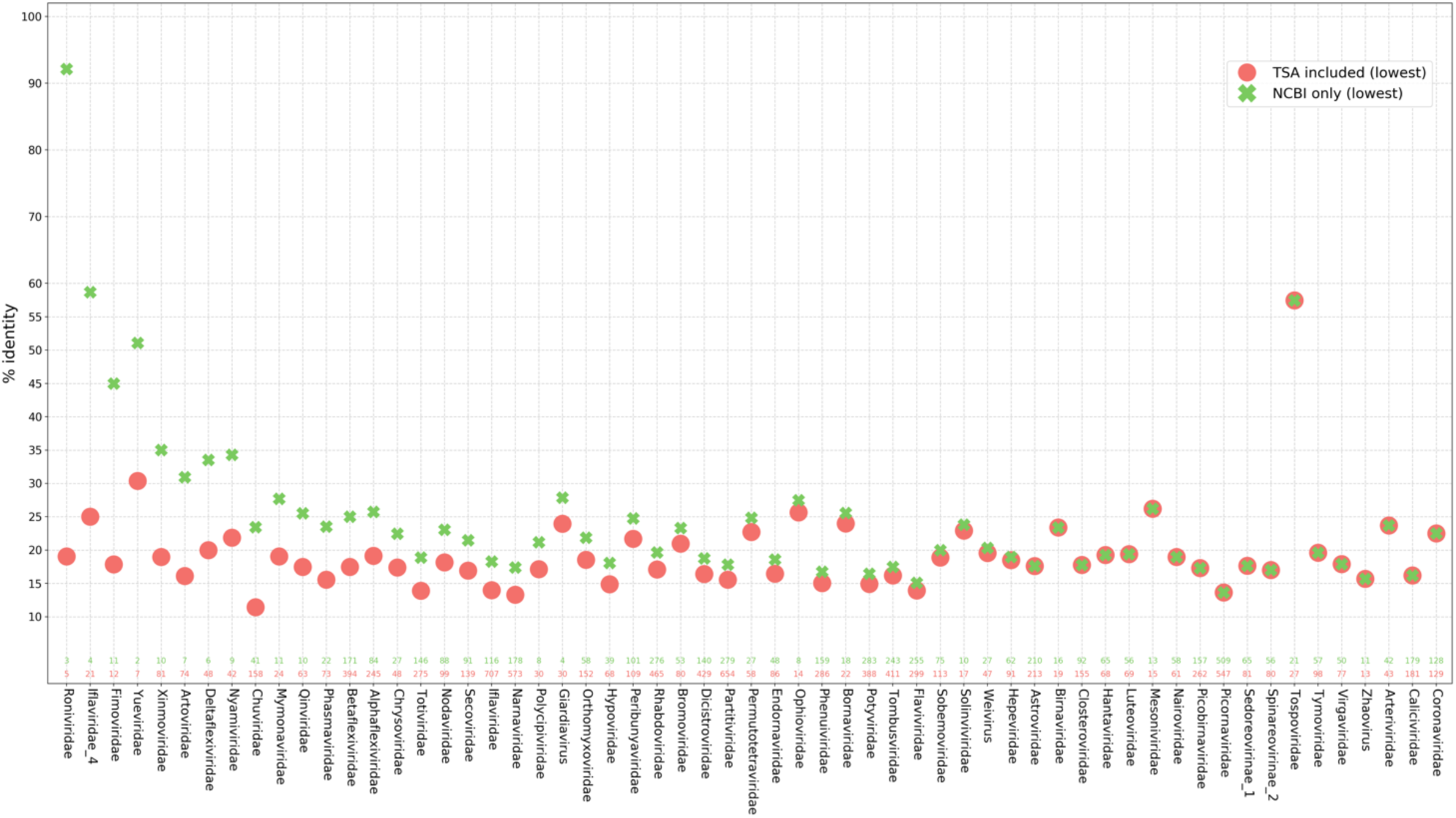
Lowest pairwise identities among sequences classified to each pHMM group. All-against-all pairwise identities were calculated with BLASTP (Altschul et al., 1990; Camacho et al., 2009). For each pHMM family, the lowest pairwise identity scores for NCBI sequences (green crosses) and for NCBI and TSA sequences combined (coral circles) are shown. Families were plotted only if after discarding 100%-identical trimmed RdRp sequences there was >1 NCBI sequence left. If there was no TSA sequence left, the circle was not plotted. Numbers (using the same colours) at the bottom of the graph show the numbers of unique sequences used in the BLASTP analysis.

**Supplementary Figure 2.**
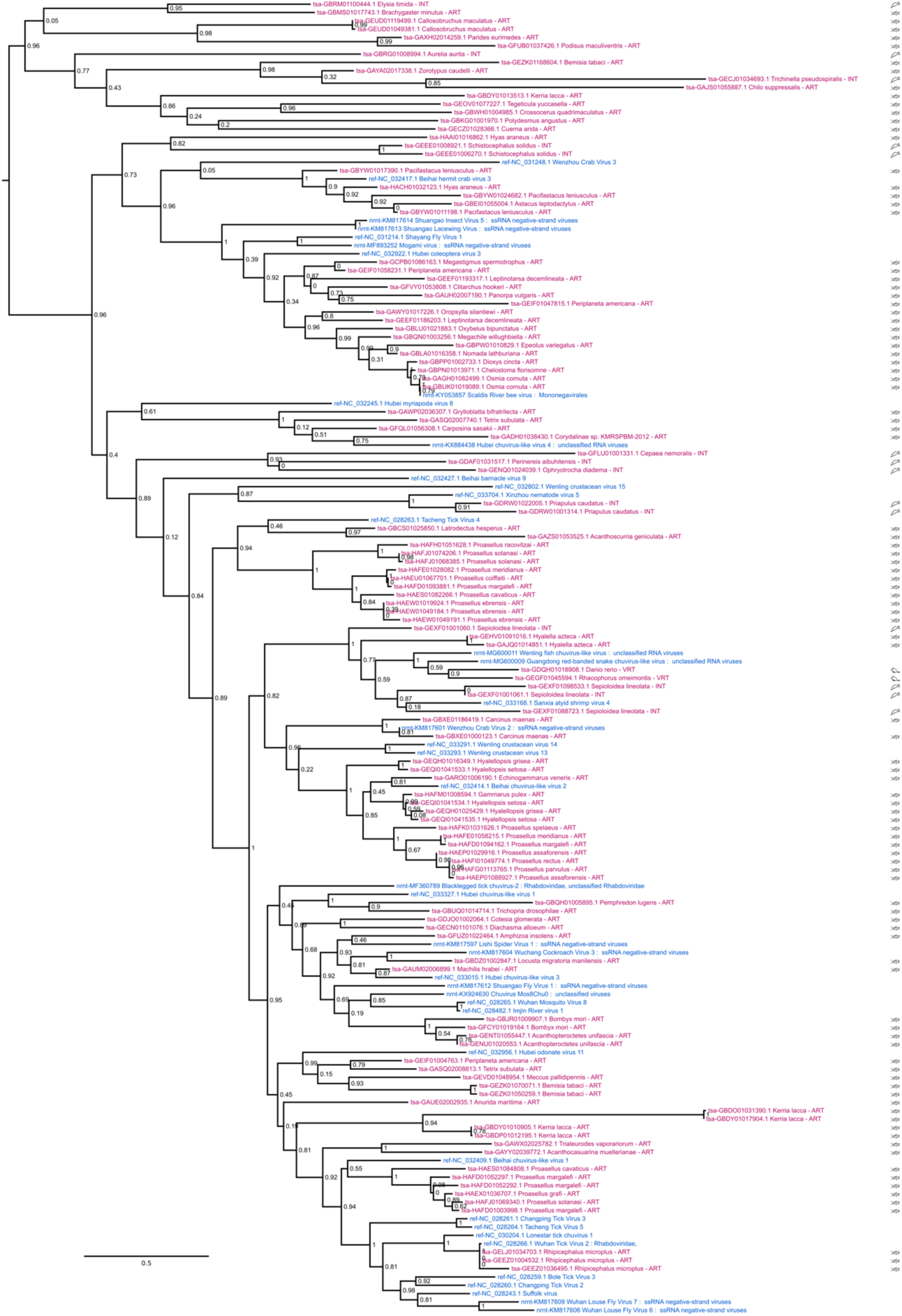
Phylogenetic tree for the core RdRp sequences of chu-like viruses. Duplicate 100%-identical RdRp core sequences were removed. Blue – nr/nt sequences; pink – TSA sequences. For TSA sequences, a category icon for the putative host (i.e. the TSA target organism) is shown at right.

**Supplementary Figure 3.**
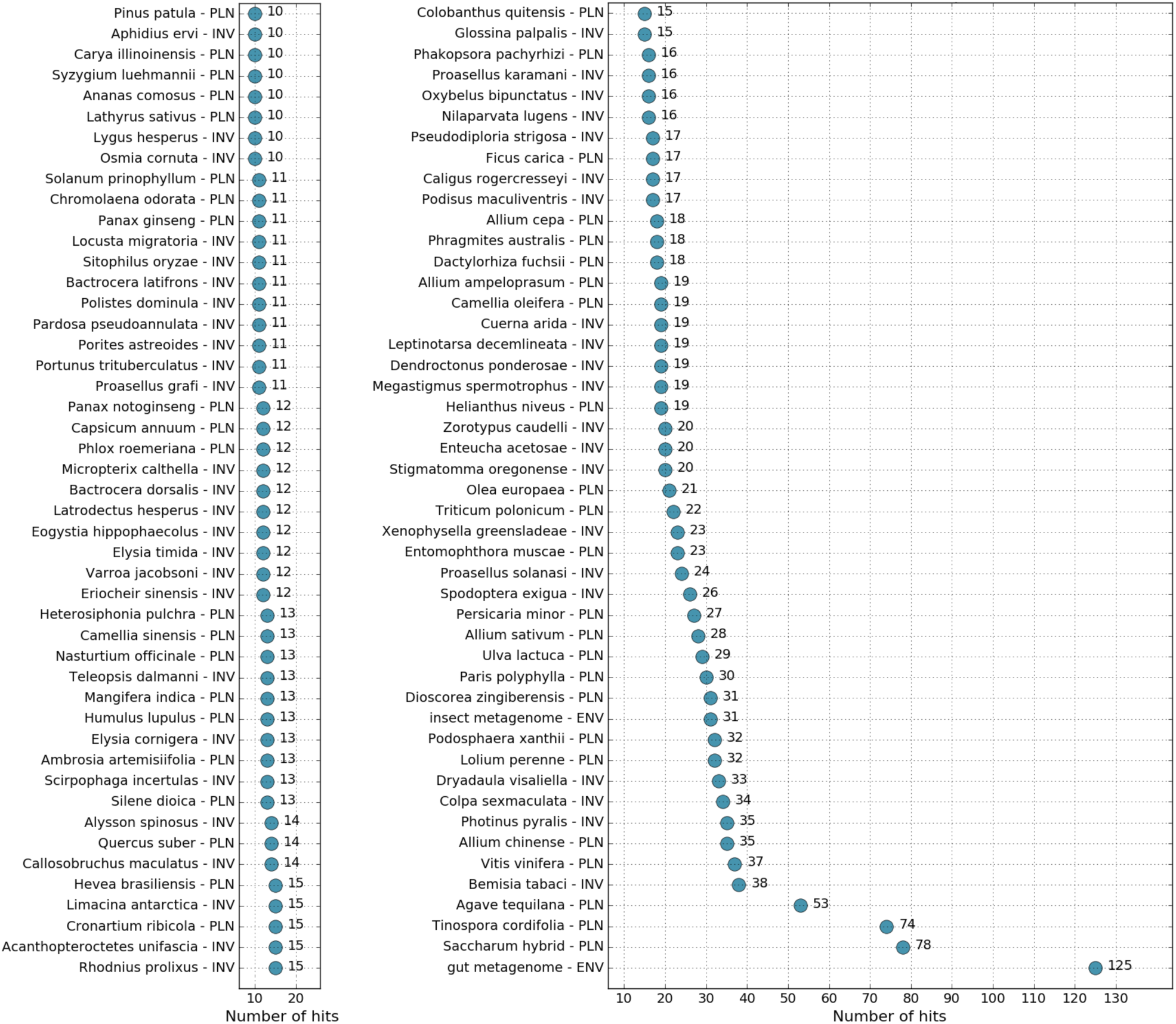
Host species with highest numbers of non-identical RdRp sequences identified in TSA datasets. PLN – plant, INV – invertebrate, ENV – environmental sample.

**Supplementary Figure 4.**
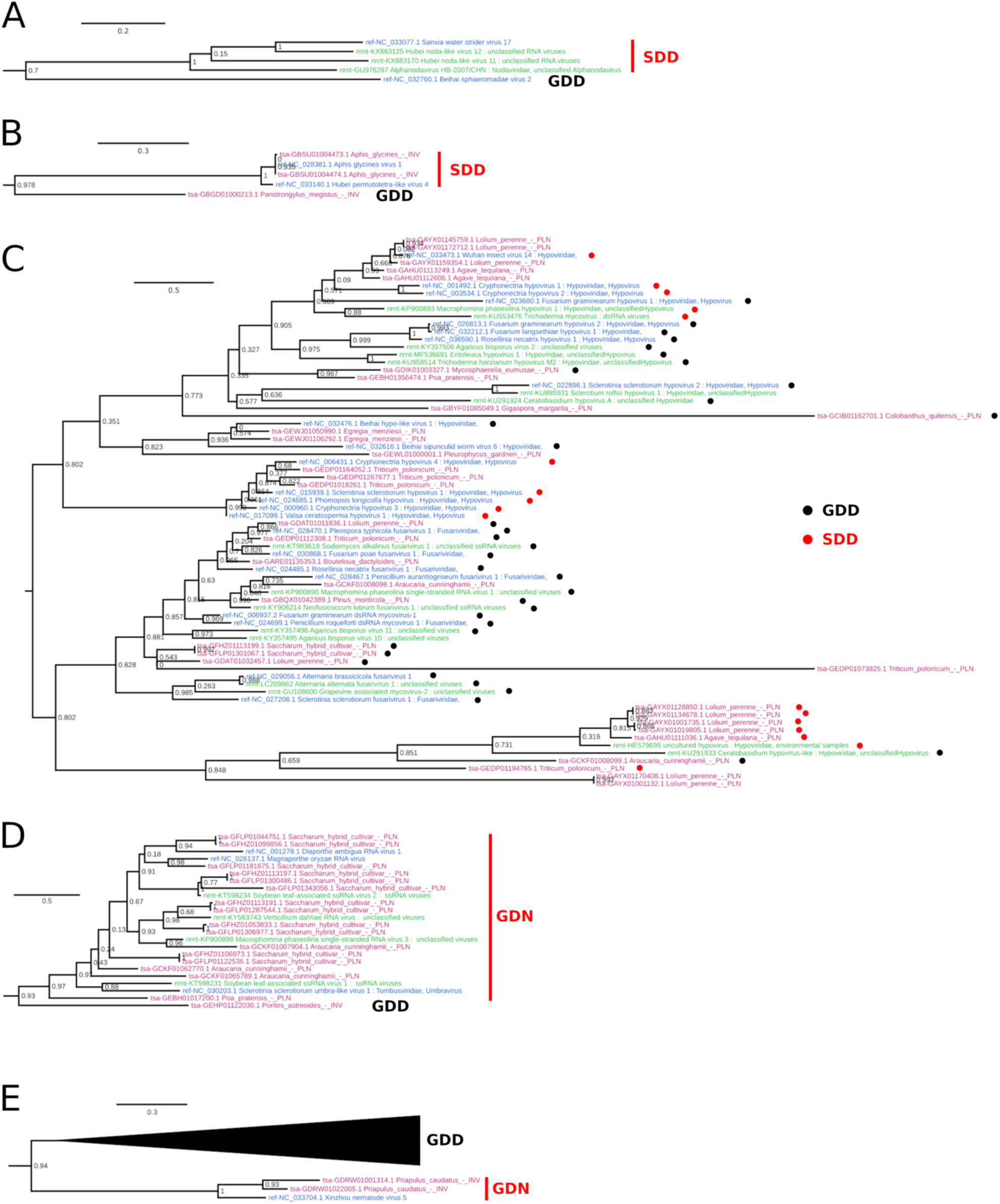
Phylogenetic trees highlighting subclades of sequences where a typical GDD motif C has been replaced by an alternative sequence. **(A)** A clade of noda-like viruses with SDD. **(B)** A clade of permutotetra-like viruses with SDD. **(C)** Hypoviruses commonly have SDD but SDD-containing viruses appear to be dispersed across the hypo-like virus phylogeny. **(D)** A clade of tombus-like viruses with GDN. **(E)** A clade of chu-like viruses with GDN. See clusters 29, 42, 39, 8 and 15, respectively, in Supplementary Dataset 3 for the complete trees of each cluster from which these subtrees were extracted.

**Supplementary Figure 5.**
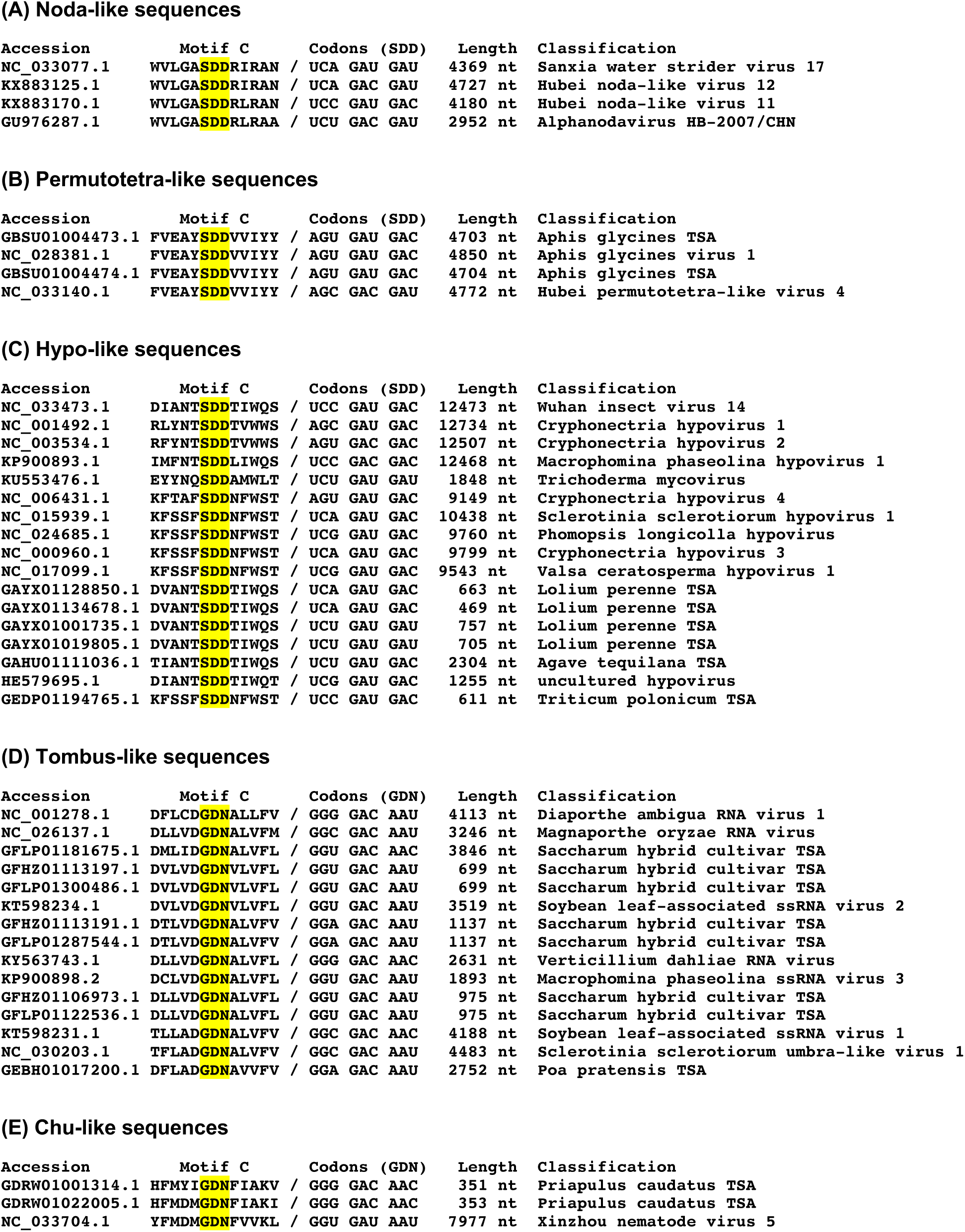
Motif C amino acid and nucleotide sequences for subclades of sequences where a typical GDD motif C has been replaced by an alternative sequence. **(A)** A clade of noda-like viruses with SDD. **(B)** A clade of permutotetra-like viruses with SDD. **(C)** Hypoviruses containing SDD. **(D)** A clade of tombus-like viruses with GDN. **(E)** A clade of chu-like viruses with GDN.

**Supplementary Figure 6.**
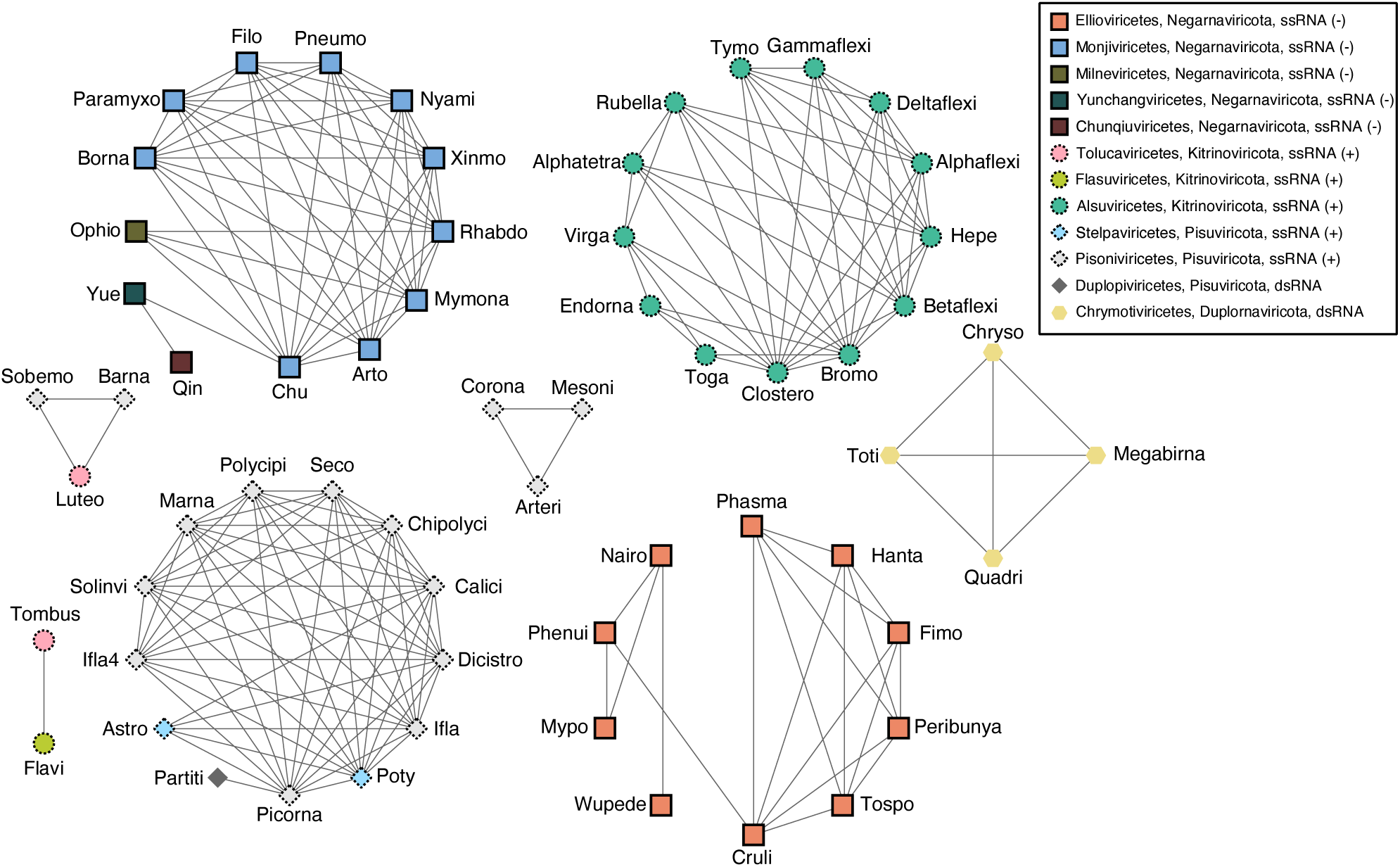
Undirected network diagram linking the HMM models which had the highest and second highest scores against RdRp ORFs identified with our pipeline. All classified-group ref, nr/nt and TSA RdRp ORFs were used (Supplementary Dataset 1). Nodes with no edges are not shown. Colours represent class, shapes represent phylum and outline styles represent Baltimore classification (+ssRNA, −ssRNA or dsRNA) as indicated in the legend. Networks were created with the Python networkx package (version 2.5.1, Hagberg et al., 2008) and visualised using Cytoscape (version 3.6.1, Shannon et al., 2003) with “degree sorted circle” layout.

**Supplementary Figure 7.**
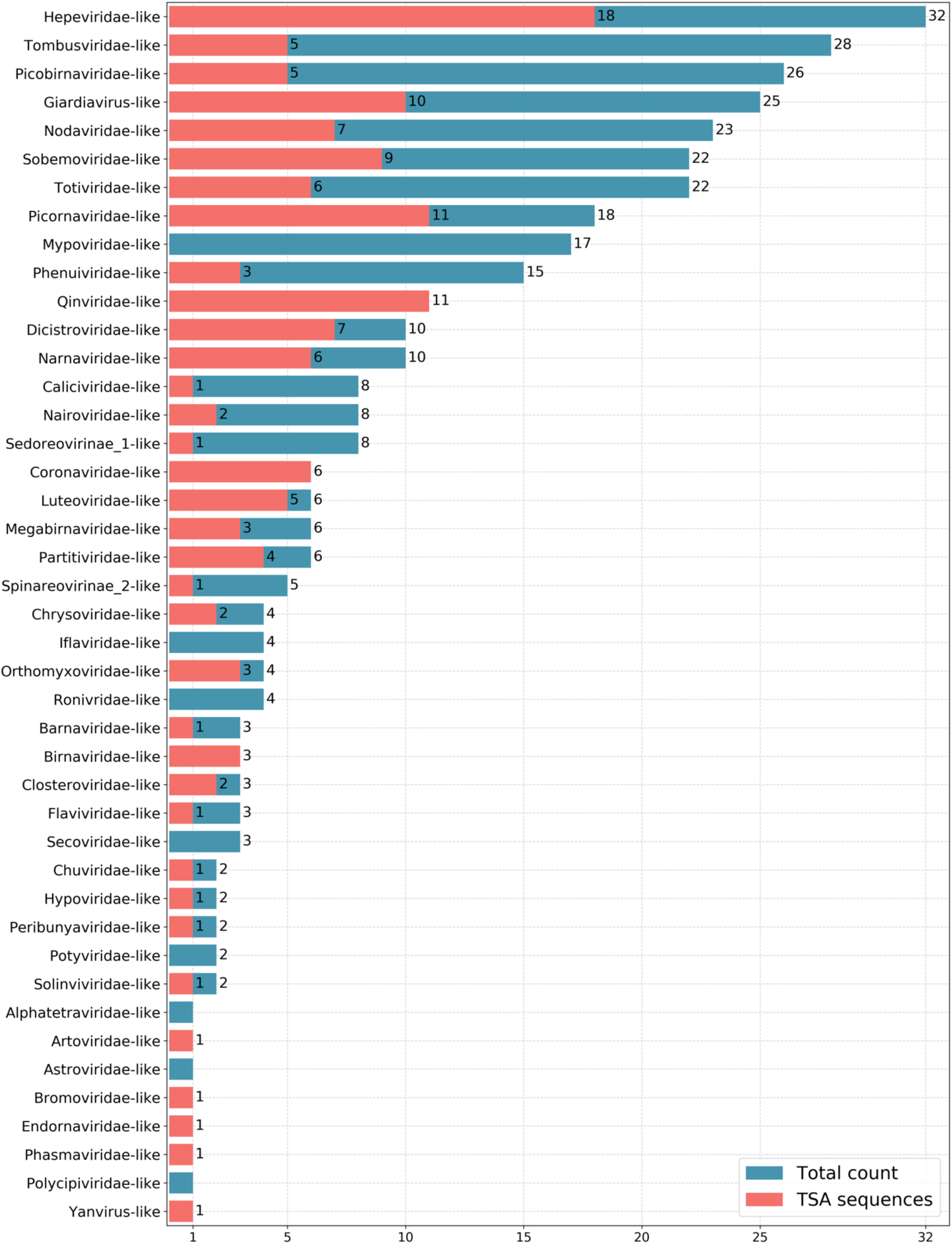
Numbers of unclassified sequences. Total count (TSA, ref and nr/nt) – blue; TSA only – red. The unclassified sequences were sorted according to the best-match pHMM virus group (43 out of 77 pHMM groups being represented).

**Supplementary Figure 8.**
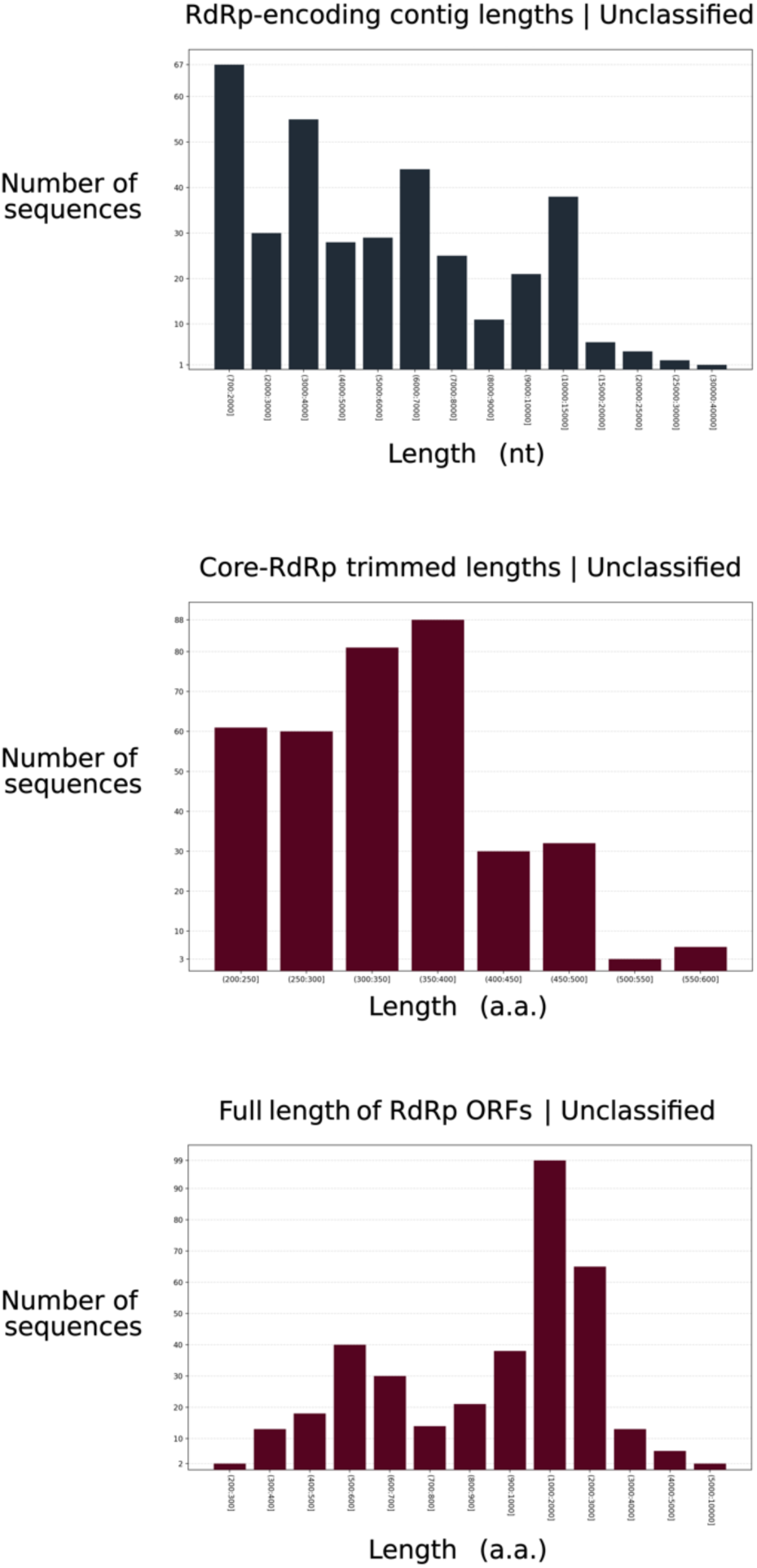
Histograms of lengths of the RdRp-encoding contigs, their full RdRp-encoding ORFs, and the RdRp ORFs when trimmed to the RdRp core, for the “unclassified” group.

**Supplementary Figure 9.**
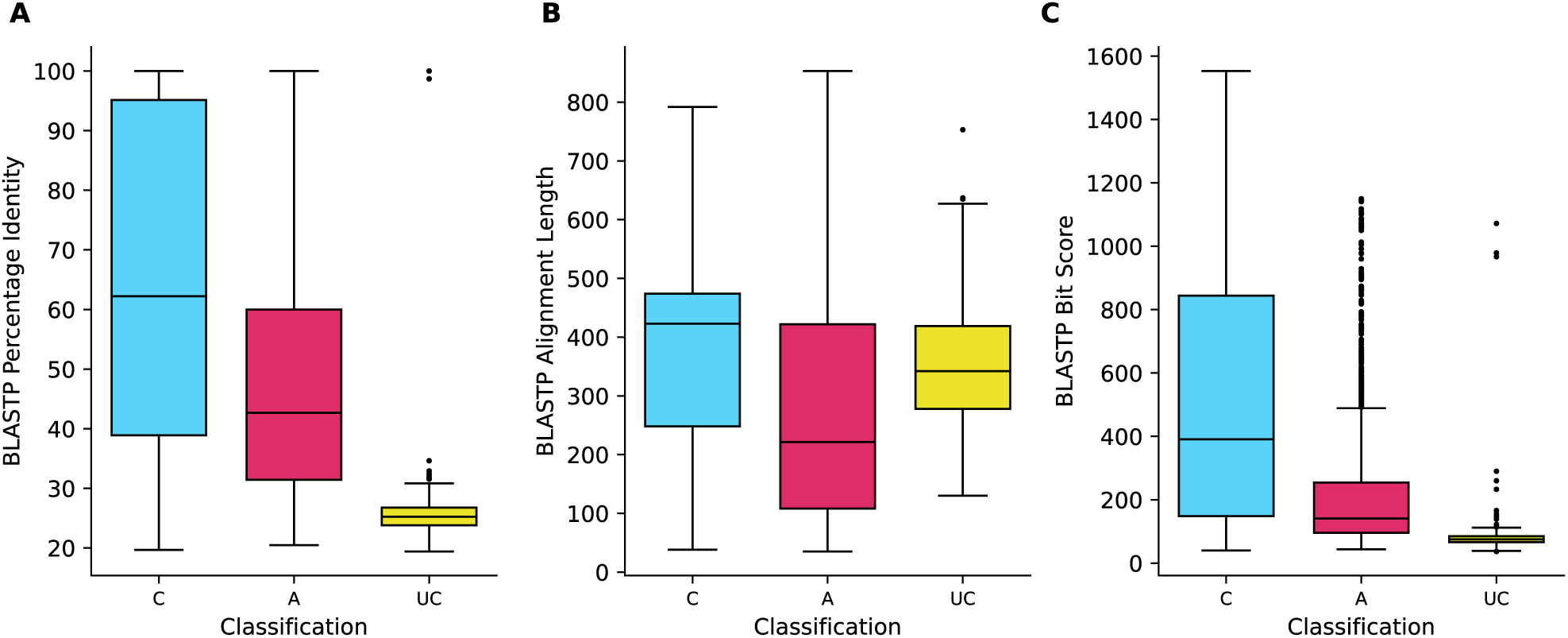
Boxplots showing the results of a BLASTP analysis comparing each RdRp ORF identified here to the most similar sequence included in the input pHMMs, excluding self matches for the Genbank sequences, by percentage identity **(A)**, alignment length **(B)** and bit score **(C)**. Sequences are categorized as classified (box labelled C), ambiguous (A) or unclassified (U) based on the IDscore (HMMER bit score / HMMER alignment length) of the sequence against the best and second best matched pHMM.

**Supplementary Figure 10.**
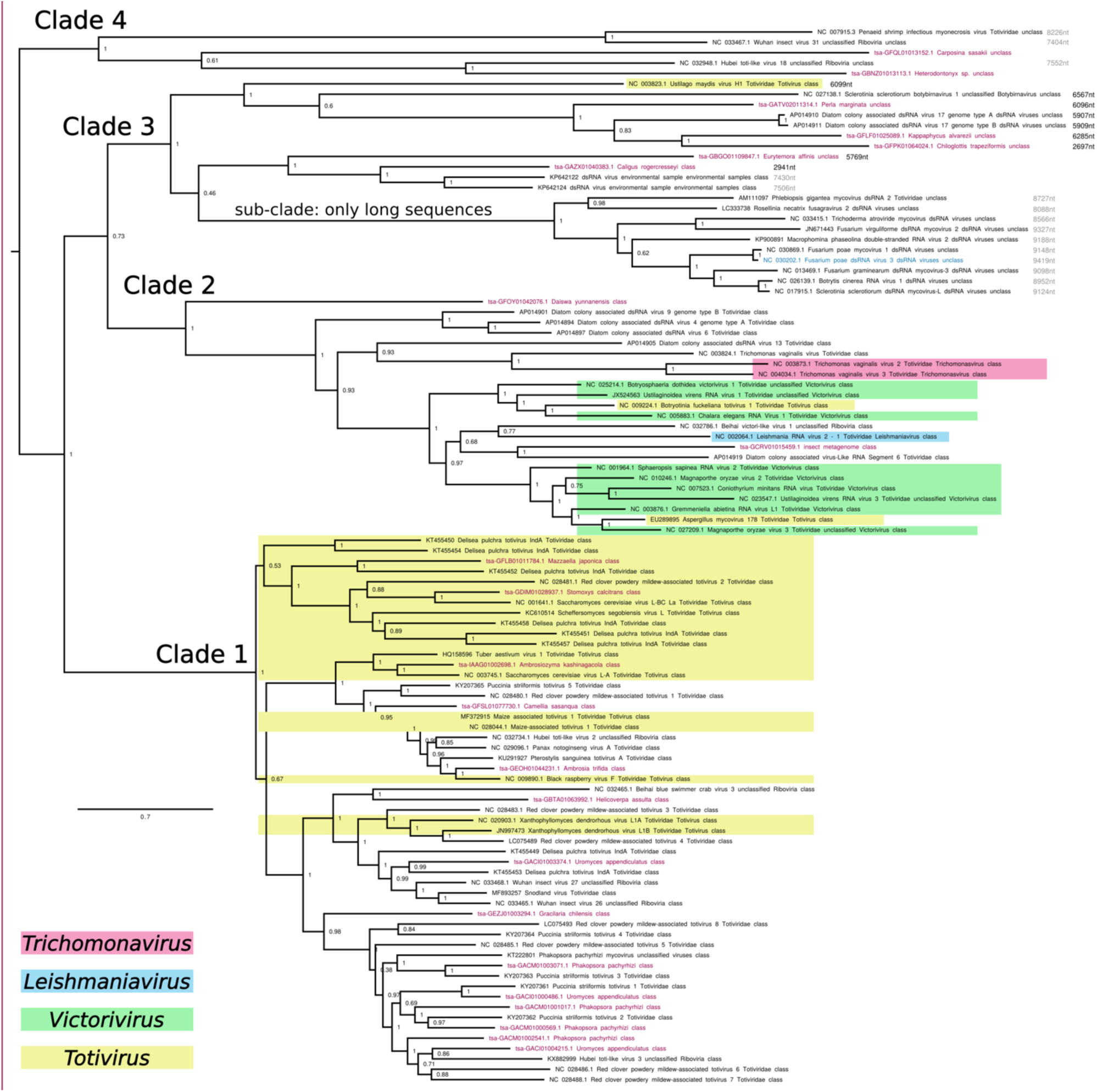
PhyML (Guindon & Gascuel, 2003; Guindon et al., 2010) tree based on the core RdRp region of 80 representative sequences classified to the *Totiviridae* pHMM, besides 22 unclassified sequences with best match to the *Totiviridae* pHMM. Four major clades are annotated (see main text). A subclade containing unusually long sequences (corresponding to a previously proposed family, Fusagraviridae) is also annotated.

**Supplementary Figure 11.**
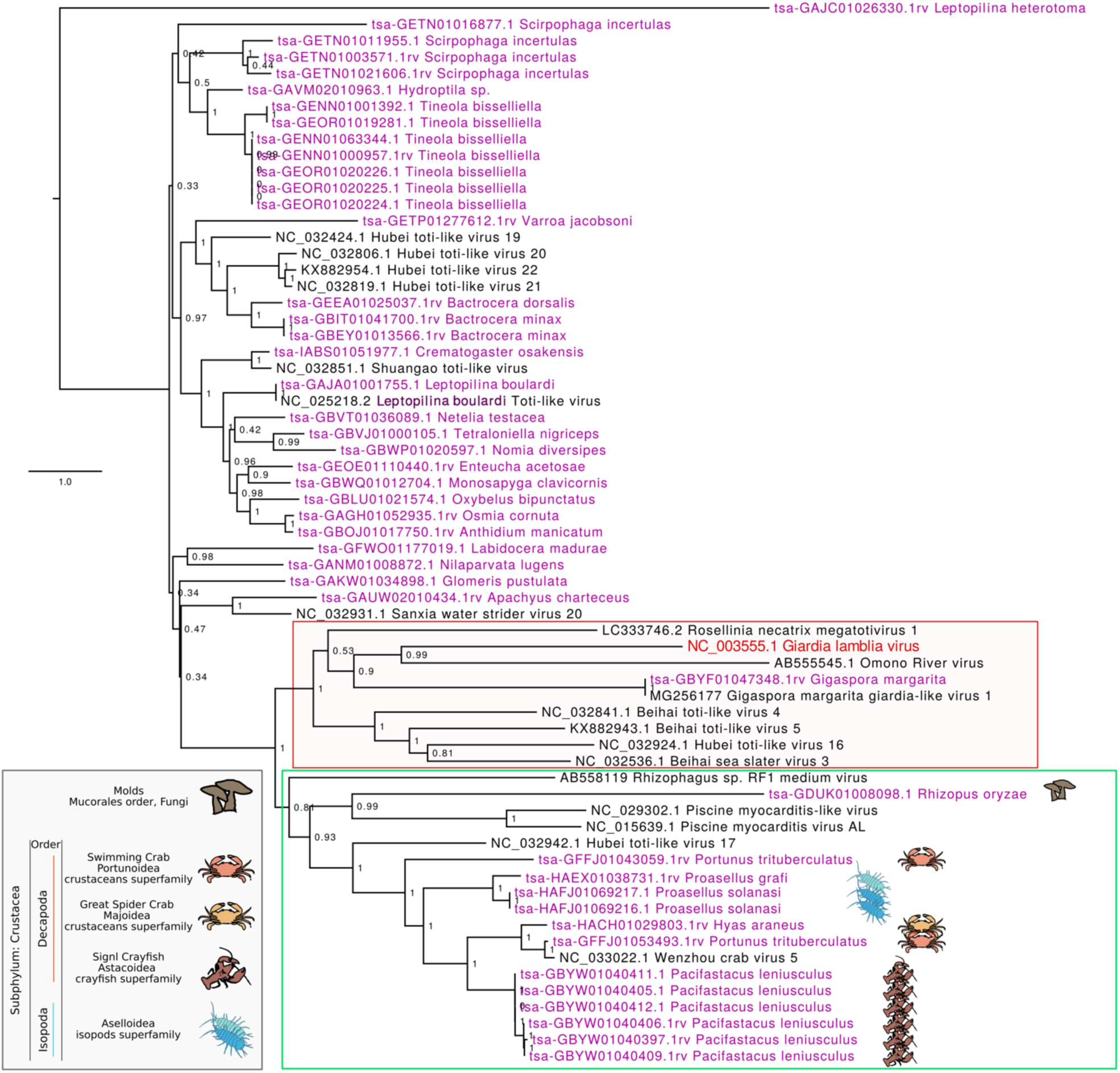
PhyML tree based on the core RdRp region of sequences classified to the *Giardiavirus* pHMM, besides unclassified sequences with best match to the *Giardiavirus* pHMM. Red box – Giardia lamblia virus clade. Green box – clade of mainly crustacean-infecting giardia-like viruses. Note that the outlying position of GAJC01026330.1 is a result of it containing a fragmented RdRp ORF (a likely EVE sequence).

**Supplementary Figure 12.**
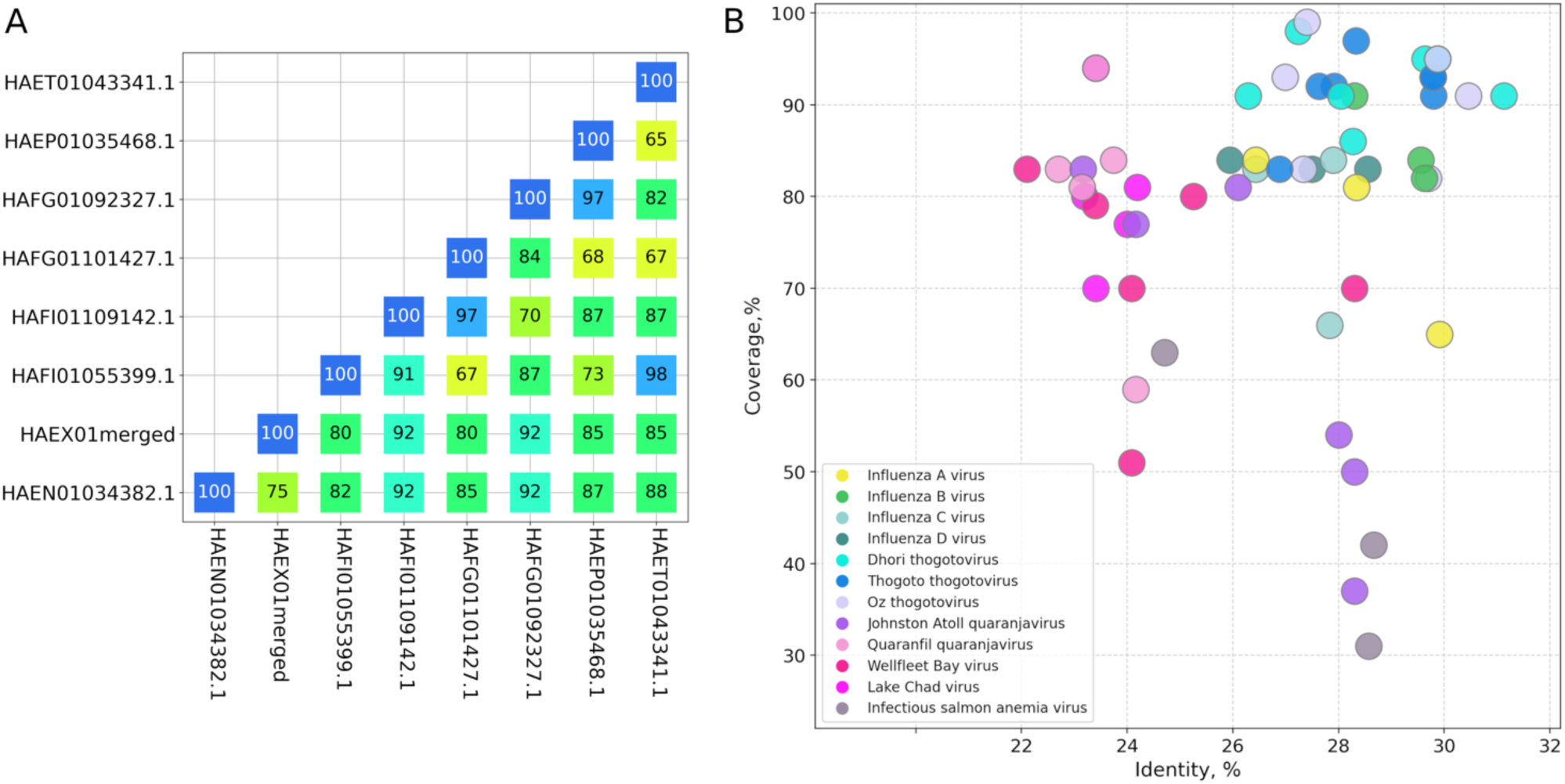
Sequences in the Asellidae-associated clade of orthomyxovirus-like sequences. **(A)** Pairwise nucleotide identities for Asellidae-associated PB1-encoding contigs. Identities were calculated with Biopython pairwise2 (globalms, match score 1, nonidentical score 0, opening gap −0.1, extending gap −0.01) (Cock et al., 2009). Alignment identity scores were normalized by the length of the shorter sequence in each sequence pair. **(B)**. Coverage and identity values for BLASTX comparisons between Asellidae-associated PB1-encoding contigs and various *Orthomyxoviridae* PB1 sequences (see Supplementary Table 3 for BLASTX E-values).

**Supplementary Figure 13.**
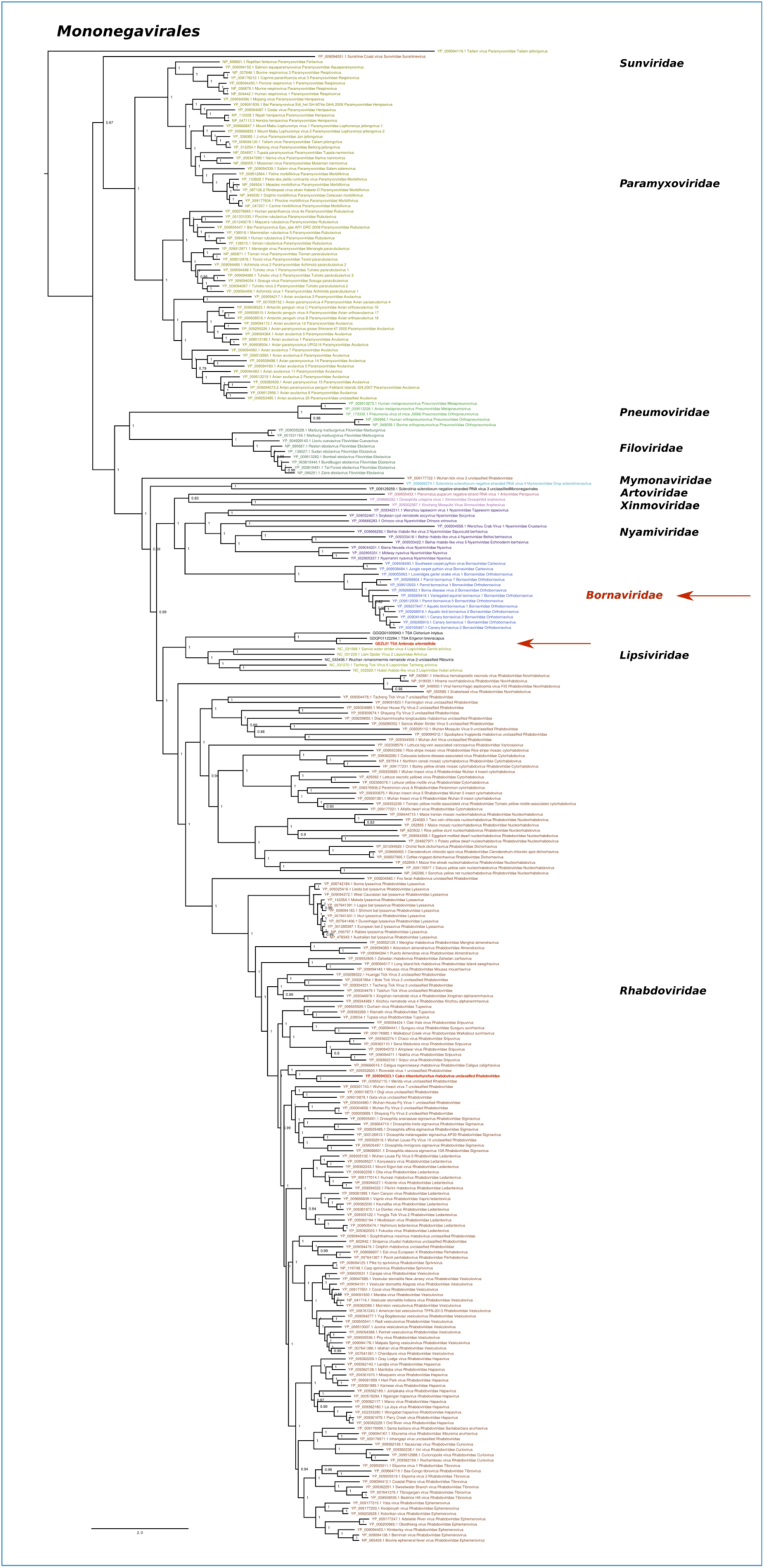
Phylogenetic tree of order *Mononegavirales* L protein sequences *s*howing the placement of the GEZL01-derived rhabdo-like virus (red arrow). NCBI RefSeq *Mononegavirales* L protein sequences were downloaded and similar sequences were discarded using CD-HIT (Li & Godzik, 2006; Fu et al., 2012) with a 90% identity threshold (cdhit -c 0.9), keeping the longest sequence in each cluster. These sequences were supplemented with the GEZL01-derived rhabdo-like virus and two additional TSA sequences identified by applying TBLASTX to the GEZL01-derived sequence, querying NCBI nr/nt sequences as well as *Asteraceae* TSA datasets. L protein sequences were aligned with MUSCLE (Edgar, 2004), and a phylogenetic tree was inferred using PhyML with default settings. Members of different *Mononegavirales* families (labelled at right) are indicated in different colours. Sequences/groups with known splicing are indicated in red.

**Supplementary Figure 14.**
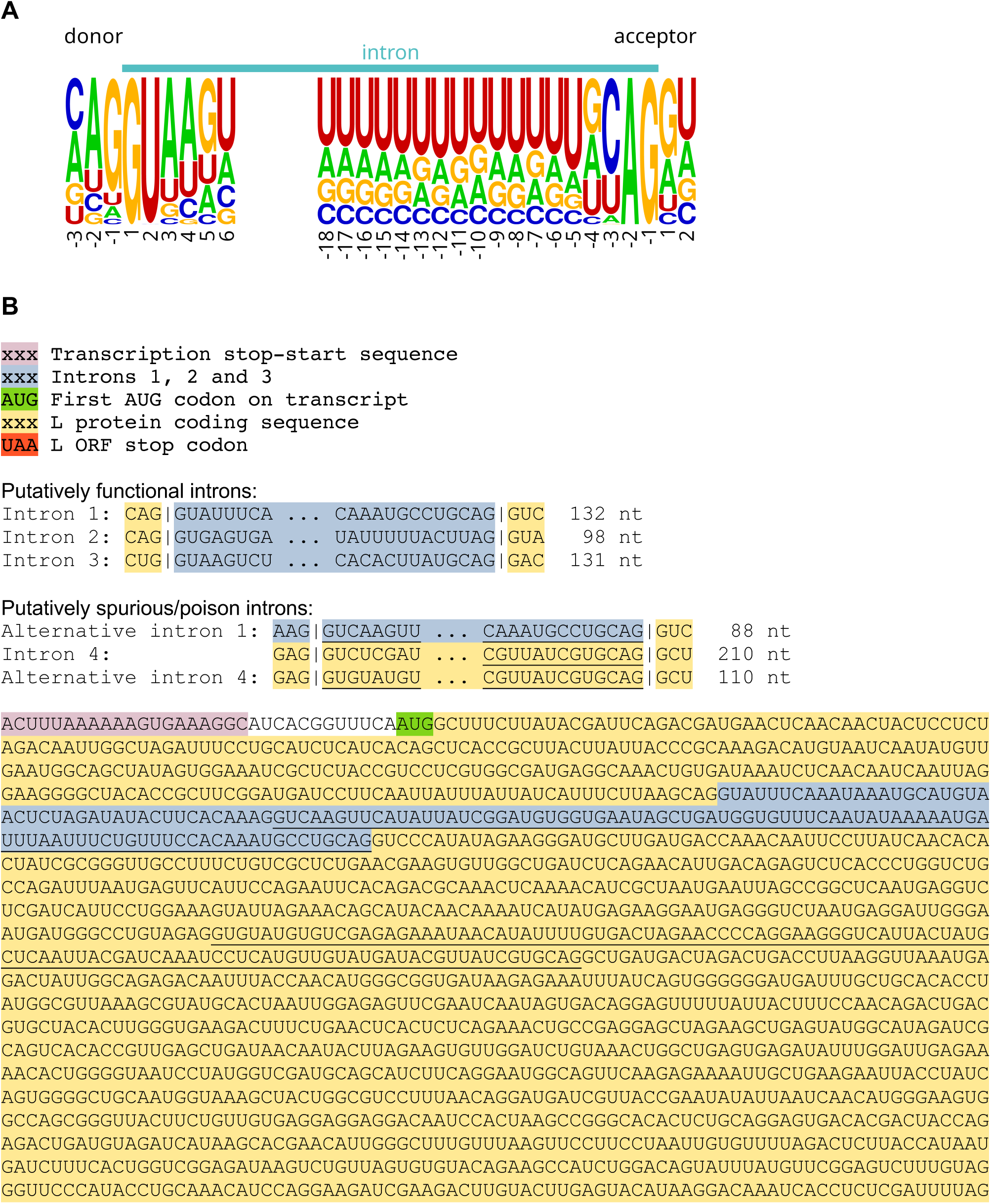
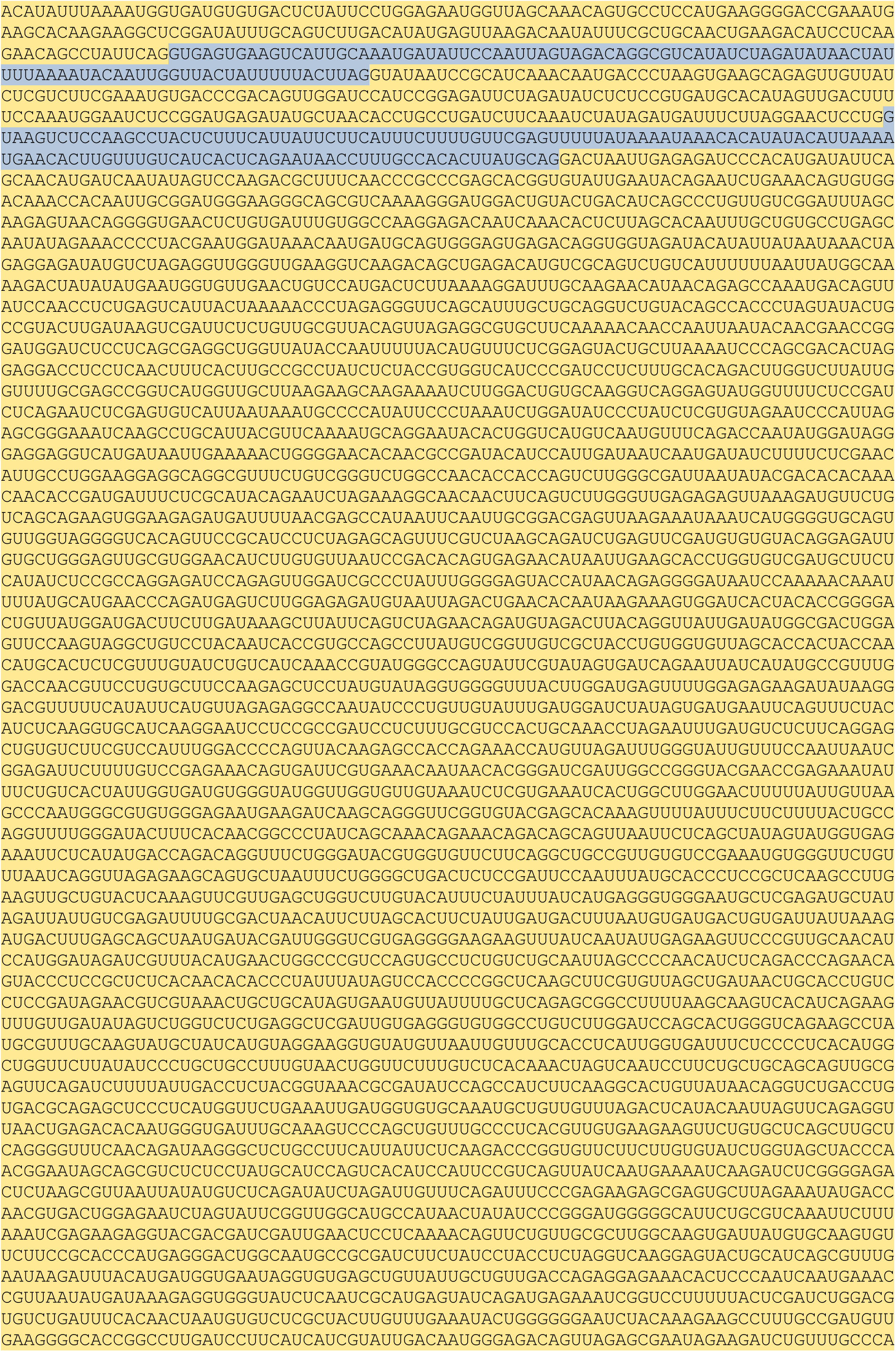
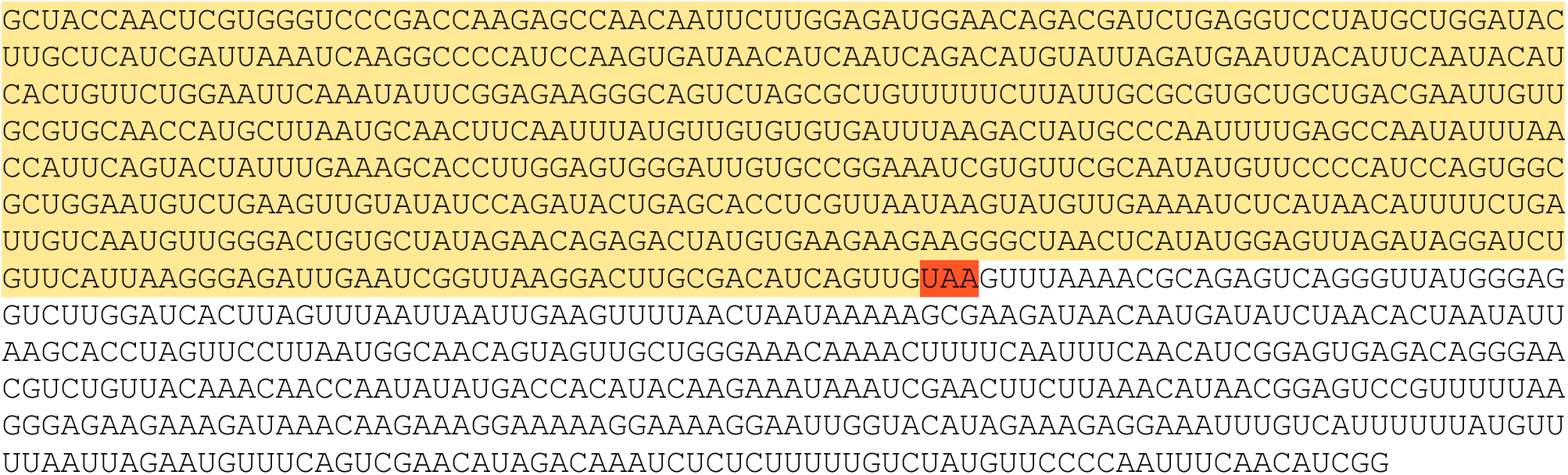
Splicing in the GEZL01-derived sequence. **(A)** Relative nucleotide frequencies from 998 *Arabidopsis thaliana* splice sites (data from Brown et al., 1996); sequence logo produced with WebLogo (Crooks et al., 2004). **(B)** Section of the GEZL01-derived sequence from the third transcription stop-start sequence to the end of the merged contig. The locations of the putative L ORF and introns supported by Hisat2 spliced read mapping are annotated. Removal of introns 1–3 (blue highlight) is required to express an intact L protein (yellow highlight). Splicing of the alternative introns 1 or 4 (underlined) disrupts the L protein ORF leading to truncation, whereas splicing of intron 4 (red) removes 70 a.a. from the L protein.

**Supplementary Figure 15.**
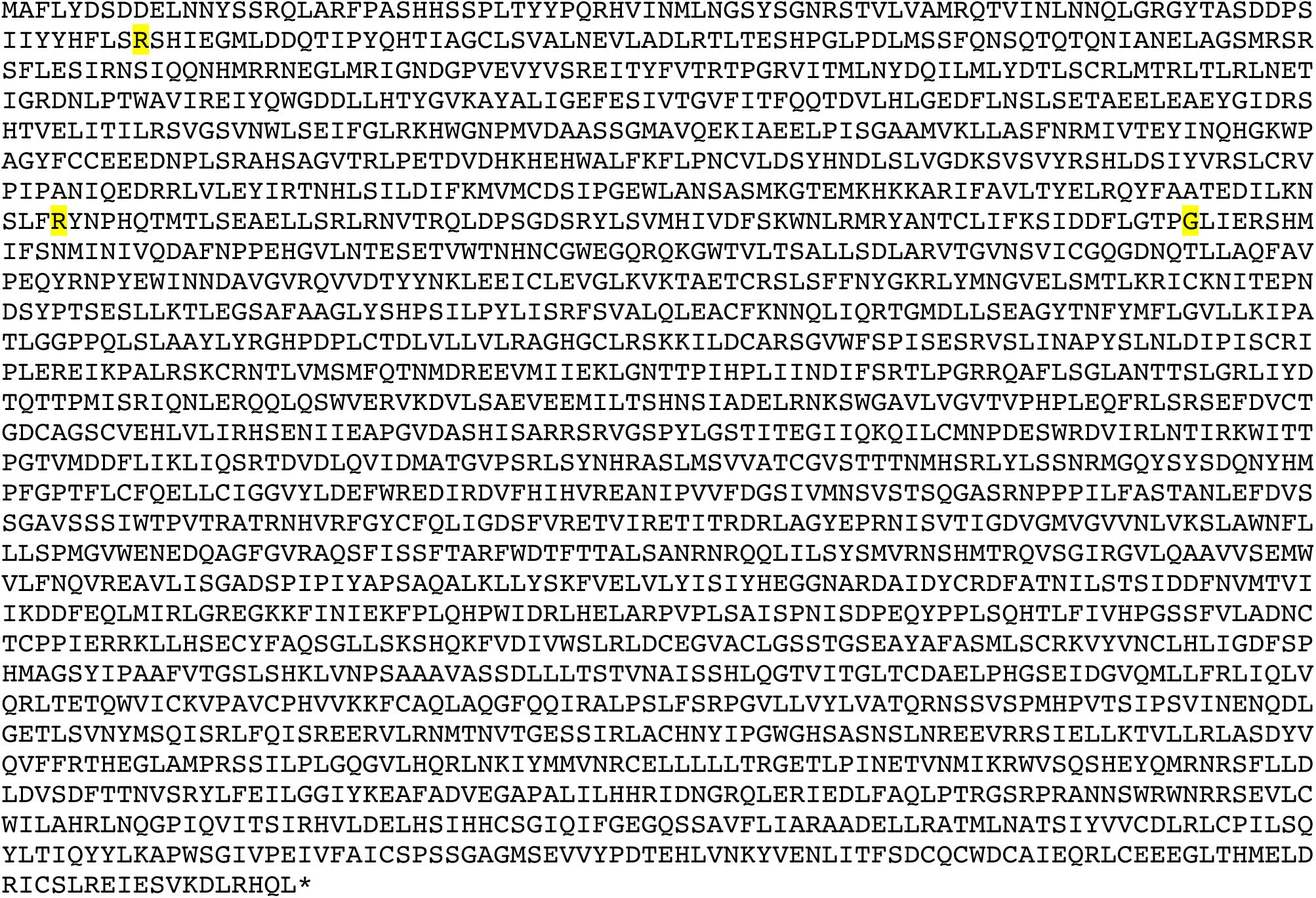
Translation of the predicted L ORF of the GEZL01-derived sequence. Only introns 1–3 were removed. The three amino acids that are encoded by a codon spanning one of the exon-exon junctions are highlighted in yellow.

**Supplementary Figure 16.**
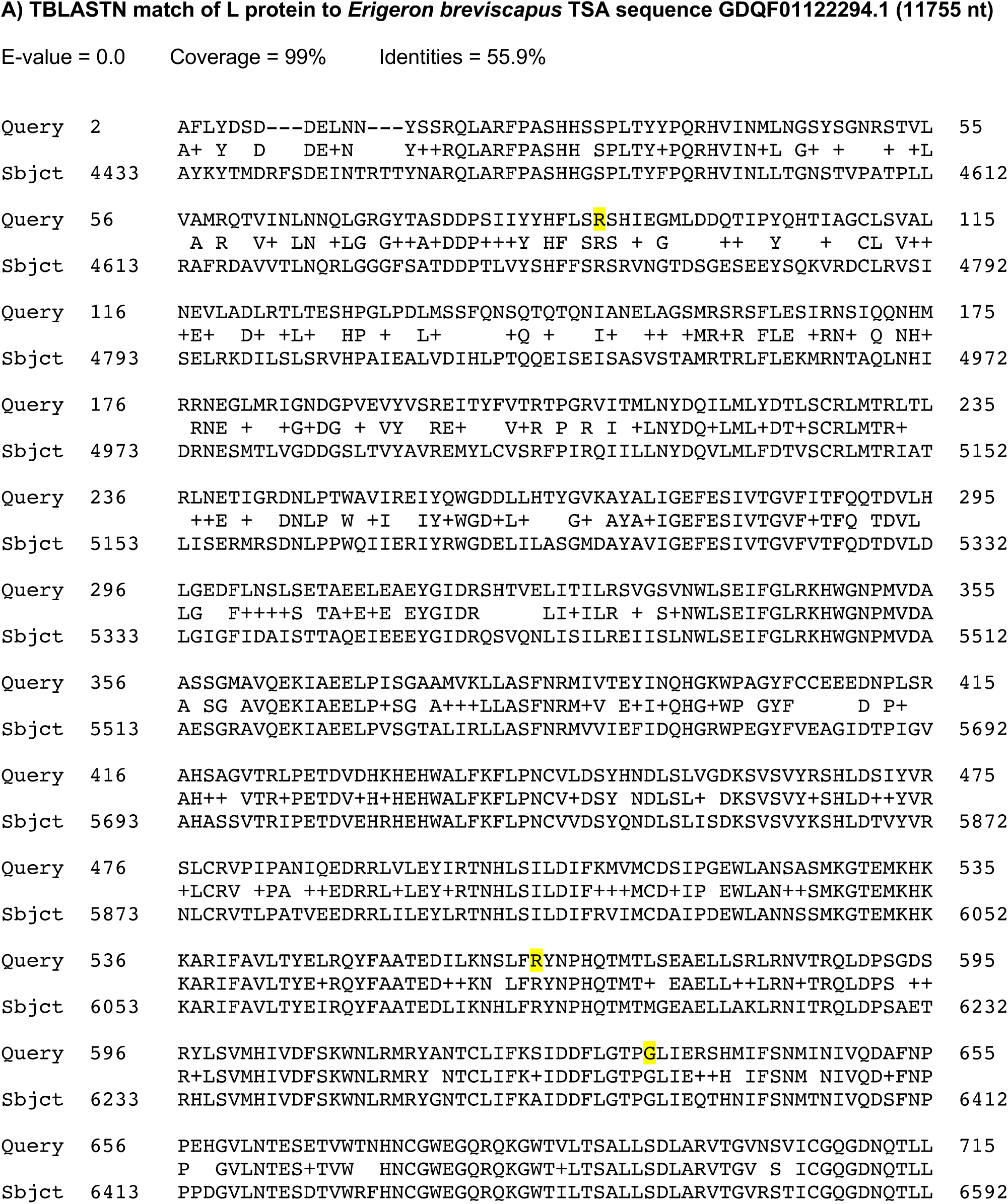
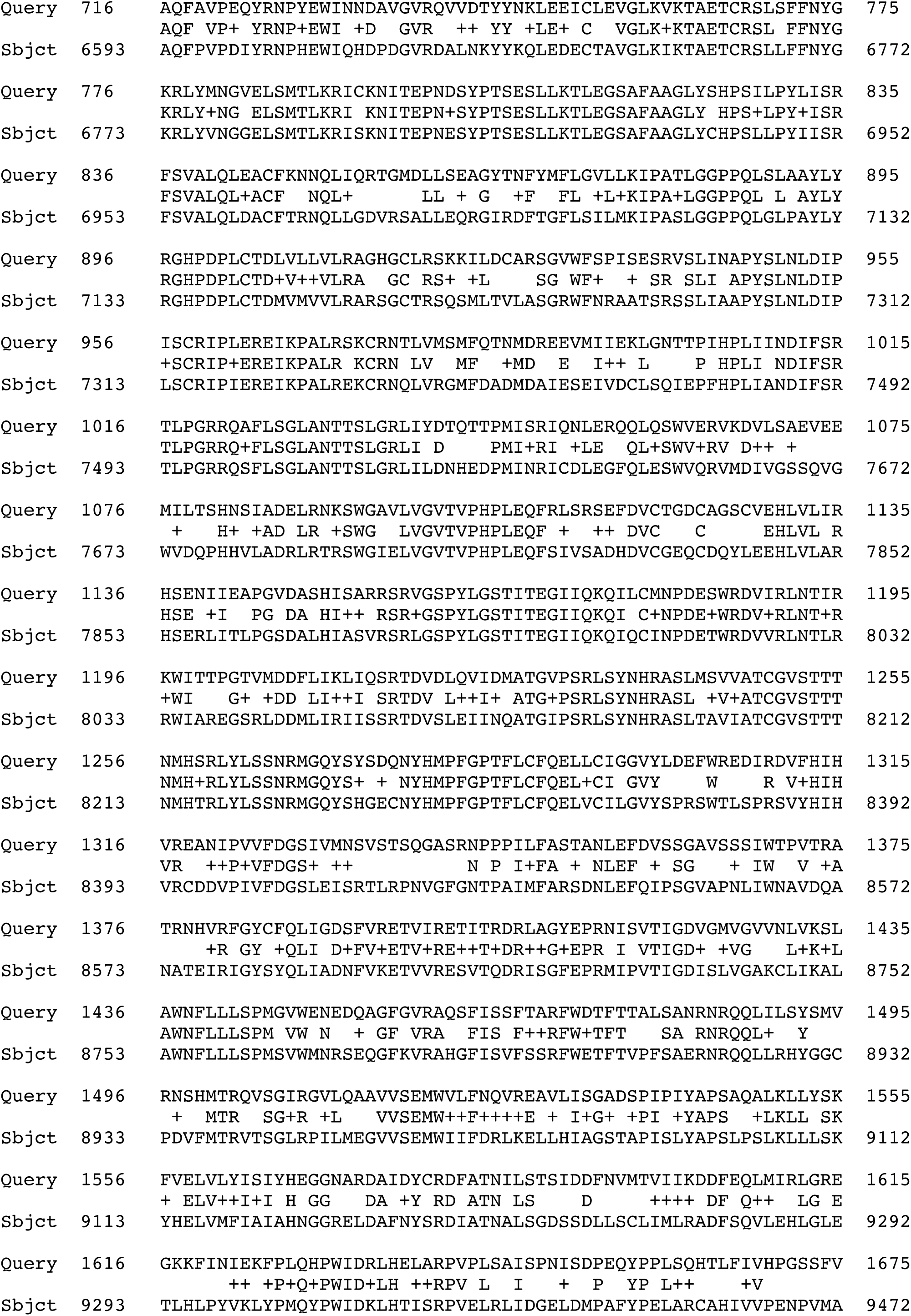
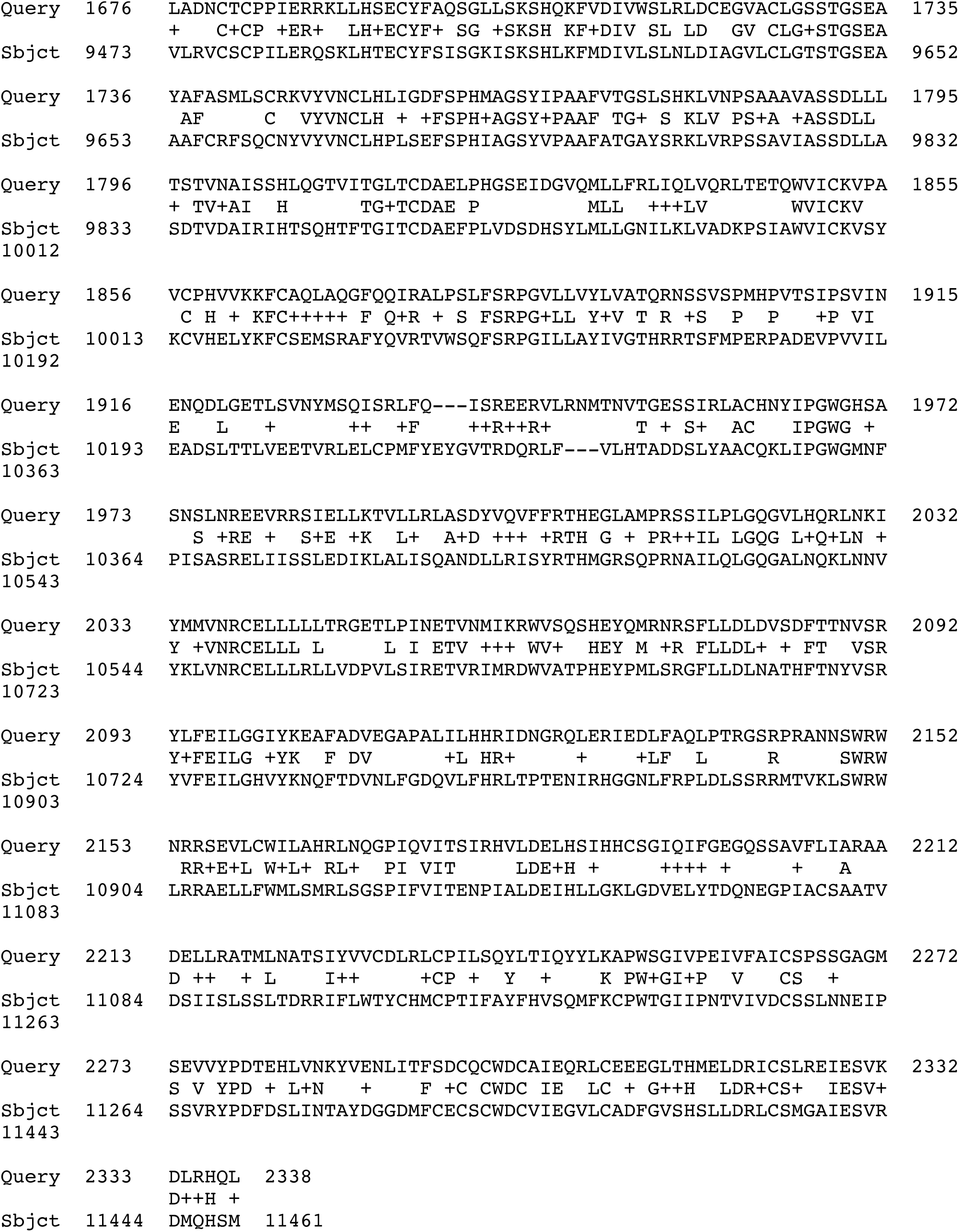
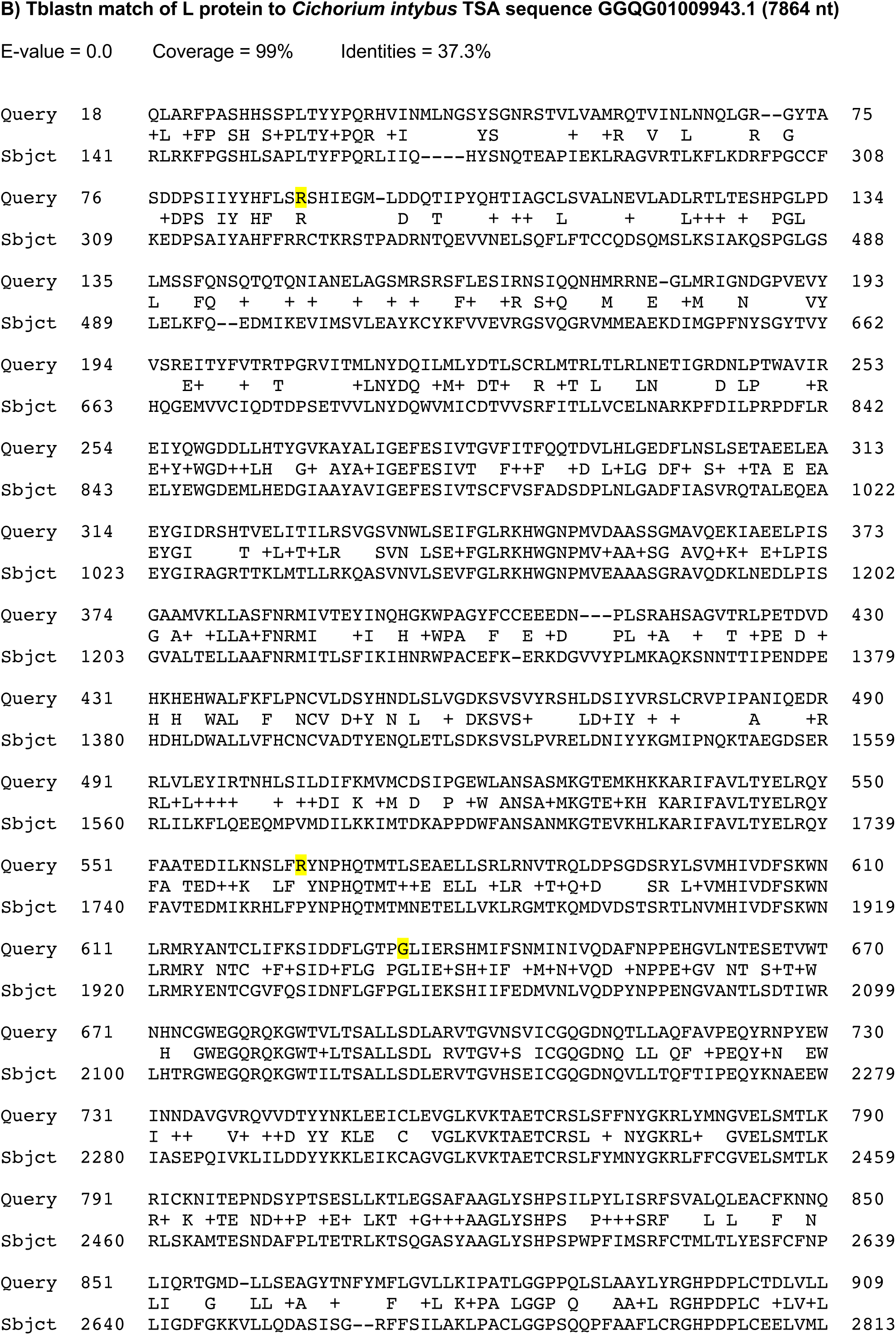
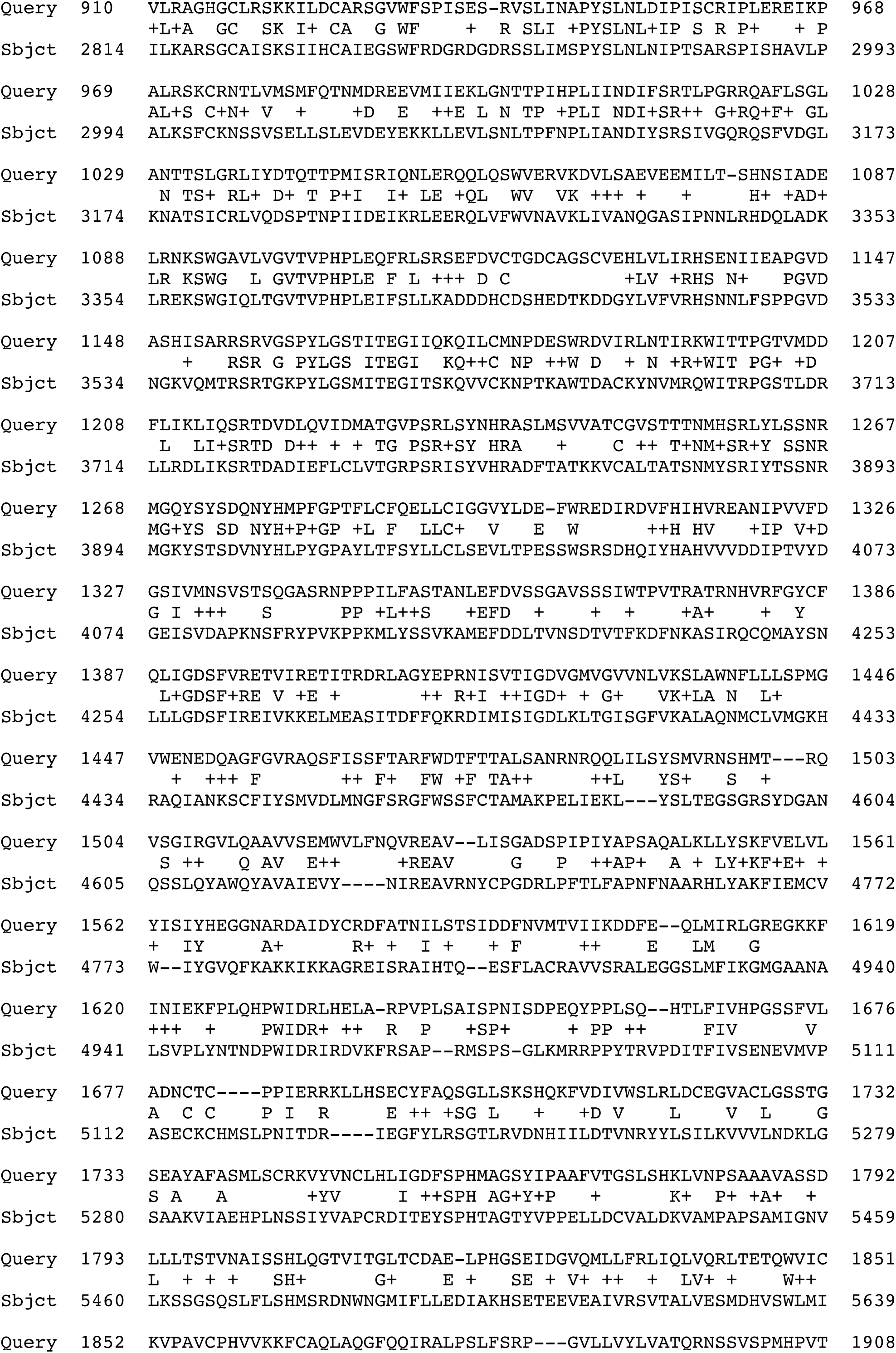
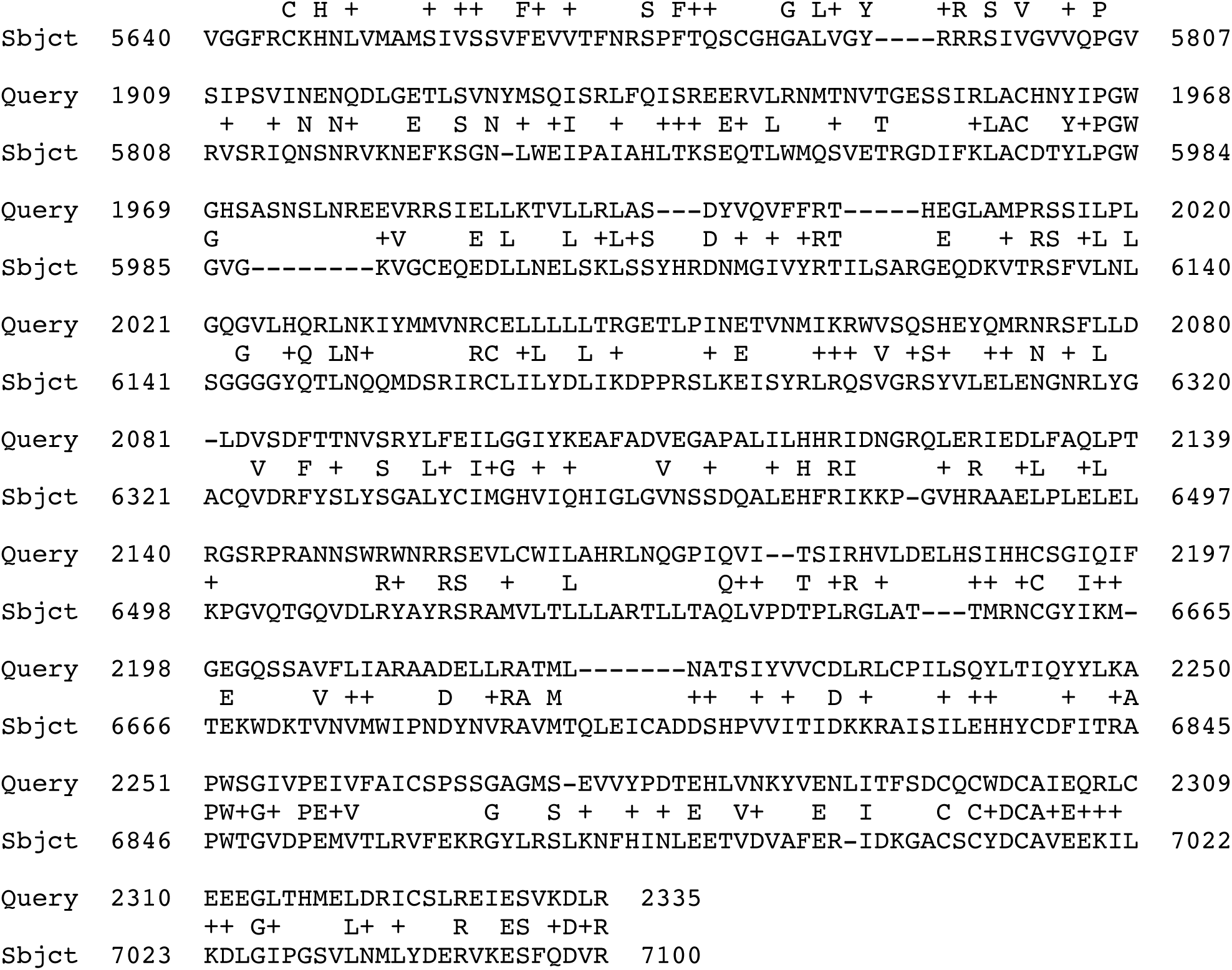
Alignments of the predicted L protein of the GEZL01-derived sequence against related sequences from *Erigeron breviscapus* **(A)** and *Cichorium intybus* **(B)** TSA datasets. Alignments were performed using TBLASTN in 2-sequence mode. The three amino acids in the L protein of the GEZL01-derived sequence that are encoded by a codon spanning one of the intron 1–3 exon-exon junctions are highlighted in yellow. A lack of alignment gaps at these sites and continuity in the alignment elsewhere suggests that introns 1–3 are functionally utilised whereas intron 4 and the alternative introns 1 and 4 are not.

**Supplementary Figure 17.**
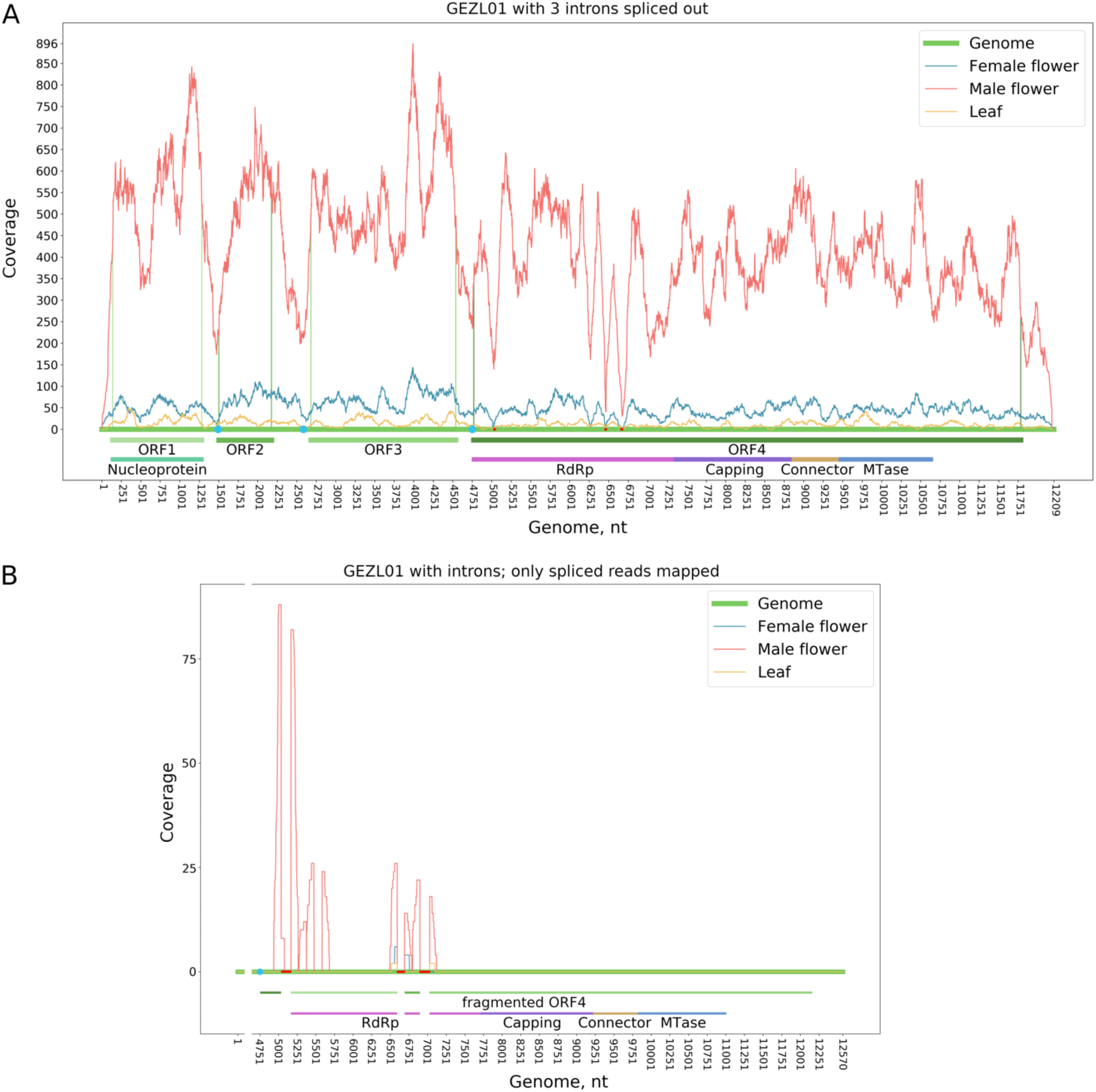
Mapping of Illumina sequencing reads to the GEZL01-derived sequence. **(A)** Genome coverage by reads from three different samples (female flower – blue, male flower – red, leaf – yellow). The ORFs and identified protein domains are indicated below. Light blue dots mark putative transcription stop-start sequences. Introns 1– 3 were removed before read mapping. Mapping was performed with Bowtie2 (Langmead & Salzberg, 2012) (end-to-end, seed length L 32, N 0; only reads with MAPQ > 30 and with length > 80 nt are plotted). Both strands were combined for total coverage. **(B)** Positions of spliced reads mapped to the region encoding the L protein. Reads were mapped to the GEZL01-derived sequence (before removal of any introns) using HISAT2 (Kim et al., 2019) to identify splice sites; spliced reads (only) were plotted if MAPQ > 10 and read length > 70 nt. Introns 1–3 are annotated.

**Supplementary Figure 18.**
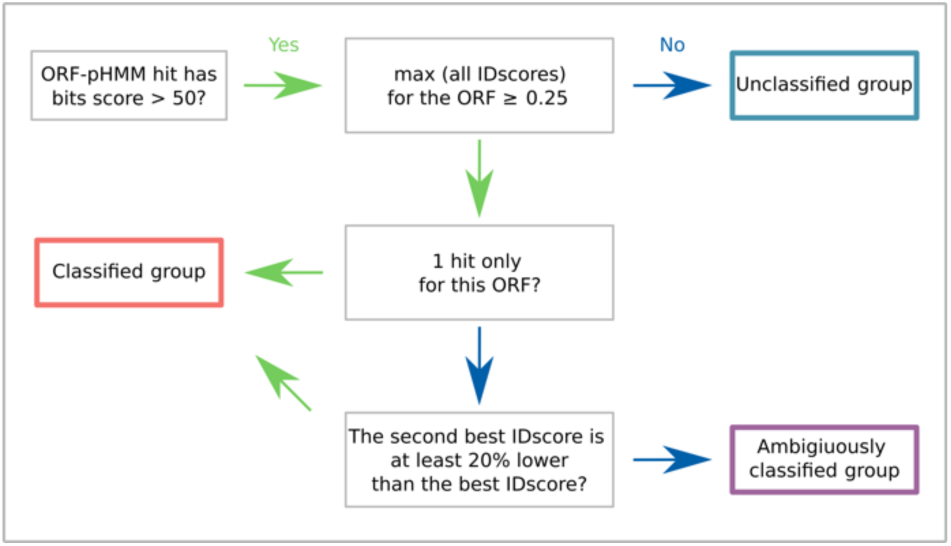
Scheme for grouping HMMsearch (Eddy, 2011) results to pHMMs. Green arrows correspond to “yes”, while blue arrows correspond to “no”.

**Supplementary Table 1.** Numbers of identified RdRp sequences, TSA datasets and unique TSA target species in different taxonomic groups. Only non-identical RdRp core sequences were used (i.e. discarding duplicate 100%-identical RdRp sequences within each classified-pHMM group, including any identical to nr/nt sequences, leaving the longest representative).

|  | RdRps | Datasets | Unique species | RdRps/dataset | RdRps/unique species |
| --- | --- | --- | --- | --- | --- |
| Plants | 1440 | 650 | 275 | 2.22 | 5.24 |
| Invertebrates | 238 | 300 | 172 | 0.79 | 1.38 |
| Arthropods | 1976 | 918 | 592 | 2.15 | 3.34 |
| Vertebrates | 70 | 525 | 202 | 0.133 | 0.347 |
| Fungi | 125 | 77 | 51 | 1.62 | 2.45 |
| Protists | 60 | 155 | 110 | 0.387 | 0.545 |
| Metagenomic | 158 | 24 | – | 6.58 | – |
| Total | 4067 | 2649 | 1413 | 1.54 | 2.88 |

**Supplementary Table 2.** Influenza A virus-like sequences identified in a *Salaria pavo* TSA dataset (GEVH).

| *Salaria pavo*<br>TSA contig |  | BLASTX result (vs NCBI nr protein database);<br>best hit when sorted by 'total score' |  |  |  |
| --- | --- | --- | --- | --- | --- |
| Accession number | Length (nt) | Protein | Accession | Cover | Identity |
| GEVH01318059.1 | 2364 | PB1 | ABB04936.1 | 96% | 99.5% |
| GEVH01036606.1 | 2362 | PB2 | P31345.1 | 96% | 99.9% |
| GEVH01036607.1 | 601 | PB2 | ACN50227.1 | 49% | 98.0% |
| GEVH01006513.1 | 2252 | PA | Q2RF99.1 | 95% | 100% |
| GEVH01127913.1 | 1782 | HA | AFG98995.1 | 95% | 99.8% |
| GEVH01391839.1 | 1046 | M1 | BAA01449.1 | 78% | 94.1% |
| GEVH01289360.1 | 1584 | NP | AAD12236.1 | 94% | 100% |
| GEVH01166396.1 | 1490 | NA | AAB03361.1 | 94% | 99.6% |
| GEVH01333380.1 | 921 | NS1 | ABB04933.1 | 77% | 100% |

**Supplementary Table 3.** Influenza A virus-like sequences identified in a *Nibea albiflora* dataset (GFMR).

| *Nibea albiflora*<br>TSA contig |  | BLASTX result (vs NCBI nr protein database);<br>best hit when sorted by 'total score' |  |  |  |
| --- | --- | --- | --- | --- | --- |
| Accession number | Length (nt) | Protein | Accession | Cover | Identity |
| GFMR01005537.1 | 933 | PB1 | ARX92944.1 | 97% | 99.0% |
| GFMR01048742.1 | 933 | PB1 | AJS16067.1 | 97% | 99.7% |
| GFMR01048741.1 | 430 | PB1 | AFF27591.1 | 62% | 98.9% |
| GFMR01060712.1 | 280 | PB2 | BAM76301.1 | 98% | 98.9% |
| GFMR01010405.1 | 316 | PB2 | QCT26991.1 | 95% | 99.0% |
| GFMR01060713.1 | 490 | PB2 | AAT65240.1 | 63% | 99.0% |
| GFMR01060714.1 | 517 | PB2 | ABC48847.1 | 59% | 97.1% |
| GFMR01004056.1 | 559 | PA | ADM95244.1 | 74% | 93.0% |
| GFMR01045192.1 | 559 | PA | ADM95244.1 | 74% | 93.0% |
| GFMR01002354.1 | 1566 | NP | ABK00128.1 | 95% | 99.6% |
| GFMR01040331.1 | 1566 | NP | AAY87414.1 | 95% | 99.0% |
| GFMR01008042.1 | 996 | M1 | AIX94866.1 | 75% | 98.4% |
| GFMR01055396.1 | 996 | M1 | AAO52906.1 | 75% | 99.2% |
| GFMR01001558.1 | 792 | NS1 | Q809Y0.1 | 85% | 99.1% |
| GFMR01037854.1 | 792 | NS1 | ACA47807.1 | 87% | 96.6% |

**Supplementary Table 4.** Comparison of different influenza A virus strains. Proteins from influenza A virus strain A/New York/392/2004(H3N2) were queried against NCBI influenza A virus reference proteins using BLASTP (Altschul et al., 1990; Camacho et al., 2009).

| Query NCBI accession | Target protein | Lowest/highest non-self cover, % | Lowest/highest non-self identity, % |
| --- | --- | --- | --- |
| YP_308849.1 | PB2 | 100 | 93.9/97.6 |
| YP_308846.1 | PA | 99/100 | 92.2/97.1 |
| YP_308842.1 | NA | 98/100 | 43.2/85.3 |
| YP_308843.1 | NP | 100 | 89.6/94.8 |
| YP_308847.1 | PB1 | 99/100 | 95.4/97.4 |
| YP_308841.1 | M1 | 100 | 90.9/95.2 |
| YP_308840.1 | M2 | 100 | 81.4/95.9 |
| YP_308839.1 | HA | 95/100 | 40.2/46.7 |
| YP_308845.1 | NS1 | 94/100 | 67.0/91.3 |
| YP_308844.1 | NS2 | 100 | 81.0/97.5 |

**Supplementary Table 5.** Comparison of influenza A and influenza B virus reference sequences. Proteins from influenza A virus strain A/New York/392/2004(H3N2) were queried against NCBI influenza B virus reference proteins using BLASTP.

| Query NCBI accession | Target protein | Cover, % | Identity, % |
| --- | --- | --- | --- |
| YP_308849.1 | PB2 | 99 | 37.8 |
| YP_308846.1 | PA | 99 | 36.3 |
| YP_308842.1 | NA | 80 | 31.7 |
| YP_308843.1 | NP | 96 | 37.8 |
| YP_308847.1 | PB1 | 98 | 61.6 |
| YP_308841.1 | M1 | 97 | 30.5 |
| YP_308840.1 | M2 | none | none |
| YP_308839.1 | HA | 95 | 28.5 |
| YP_308845.1 | NS1 | none | none |
| YP_308844.1 | NS2 | none | none |

**Supplementary Table 6.**
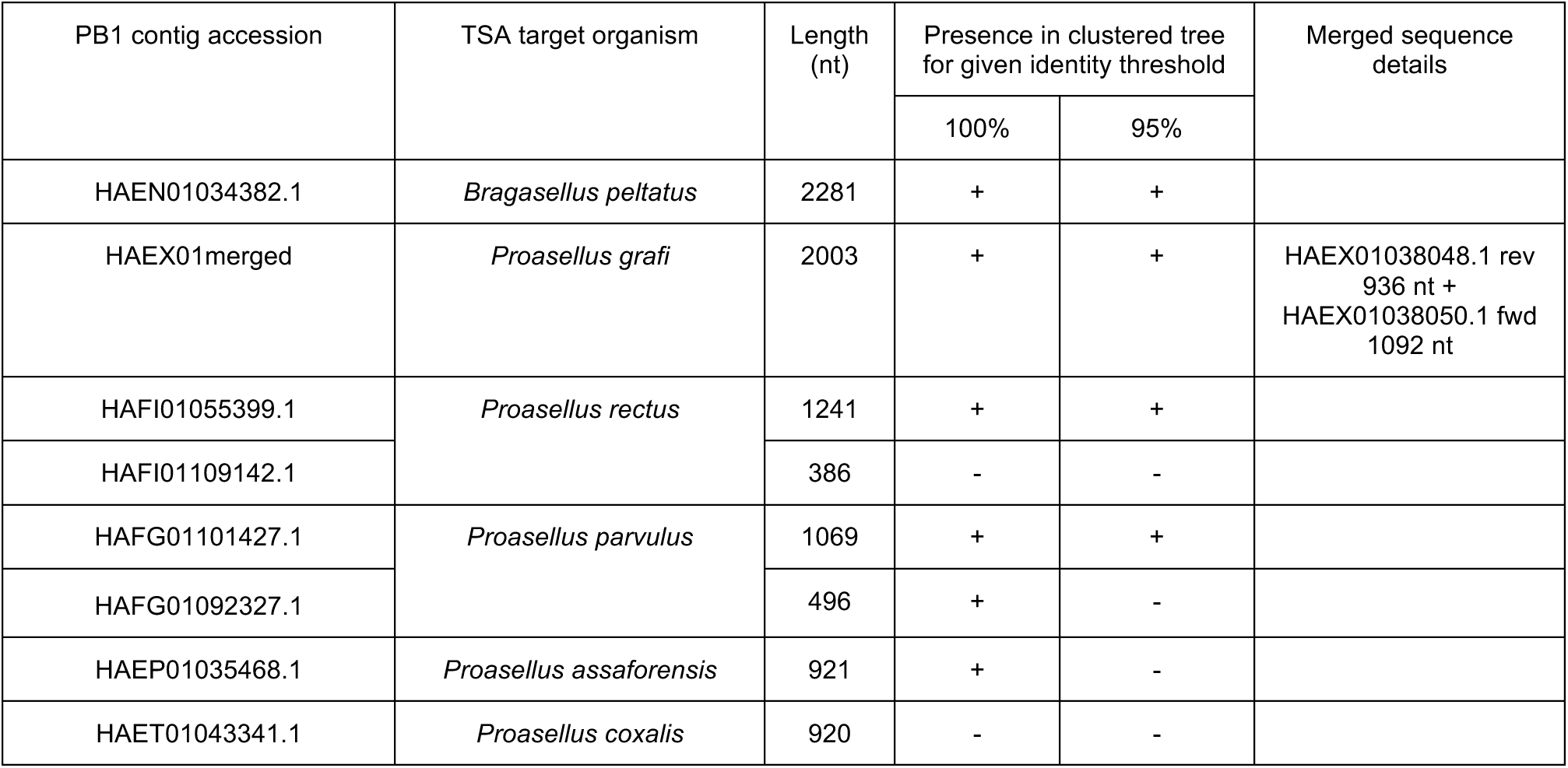
The Asellidae-associated clade of orthomyxovirus-like sequences.

| PB1 contig accession | TSA target organism | Length (nt) | Presence in clustered tree for given identity threshold |  | Merged sequence details |
| --- | --- | --- | --- | --- | --- |
|  |  |  | 100% | 95% |  |
| HAEN01034382.1 | *Bragasellus peltatus* | 2281 | + | + |  |
| HAEX01merged | *Proasellus grafi* | 2003 | + | + | HAEX01038048.1 rev 936 nt + HAEX01038050.1 fwd 1092 nt |
| HAFI01055399.1 | *Proasellus rectus* | 1241 | + | + |  |
| HAFI01109142.1 |  | 386 | - | - |  |
| HAFG01101427.1 | *Proasellus parvulus* | 1069 | + | + |  |
| HAFG01092327.1 |  | 496 | + | - |  |
| HAEP01035468.1 | *Proasellus assaforensis* | 921 | + | - |  |
| HAET01043341.1 | *Proasellus coxalis* | 920 | - | - |  |

**Supplementary Table 7.** BLASTX comparison of Asellidea-associated PB1-encoding sequences with selected *Orthomyxoviridae* NCBI reference PB1 proteins (27 July 2022). id. = identity (%), cov. = coverage (%).

| TSA organism | Length (nt) | PB1 contig accession | Influenza B virus |  | Dhori thogoto-virus |  | Thogoto thogoto-virus |  | Quaranfil quaranja-virus |  | Wellfleet Bay virus |  | Infectious salmon anemia virus |  |
| --- | --- | --- | --- | --- | --- | --- | --- | --- | --- | --- | --- | --- | --- | --- |
|  |  |  | id./cov. | E-value | id./cov. | E-value | id./cov. | E-value | id./cov. | E-value | id./cov. | E-value | id./cov. | E-value |
| *Bragasellus peltatus* | 2281 | HAEN01034382.1 | 28.3 /91 | 1× 10<sup>-53</sup> | 26.3 /91 | 8× 10<sup>-68</sup> | 27.9 /92 | 2× 10<sup>-68</sup> | 22.7 /83 | 2× 10<sup>-28</sup> | 22.1 /83 | 2× 10<sup>-31</sup> | 24.7 /63 | 3× 10<sup>-16</sup> |
| *Proasellus grafi* | 2003 | HAEX01merged* | 29.6 /84 | 9× 10<sup>-40</sup> | 28.0 /91 | 3× 10<sup>-55</sup> | 27.6 /92 | 7× 10<sup>-56</sup> | 24.2 /59 | 5× 10<sup>-16</sup> | 23.4 /79 | 2× 10<sup>-15</sup> | 28.7 /42 | 2× 10<sup>-12</sup> |
| *Proasellus rectus* | 1241 | HAFI01055399.1 | - | - | 27.2 /98 | 5× 10<sup>-33</sup> | 28.3 /97 | 3× 10<sup>-38</sup> | 23.7 /84 | 1× 10<sup>-12</sup> | 24.1 /51 | 1× 10<sup>-10</sup> | - | - |
|  | 386 | HAFI01109142.1 | - | - | - | - | - | - | - | - | - | - | - | - |
| *Proasellus parvulus* | 1069 | HAFG01101427.1 | 29.6 /82 | 1× 10<sup>-30</sup> | 28.3 /86 | 2× 10<sup>-26</sup> | 26.9 /83 | 9× 10<sup>-26</sup> | 23.2 /81 | 9× 10<sup>-13</sup> | 25.3 /80 | 6× 10<sup>-18</sup> | 28.6 /31 | 1× 10<sup>-6</sup> |
|  | 496 | HAFG01092327.1 | - | - | 31.1 /91 | 4× 10<sup>-15</sup> | 29.8 /91 | 1× 10<sup>-15</sup> | - | - | - | - | - | - |
| *Proasellus assaforensis* | 921 | HAEP01035468.1 | - | - | 29.6 /95 | 8× 10<sup>-29</sup> | 29.8 /93 | 5× 10<sup>-30</sup> | 23.4 /94 | 2× 10<sup>-12</sup> | 28.3 /70 | 1× 10<sup>-10</sup> | - | - |
| *Proasellus coxalis* | 920 | HAET01043341.1 | - | - | 29.9 /95 | 8× 10<sup>-29</sup> | 29.8 /93 | 6× 10<sup>-30</sup> | 23.4 /94 | 3× 10<sup>-12</sup> | 24.1 /70 | 2× 10<sup>-10</sup> | - | - |

\* HAEX01038048.1 rev 936 nt + HAEX01038050.1 fwd 1092 nt

**Supplementary Table 8.** Divergent orthomyxovirus-like TSA and nr/nt sequences. All contigs were compared using BLASTX against the entire nr protein NCBI database (Sayers et al., 2022; 17 July 2020) and the best scoring matches are shown. id. = identity (%), cov. = coverage (%).

| Orthomyxovirus-like sequence |  |  |  | BLASTX best nr match |  |  |  |
| --- | --- | --- | --- | --- | --- | --- | --- |
| Data-base | Accession number | Target organism (TSA) or virus name (nr/nt) | Contig length (nt) | Protein accession | Virus name | E-value | id./cov. |
| TSA | GFBM010604515.1 | *Ambystoma mexicanum* | 2551 | QKK82921.1 (PB1) | Neke Harbour virus | $2 \times 10^{-8}$ | 24.6/33 |
| TSA | JP343076.1 | *Lepeophtheirus salmonis* | 557 | MG600033.1 (PB1) | Wenling orthomyxo-like virus 1 | $3 \times 10^{-10}$ | 85.3/12 |
| TSA | GFRZ01merged (GFRZ01104574.1 + GFRZ01104573.1) | *Symsagittifera roscoffensis* | 1823 (1262 + 605) | AVM87616.1 (PB1) | Wenling orthomyxo-like virus 1 | $9 \times 10^{-40}$ | 32.8/73 |
| TSA | GFAT01merged (GFAT01052308.1 + GFAT01014782.1rv) | *Mnemiopsis leidyi* | 2452 (1659 + 1987) | ASM94088.1 (PB1) | Barns Ness dog whelk orthomyxo-like virus 1 | $2 \times 10^{-10}$ | 25.9/54 |
| TSA | GFAT01139990.1 | *Mnemiopsis leidyi* | 723 | AVM87618.1 (PB1) | Yancheng orthomyxo-like virus | $5 \times 10^{-6}$ | 28.7/94 |
| TSA | GFKU01079218.1 | *Apostichopus japonicus* | 2170 | APG77905.1 (PB1) | Hubei orthomyxo-like virus 5 | $5 \times 10^{-9}$ | 20.9/81 |
|  |  |  |  | Best non-self BLASTX nr match |  |  |  |
| nr/nt | MG600033.1 | Wenling orthomyxo-like virus 1 | 2324 | ASM94088.1 (PB1) | Barns Ness dog whelk orthomyxo-like virus 1 | $2 \times 10^{-83}$ | 32.0/90 |
| nr/nt | MF190045.1 | Barns Ness dog whelk orthomyxo-like virus 1 | 2219 | AVM87618.1 (PB1) | Yancheng orthomyxo-like virus | $2 \times 10^{-94}$ | 34.3/86 |
| nr/nt | MG600035.1 | Yancheng orthomyxo-like virus | 2055 | ASM94088.1 (PB1) | Barns Ness dog whelk orthomyxo-like virus 1 | $1 \times 10^{-94}$ | 34.3/92 |
| nr/nt | NC_033319.1 | Changping earthworm virus 2 | 4027 | AVM87618.1 (PB1) | Yancheng orthomyxo-like virus | 0.19 | 24.8/22 |

**Supplementary Table 9.**
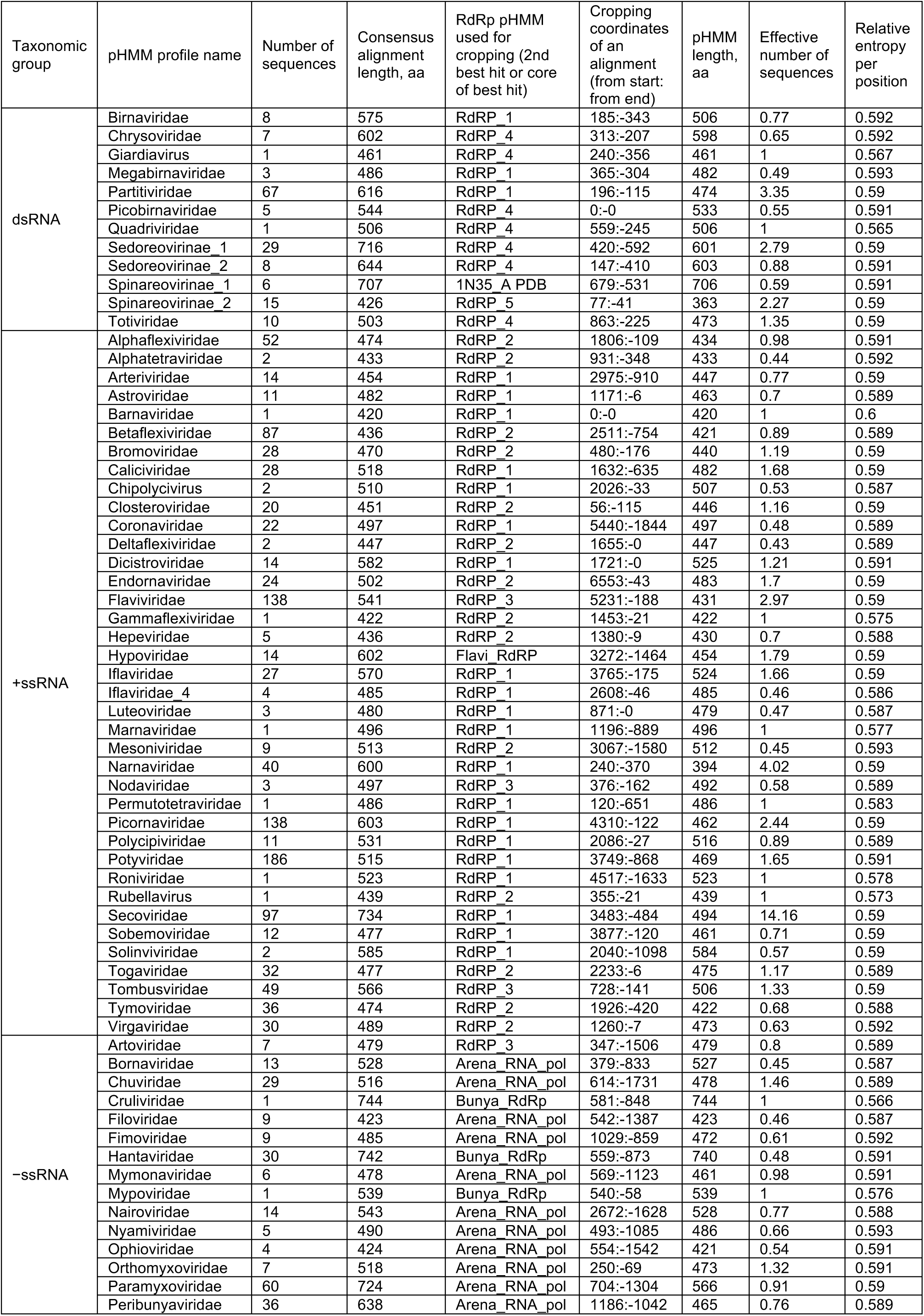
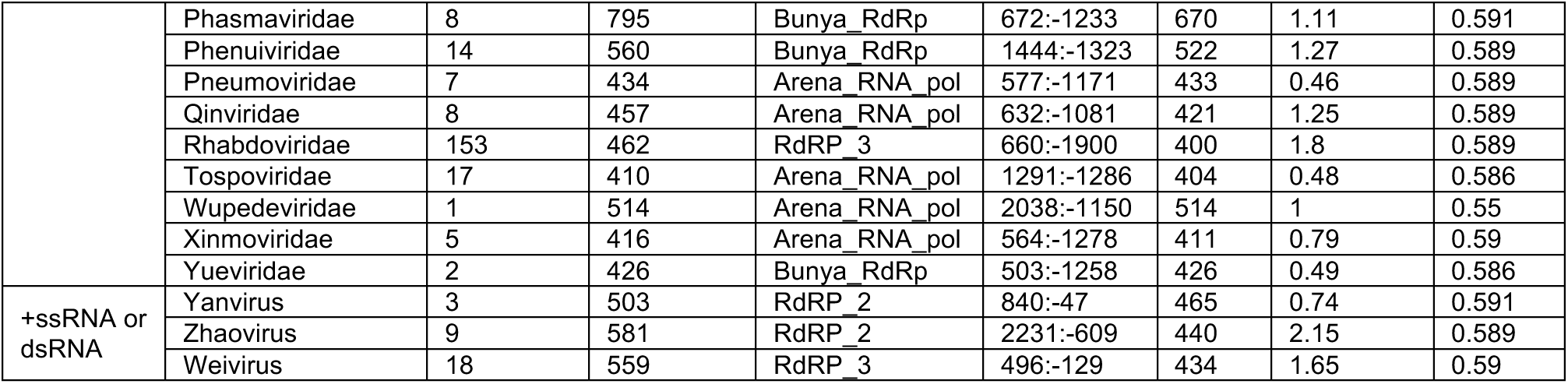
List of RdRp pHMMs with the corresponding number of input sequences, RdRp cropping coordinates, and the HMMbuild (Eddy, 2011) output information.

**Supplementary Table 10.** Search settings used to query the NCBI non-redundant nucleotide database (Sayers et al., 2022).

| Group | Number of sequences | Query used to retrieve sequences | Download date |
| --- | --- | --- | --- |
| dsRNA viruses | 48,989 | txid35325[Organism:exp] AND (viruses[filter] AND ("1000"[SLEN] : "1000000"[SLEN])) | 14 May 2018 |
| –ssRNA viruses excluding influenza A virus | 123,576 | txid35301[Organism:exp] AND (viruses[filter] AND ("1000"[SLEN] : "1000000"[SLEN])) NOT txid11320[Organism:exp] | 14 May 2018 |
| +ssRNA viruses excluding hepacivirus C virus | 91,797 | txid35278[Organism:exp] AND (viruses[filter] AND ("1000"[SLEN] : "1000000"[SLEN])) NOT txid11103[Organism:exp] | 15 May 2018 |
| unassigned ssRNA viruses | 5 | txid674978[Organism:exp] AND (viruses[filter] AND ("1000"[SLEN] : "1000000"[SLEN])) | 14 May 2018 |
| unclassified ssRNA viruses | 59 | txid439490[Organism:exp] AND (viruses[filter] AND ("1000"[SLEN] : "1000000"[SLEN])) | 14 May 2018 |
| unassigned viruses | 0 | txid686617[Organism:exp] | 14 May 2018 |
| unclassified archaeal viruses | 22 | txid451344[Organism:exp] AND (viruses[filter] AND ("1000"[SLEN] : "1000000"[SLEN])) | 14 May 2018 |
| unclassified bacterial viruses | 1833 | txid12333[Organism:exp] AND (viruses[filter] AND ("1000"[SLEN] : "1000000"[SLEN])) | 14 May 2018 |
| unclassified RNA viruses | 3216 | txid1922347[Organism:exp] AND (viruses[filter] AND ("1000"[SLEN] : "1000000"[SLEN])) | 14 May 2018 |
| unclassified virophages | 17 | txid552364[Organism:exp] | 14 May 2018 |
| unclassified viruses | 1362 | txid12429[Organism:exp] AND (viruses[filter] AND ("1000"[SLEN] : "1000000"[SLEN])) | 14 May 2018 |
| environmental samples | 3745 | txid186616[Organism:exp] AND (viruses[filter] AND ("1000"[SLEN] : "1000000"[SLEN])) | 14 May 2018 |

**Supplementary Table 11.**
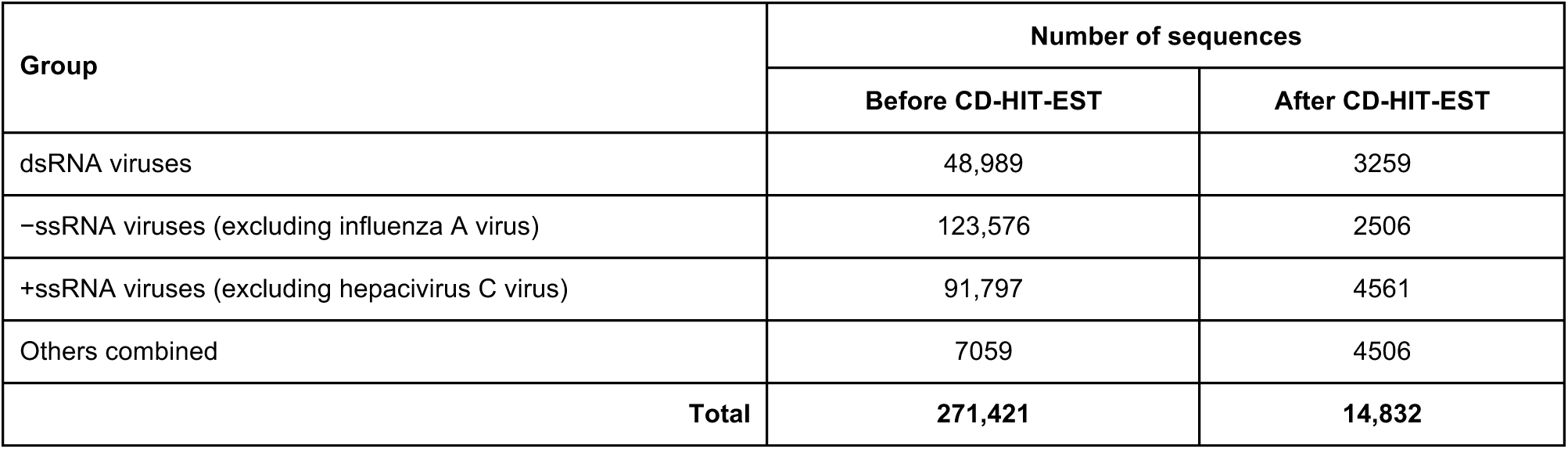
Numbers of sequences before and after applying CD-HIT-EST (Li & Godzik, 2006; Fu et al., 2012) to remove similar sequences.

**Supplementary Table 12.** Numbers of ORFs found for each genetic code table for each sequence dataset.

| Sequences | Number of ORFs |  |  |  |
| --- | --- | --- | --- | --- |
|  | Table 1 | Table 4 | Table 6 | Total |
| Reference | 6,618,649 | 5,842,373 | 5,257,850 | 17,718,872 |
| nr/nt | 1,105,977 | 1,049,250 | 940,742 | 3,095,969 |
| TSA | 4,888,516,023 | 4,484,941,615 | 4,219,750,493 | 13,593,208,131 |
| Totals | 4,896,240,649 | 4,491,833,238 | 4,225,949,085 | 13,614,022,972 |
| Grand total |  |  |  |  |

**Supplementary Dataset 1:** Excel sheet of all 15,044 putative virus-derived RdRp ORFs with associated data.

**Supplementary Dataset 2:** Excel sheet containing the results of our BLAST (Altschul et al., 1990; Camacho et al., 2009) and HHSearch (Steinegger et al., 2019) verification of putative viral ORFs.

**Supplementary Dataset 3:** PhyML (Guindon & Gascuel, 2003; Guindon et al., 2010) phylogenetic trees for the 60 clusters of classified virus-derived RdRp sequences.

**Supplementary Dataset 4:** The 77 viral RdRp pHMMs and their associated sequence alignments.

## Supplementary Methods

### Verification of pHMM matches

In order to verify that sequences identified using the pHMMs were true viral RdRps rather than false positives, and to identify any chimeric sequences, all ORFs identified as encoding a viral RdRp or fragments thereof were checked using a BLAST-based approach. Candidate RdRp-encoding ORF sequences were compared to the NCBI non-redundant (nr) protein database, downloaded 19 Jun 2022, using the DIAMOND v.0.9.14 (Buchfink et al., 2021) implementation of BLASTP (Altschul et al., 1990; Camacho et al., 2009) with the default settings, filtered to keep only hits with an amino acid identity >30% and an alignment length >50 amino acids. Where a more sensitive search was required, sequences were compared to either the full nr database or the subset of this database derived from *Orthornavirae* using the online BLASTP web server (blast.ncbi.nlm.nih.gov; Altschul et al., 1990; Camacho et al., 2009; searches performed 10–17 Aug 2022), using the default word size of six initially and a word size of three where required. Online BLASTP hits were only filtered to have an E-value < 0.05.

BLAST target sequences were classified as known RNA viruses, uncharacterised proteins or non-viral proteins. For the uncharacterised proteins (proteins with names including uncharacterised, unclassified, unnamed, hypothetical or similar), many of which are likely to be RNA viral in origin, we used the DIAMOND settings above to verify similarity to at least one known RNA virus; sequences meeting these criteria are referred to hereafter as uncharactered virus-like proteins (UVPs). RNA viral sequences without an NCBI taxonomic classification were also assigned as *Orthornavirae* in this manner. Sequences were considered to be valid and non-chimeric if they met the following criteria: (i) the best match against any sequence was a significant (E-value <0.05) hit against an *Orthornavirae* sequence or a UVP, (ii) there was at least one significant match against an *Orthornavirae* sequence, and (iii) if the best hit was not an *Orthornavirae* sequence then all hits scoring higher than the best *Orthornavirae* hit were UVPs. Chimeric sequences were those which met these criteria except that not all sequences scoring higher than the best scoring *Orthornavirae* sequence were UVPs. Using this method, all 12,136 ORFs were valid hits, and 25 of these were chimeric.

For verification with an alternative HMM-based approach, HHSearch (part of HHSuite v3.3.0; Steinegger et al., 2019) was used on our set of putative viral ORFs, searching against the full Pfam database (Finn et al., 2014; version 35), with the default settings. Results were then filtered to include only those with a query length >20, probability >20 and p-value <0.05. Where no RdRp was detected, inferred alignments were generated using HHBlits (also part of HHSuite) searching against the uniclust30_2018_08 database (Mirdita et al., 2017) with one iteration. These were then used as queries against the same Pfam database, first with HHBlits, then – if no RdRp was detected – with HHSearch (which is more sensitive). Pfam results were manually classified as RNA viral RdRp, RNA viral non-RdRp, non-RNA-viral, unknown (based on annotation as “domain of unknown function” or similar) or uninformative, a category consisting of helicase and AAA domains which are common in both RNA viruses and cellular proteins (Hickman & Dyda, 2005). Sequences were considered to be valid if either the best match was a viral RdRp or everything scoring higher than the best scoring viral RdRp was a viral non-RdRp, unknown or uninformative.

To compare BLASTP directly with HMMSearch, the 1,784 cropped RdRp sequences used to create the 77 viral pHMMs were used to create a BLASTP (Altschul et al., 1990; Camacho et al., 2009; standalone version 2.12.0) database and this database was searched using the default settings with the 12,136 RdRp-encoding ORF sequences as queries. All of our viral ORFs were detected, using a relaxed E-value cutoff of 0.05. To establish the false positive rate, a reference set of human proteins, the “reviewed, Swiss-Prot” set, which should not be RNA viral in origin, was downloaded from UniprotKB (UniProt Consortium, 2021; downloaded from https://www.uniprot.org on 26 Aug 2022) and compared to the same database with the same settings; here, 824 significant hits were found. HMMSearch was applied to the same set of proteins using the settings described above and our 77 pHMM profiles; here, no results were detected with a p-value (adjusted for database size) meeting our criteria.

## Notes

https://github.com/ingridole/ViralRdRp_pHMMs

https://github.com/ingridole/ViralRdRp_pHMMs_2

### Competing Interest Statement

The authors have declared no competing interest.

